# Psychedelics Align Brain Activity with Context

**DOI:** 10.1101/2025.03.09.642197

**Authors:** Devon Stoliker, Leonardo Novelli, Moein Khajehnejad, Mana Biabani, Matthew D. Greaves, Tamrin Barta, Martin Williams, Sidhant Chopra, Olivier Bazin, Otto Simonsson, Richard Chambers, Frederick Barrett, Gustavo Deco, Katrin H. Preller, Robin Carhart-Harris, Anil Seth, Suresh Sundram, Gary F. Egan, Adeel Razi

## Abstract

Psychedelics can profoundly alter consciousness by reorganising brain connectivity; however, their effects are context-sensitive. To understand how this reorganisation depends on context, we collected and comprehensively analysed the largest psychedelic neuroimaging dataset to date. Sixty-two adults were scanned with functional MRI and EEG during rest and naturalistic stimuli (meditation, music, and movie), before and after ingesting 19 mg of psilocybin (functional MRI ≈80 min post-dose; EEG ≈150 min post-dose). Half the participants ranked the experience among the most meaningful of their lives. Under psilocybin, functional MRI and EEG signals recorded during eyes-closed conditions became similar to those recorded during an eyes-open condition. Global functional connectivity increased in associative regions and decreased in sensory areas. Using machine learning to represent neural activity as low-dimensional trajectories, we found that psilocybin reorganised these into structured, context-sensitive patterns of brain activity that reflected both experimental condition and the quality of subjective experience, revealing an organisation that was missed by time-averaged connectivity measures. Under psilocybin, brain networks that ordinarily segregate internal and external processing coherently integrated and aligned neural dynamics with context. This context-alignment manifested as distinct and cohesive neural trajectories in participants reporting positively felt self- and boundary-dissolving effects, corresponding to the felt experience of being part of the environment, which we refer to as *embeddedness*—the subjective experience of being continuous with, rather than separate from, the surrounding environment. The strength of this context-alignment was associated with next-day mindset change, bridging the neural, experiential, and therapeutic dimensions of the psychedelic state. These findings show that the organisation of brain activity covaries with the experiential coherence of the psychedelic state, and provide a systems-level framework for how context-sensitive brain dynamics link neurobiology to subjective experience and behavioural change.

## Main

The profound effects of psychedelics reshape subjective responses to internal and external sensations and are frequently reported as among the most meaningful experiences in life (Griffiths et al., 2006). These states can manifest sustained therapeutic benefits, including reductions in depression, anxiety, and addiction, alongside an increase in social connectedness and overall well-being (Roseman et al., 2018; Ross et al., 2016; Griffiths et al., 2016; Carhart-Harris et al., 2016; Bogenschutz et al., 2015; Pokorny et al., 2017; Kettner et al., 2021; Lyons and Carhart-Harris, 2018).

The brain constructs perception and selfhood by integrating external sensory inputs with internal models of the environment (Friston, 2010). These interactions between sensory and associative brain regions are enabled by structural and functional connectivity (Sporns et al., 2005). Psychedelics such as psilocybin disrupt these interactions by acting at the serotonin 5-*HT*_2A_ receptor, inducing structural and functional plasticity in preclinical models—rapid mPFC spinogenesis persisting for weeks, mechanistically linked to enduring behavioural effects (Shao et al., 2021, 2025)—that can reshape macro-level connectivity (Ly et al., 2018; Kwan et al., 2022; Burt et al., 2021; Schmidt et al., 2024). Studies that examine effective connectivity further suggest that associative network communication becomes reconfigured (Stoliker et al., 2023, 2024a,b; Preller et al., 2019; Avram et al., 2024). Reflecting this reorganisation, individuals frequently report an intensified sense of immersion, in which the context of space, time, and selfhood feels deconstructed and interconnected (Swanson, 2018; Maclean et al., 2012). While the exact mechanisms linking brain network-level shifts to subjective experience remain unknown, these dynamics have typically been characterised as entropic and desynchronised (Siegel et al., 2024; Carhart-Harris, 2018; Muthukumaraswamy et al., 2013; Herzog et al., 2023). A central mediator of these effects is the default mode network (DMN), which supports the integration of information from diverse associative regions spanning spatial, temporal, and self-referential contexts (Greicius et al., 2003; Raichle et al., 2001; Margulies et al., 2016; Gattuso et al., 2022; Smallwood et al., 2021). Under psilocybin, connectivity patterns that constrain networks relax, permitting novel interactions between sensory and associative regions to emerge (Tagliazucchi et al., 2014; Girn et al., 2026). This reconfiguration alters connectivity patterns that underlie emotion, cognition, and perception (Preller and Vollenweider, 2018). Notably, in healthy adults psilocybin produces a persistent reduction in anterior hippocampus–DMN connectivity, and in rodents, preclinical studies report acute DMN hypoconnectivity (Siegel et al., 2024; Reinwald et al., 2023). Although observed in non-clinical samples, this pattern has been proposed as mechanistically relevant to therapeutic effects. However, despite mounting evidence for these shifts in connectivity, how they give rise to meaningful and therapeutically relevant experiences remains unclear.

Psychedelic effects are also context-sensitive, shaped by mindset and setting (Carhart-Harris et al., 2018) and recognised in clinical guidelines (Johnson et al., 2008). Understanding how context influences therapeutic change, and the underlying brain connectivity that can be harnessed clinically, requires a comprehensive examination of how psychedelics reshape the brain’s functional integration.

Existing studies often rely on small sample sizes, limited imaging modalities, or single-context designs, making it difficult to generalise findings. Structured cognitive tasks imposed during imaging also risk conflating task demands with the psychedelic state itself, rather than capturing how the state unfolds naturalistically. Some recent work has partly addressed these limitations (Carhart-Harris et al., 2016; Timmermann et al., 2023; Mediano et al., 2024), but no study to date has combined large-scale sampling, multimodal imaging, and diverse naturalistic contextual manipulations within a single acute session and computational framework.

We acquired the largest single-site acute-phase psychedelic neuroimaging dataset to date, integrating multimodal fMRI and EEG with controlled contextual manipulations across eyes-open and eyes-closed states in a cohort with no lifetime psychedelic experience (N=62). Experimental conditions of rest, meditation, music, and movie were systematically varied, and extensive behavioural measures were collected (see Fig. 1).

**Fig. 1.**
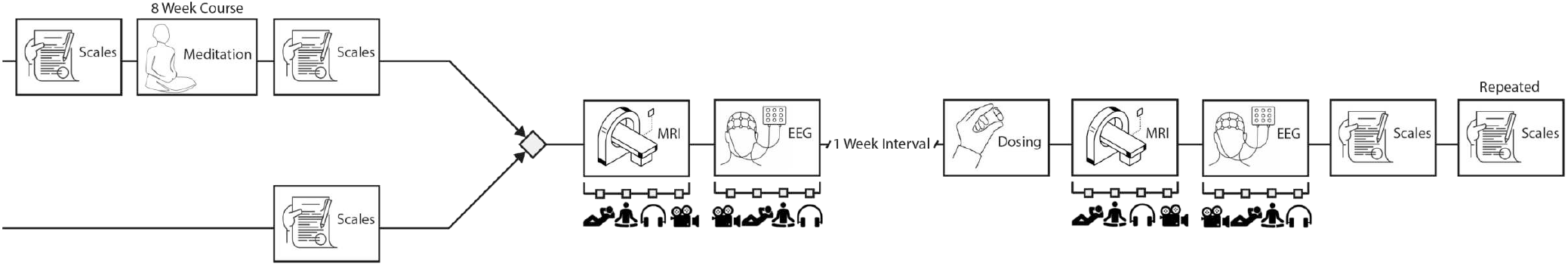
Simplified representation of study design. Participants were assigned to 8 week mindfulness meditation course or immediately enrolled to multimodal (fMRI and EEG) imaging sessions. An open label ‘baseline’ (no-psilocybin) imaging session was conducted approximately one week before imaging under administration of 19 mg psilocybin. MRI and EEG imaging began ≈80 and ≈150 minutes post-dose, respectively. Music, mindfulness, personality and psychedelic measures were collected before and longitudinally after psilocybin. Meditators received additional assessments before and after the mindfulness meditation course. Mindfulness meditation training showed no differences in our connectivity and subjective effect analyses, and was subsequently collapsed for group-level analyses. See Methods for details.

Our analyses show that psilocybin redistributes integration, increasing global functional connectivity in associative regions while reducing it in sensory areas, altering the balance of internally and externally directed processing. Critically, using machine learning (ML), we uncovered context-aligned organisation in brain activity that was missed by conventional analyses. This reorganisation was experientially graded, emerging under positively felt self- and boundary-dissolving experiences, and not during negatively felt states or cognitive impairment.

These results identify a state-dependent neural signature that emerges when network organisation becomes flexibly aligned with context, coinciding with the subjective state we refer to as *embeddedness*. We show this state to be consequential for subsequent change and contextually modifiable. The finding that neural organisation tracks the transformative quality of subjective experience elucidates how the psychedelic state translates into psychological change.

### Integration increased in associative areas and decreased in sensory areas

Psilocybin induced distinct shifts in global integration of functional connections across brain regions. This was quantified using Global Functional Connectivity (GFC), a measure of the average correlation of blood-oxygen-level-dependent (BOLD) signals between each vertex and every other vertex in a high-resolution cortical surface map (Cole et al., 2010). During eyes-closed conditions (rest, meditation, and music), psilocybin reduced the GFC of sensory regions (Fig. 2a), and these reductions formed statistically significant clusters (see Fig. Extended Data 1). Occipital areas showed the strongest overall reductions (percentage changes up to −39% and Cohen’s *d* = −0.55), with midline ventral parietal decreases prominent during rest, meditation and music. In contrast, GFC increased in associative areas during these conditions (percentage changes up to +72%, Cohen’s *d* = 0.53). This evidence indicates that, during eyes-closed conditions, psilocybin suppressed the integration of sensory regions with the broader cortical network and enhanced the integration of associative regions. In contrast, during the eyes-open movie condition, GFC increased in both sensory and associative areas (Fig. 2a), enhancing global synchrony. We confirmed this was not due to head motion (Fig. Extended Data 2). These eyes-closed shifts in GFC are directionally consistent with prior global connectivity studies under psychedelics during eyes-closed rest, which commonly report increases in associative or frontoparietal systems (Tagliazucchi et al., 2016; Madsen et al., 2021) and, in degree centrality analyses, decreases in sensory or visual areas Avram et al. (2025). However, differences in metrics (such as global correlation and functional connectivity density), spatial domains (volumetric vs. surface), and subcortical coverage limit direct comparisons.

**Fig. 2.**
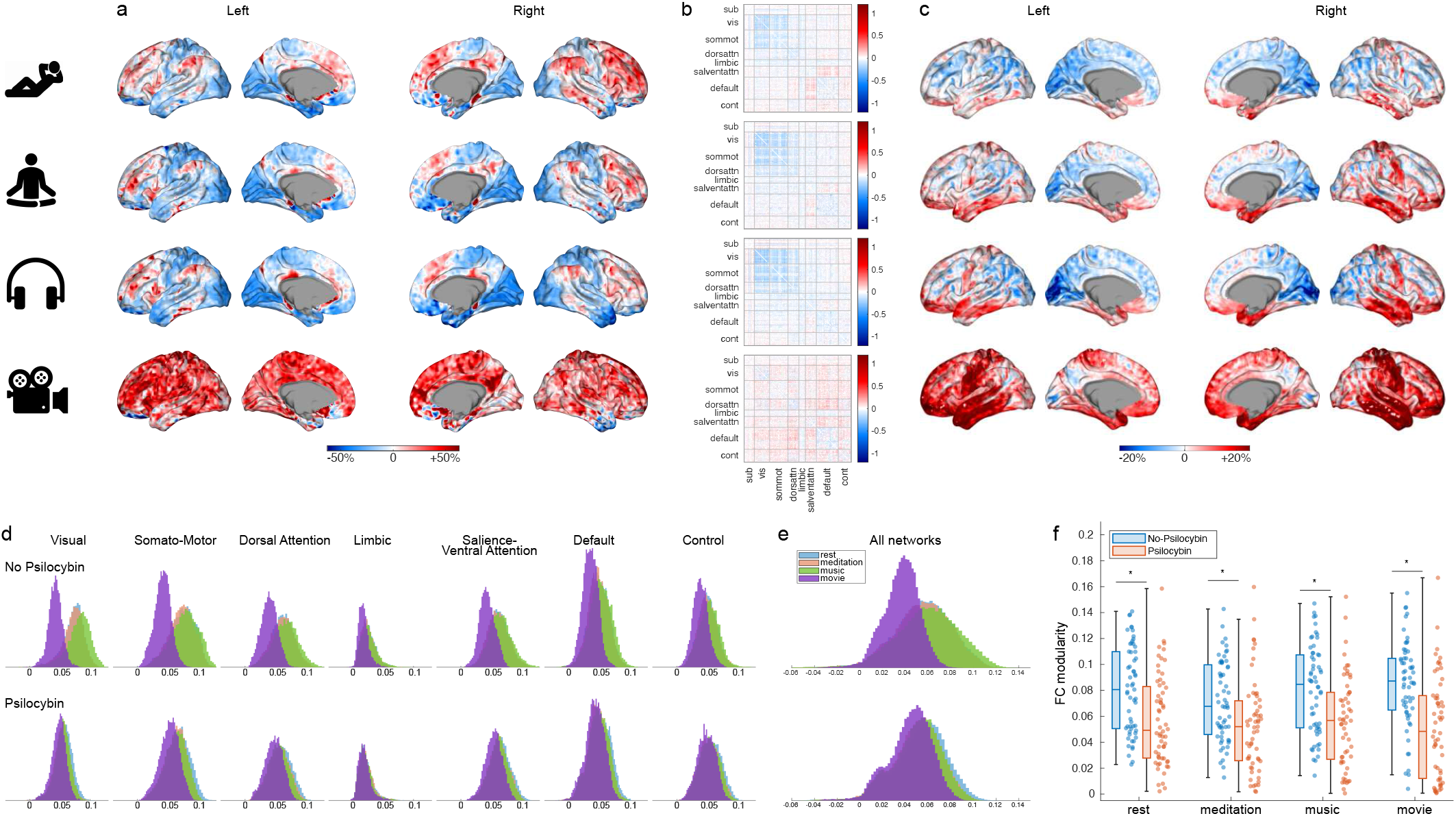
a) Global functional connectivity (GFC) percentage change (psilocybin vs no-psilocybin) averaged across all subjects for each condition (row). The significant clusters after threshold-free cluster enhancement are shown in Fig. Extended Data 1. b) Average functional connectivity difference (psilocybin minus no-psilocybin) across subjects. Psilocybin reduced functional connectivity within networks during eyes-closed conditions. The network names are: subcortex (sub), visual (vis), somato-motor (sommot), dorsal-ventral attention (dorsventattn), limbic, salience-ventral attention (salventattn), default mode (default), and control (cont). A larger version of these FC matrices is provided in Fig. Extended Data 6. c) Standard deviation percentage change (psilocybin minus no-psilocybin) computed for each subject and then averaged across subjects, for each condition. Only cortical vertices where SD changes were not significantly associated with framewise displacement are shown. d) Histograms of GFC values for 7 resting-state networks (columns) and 4 conditions (colour-coded). The gap between eyes-closed and eyes-open conditions at baseline (no-psilocybin; top row) vastly reduces after psilocybin (bottom row). This reduction was most prominent for the visual, sensory-motor, and dorsal-attention networks, but was larger than −66% in all networks and statistically significant (see Fig. Extended Data 5). e) Histograms of GFC values combining all networks. f) Psilocybin reduces functional connectivity modularity across conditions (Cohen’s *d* ≤ −0.6) for all comparisons. Each point corresponds to one participant, the boxes indicate the quartiles, and the whisker length is 1.5 times the interquartile range. Asterisks indicate a significant decrease using the Mann-Whitney U-test (right-tailed, all p-values < 10^−3^).

Further analyses using Z-scoring and global signal regression (GSR) were performed to help reconcile discrepancies reported in prior small-sample studies, where the spatial pattern of GFC changes varied across serotonergic psychedelic datasets and GSR (Preller et al., 2018, 2020; Timmermann et al., 2023). In our psilocybin dataset, analyses produced consistent group-level findings for all eyes-closed conditions, confirming the robustness of the observed effects in our large sample (see Fig. Extended Data 3). However, the movie results changed after GSR and z-scoring, demonstrating that applying these transformations can distort GFC estimates. See Methods for a mathematical rationale for omitting these steps when computing GFC. Despite the consistent group-level findings, substantial variability in GFC changes was observed between participants, indicating individual differences in response to psilocybin (see Fig. Extended Data 4).

### Spatial and context-dependent changes in BOLD signal variance

Mapping the standard deviation (SD) of the BOLD signal within subjects across the cortex revealed that signal variability was spatially reorganised and condition-dependent under psilocybin. Compared to baseline (no-psilocybin), the SD increased during the eyes-open movie condition in the orbitofrontal cortex, occipitotemporal gyri, inferior temporal lobes (particularly anterior regions), and the right hemisphere primary somatosensory cortex (percentage changes up to +19%, Cohen’s *d* = 0.54). In contrast, SD decreased in early visual areas across all eyes-closed conditions, most strongly during music (percentage changes up to −10%, Cohen’s *d* = −0.38). During eyes-open movie, we also detected spatially constrained SD decreases in early visual regions, as well as in the posterior cingulate cortex and precuneus (Fig. 2c), which otherwise showed increased SD as part of widespread cortical increases observed during eyes-closed conditions.

SD increases under psilocybin were partially lateralised to the right hemisphere during eyes-closed rest, meditation and music, and SD decreases in sensory regions were observed in the occipital lobe, particularly during music (Fig. 2c). Finally, the maps of SD change across the brain in all states were broadly consistent with the maps of GFC change (Fig. 2a).

Increased BOLD signal variability under psychedelics is consistent with increased Shannon entropy, as the BOLD-signal distribution is unimodal with finite variance (see Methods). This finding aligns with previous studies that report increased entropy across modalities, including EEG/MEG signal diversity, fMRI sample entropy, and model-derived neuronal firing-rate entropy (Schartner et al., 2017; Lebedev et al., 2016; Timmermann et al., 2019; Herzog et al., 2023).

### Reduced separation between eyes-open and eyes-closed states

Complementing the spatial maps of GFC changes, we examined the overall distribution of GFC values across all cortical vertices, independent of their spatial organisation. Under psilocybin, the histograms of GFC values in eyes-open and eyes-closed states—which were distinct at baseline (no-psilocybin) imaging—largely overlapped (Fig. 2e).

This convergence was consistent and statistically significant across sensory, limbic and associative restingstate networks (RSN) (Fig. 2d and Fig. Extended Data 5) but particularly pronounced in the visual network, where the gap between eyes-closed and eyes-open GFC values decreased by 85% after psilocybin (Cohen’s *d* = −3.23, *p* < 10^−307^).

### Decreased functional modularity

Functional modularity—the degree to which brain activity is organised into distinct, segregated networks— decreased under psilocybin across all conditions (Fig. 2f; rest: Cohen’s *d* = −0.63, *p* = 6.5 ×10^−5^; meditation: *d* = −0.60, *p* = 1.0 ×10^−4^; music: *d* = −0.70, *p* = 8.8 ×10^−5^; movie: *d* = −0.91, *p* = 2.8 × 10^−6^; Mann-Whitney U-test, right-tailed). Comparable decreases in modularity have also been reported in clinical samples, where lower post-treatment modularity was associated with longer-term symptom improvement following psilocybin therapy (Daws et al., 2022). Previous studies have consistently reported disruptions in RSN integration and segregation under psychedelics (Tagliazucchi et al., 2016; Roseman et al., 2014), and have primarily analysed default mode network (DMN) connectivity (Carhart-Harris et al., 2012, 2013; Tolle et al., 2024). Analysis of our larger dataset revealed that psilocybin-induced reorganisation of brain network architecture was primarily attributable to decreased within-network connectivity (Fig. 2b), with significant reductions observed in all networks across eyes-closed conditions (rest, meditation and music), and only in the DMN and dorsal attention network during eyes-open movie (See Supplementary Information).

### Context-aligned neural trajectories emerge with psychedelic experience

Dimensionality reduction methods have been adopted in neuroscience for uncovering meaningful low-dimensional structures in neural data (Cunningham and Yu, 2014). In ML, an *embedding* is a learned mapping from high-dimensional data to a low-dimensional representation, expressed as vector coordinates in an embedding space that preserve latent structure (Cunningham and Yu, 2014; Pillai and Jirsa, 2017; Perl et al., 2025). Here, we used CEBRA to generate low-dimensional embeddings of the preprocessed, frame-by-frame regional BOLD time series and used support vector machine (SVM) classification as a descriptive readout of how well session-specific embeddings distinguished between rest, meditation, music, and movie conditions (Schneider et al., 2023) (see Methods). We then examined the relationship between classification accuracy and subjective experiences assessed via the Mystical Experience Questionnaire (MEQ30)(Maclean et al., 2012; Barrett et al., 2015), administered at the end of the session.

Under psilocybin, structured organisation emerged in the temporal dynamics of brain activity. Notably, stronger subjective effects were associated with tighter clustering of time point embeddings within the same condition, making them more easily separable from embeddings of other conditions (Fig. 3a). This condition-specific clustering was moderated by the timing of subjective effects, as demonstrated by a participant who reported late-onset of substantial effects but minimal subjective experience during the imaging session (Fig. 3b). Their embeddings resembled baseline (no-psilocybin) scans, indicating that the observed neural reorganisation covaried with the occurrence and intensity of subjective effects within the imaging window (Fig. 3b).

**Fig. 3.**
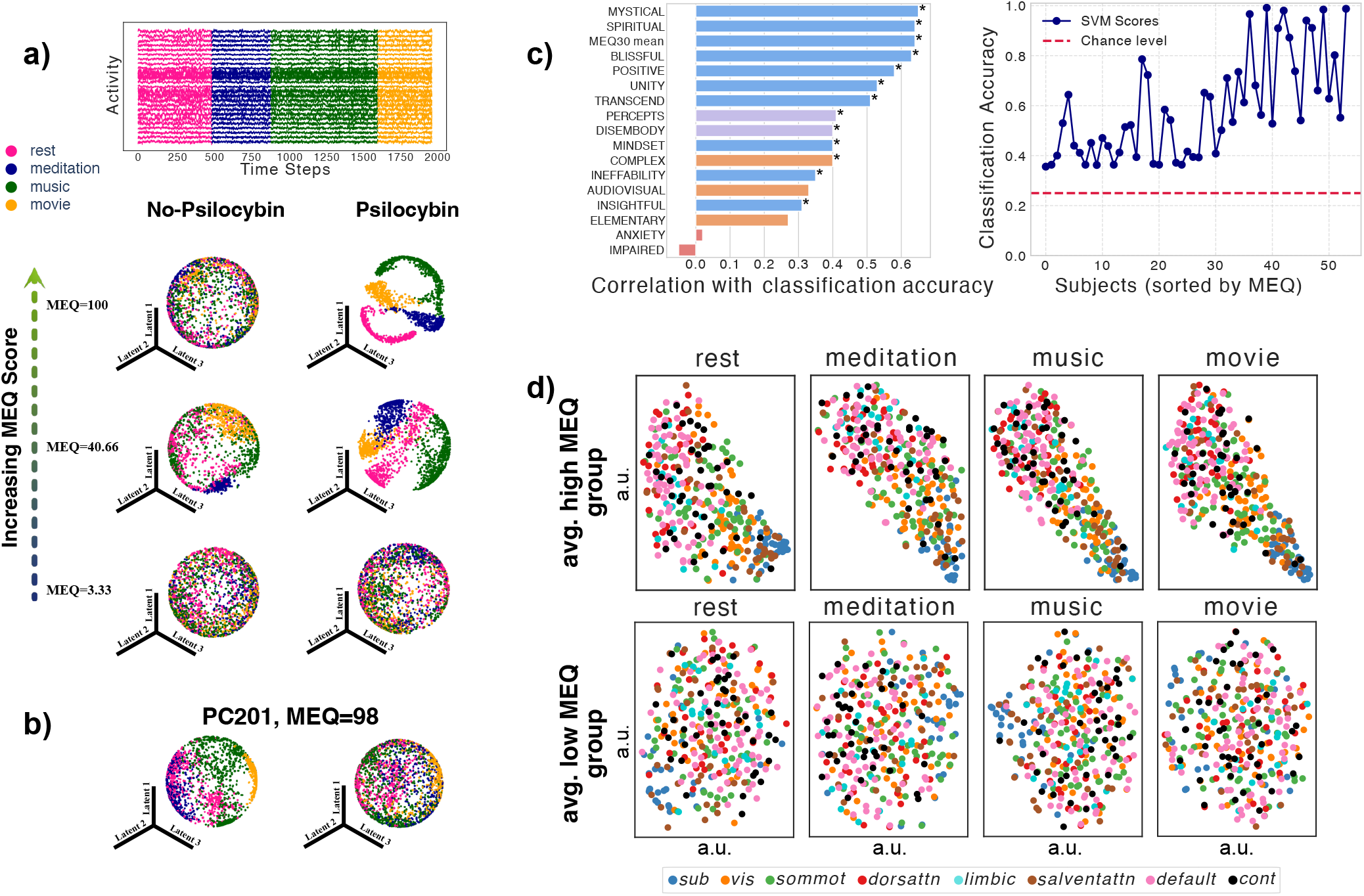
Psilocybin reorganises brain activity along low-dimensional trajectories that scale with subjective effects. a) Using CEBRA—a machine learning method that decodes neural activity over time using contrastive learning—for each participant, we mapped the fMRI time series of all 332 brain parcels at each time point into a 3D space, resulting in a trajectory spanning the entire duration of the four tasks. The pink segment represents rest, while the blue, green, and yellow segments represent the meditation, music, and movie conditions, respectively. b) Network embeddings (CEBRA-derived trajectories) for a participant (PC201), who reported no subjective effect onset during the MRI session on the day of psilocybin administration, showing similarity to their baseline (no-psilocybin) scan. This suggests that neural embedding changes depend on the alignment of subjective states within the imaging time frame. c) For each participant’s CEBRA trajectory, a Support Vector Machine (SVM) was used to classify the correct condition label for each time point. The left panel shows the Pearson correlation between the classification accuracy and acute psilocybin subjective effects including 11 Dimensions of Altered States of Consciousness (11D ASC) and Mystical Experiences Questionnaire (MEQ30) scores. Colours indicate subgroups: Blue = positively felt (euphoric) self- and boundary effects; Orange = sensory-hallucinogenic effects; Red = negative effects; Purple = other (combination) effects. * indicates statistical significance (see Fig. Extended Data 10). The right panel displays the classification accuracy for each subject, compared to the chance level of 0.25, across the four conditions. Subjects are ordered in ascending order based on their MEQ scores. d) A Temporal Attention-enhanced Variational Graph Recurrent Neural Network (TAVRNN) was used to map the 332 ROIs into a 2D space during each condition. The top row shows the average 2D embedding of participants with highest MEQ scores (80–100), while the bottom row shows that of participants with lowest MEQ scores (0–20). Each node corresponds to one ROI and nodes are colour-coded based on their brain network. The axes represent abstract latent dimensions with arbitrary unit (a.u.) learned by the model and do not correspond to physical spatial coordinates.

Further analysis of twelve representative participants, six with high MEQ scores and six with low MEQ scores reinforced these findings, revealing a gradient in brain embeddings that scaled with strength of subjective effects (Fig. Extended Data 7a,b,c,d). This gradient reflected a reorganisation of neural trajectories under psilocybin that was both context-specific and effect-dependent, suggesting that as subjective effects intensified, context increasingly differentiated neural trajectories into distinct, cohesive patterns.

To assess the robustness of these effects, we compared CEBRA embeddings with three common dimensionality reduction methods (PCA, t-SNE, and Isomap) in high-MEQ participants. Despite methodological differences, each approach recovered context-dependent embedding structure (Fig. Extended Data 9), confirming that the observed organisation was not specific to CEBRA.

To quantify the degree of this organisation, we classified the condition label at each time point for each participant with an SVM classifier on the CEBRA-derived trajectories. We examined the correlation between classification accuracy and each of the 11-Dimension Altered States of Consciousness (11-D ASC) (Studerus et al., 2010) and MEQ30 scores, after psilocybin administration. Classification accuracy was most strongly associated with positively felt (valenced) self- and boundary-dissolving effects (blue bars), followed by weaker correlations with sensory-hallucinogenic effects (orange bars), and low or negative correlations with anxiety and impaired control and cognition (red bars) (Fig. 3c, left panel and Fig. Extended Data 10). Colours denote conceptually grouped subscales (see Fig. 4 caption and Methods). We also found a correlation between classification accuracy and mindset change one day after psilocybin (Fig. Extended Data 10; see Supplementary Information for mindset measurement details). Participants with higher MEQ30 mean scores generally exhibited higher classification accuracy, indicating that the context-specific organisation of neural dynamics scales with the intensity of subjective experience (Fig. 3c, right panel). We refer to this structured alignment of neural dynamics with experiential context as *context-alignment*. Notably, these brain–behaviour associations depended on preserving temporal dynamics: per subject functional modularity—a time-averaged measure of network segregation that, despite robust group-level reductions under psilocybin (Fig. 2f; Cohen’s *d* = −0.60 to −0.91)—was not significantly associated with MEQ30 mean or next-day mindset change in any condition (Fig. Extended Data 17).

**Fig. 4.**
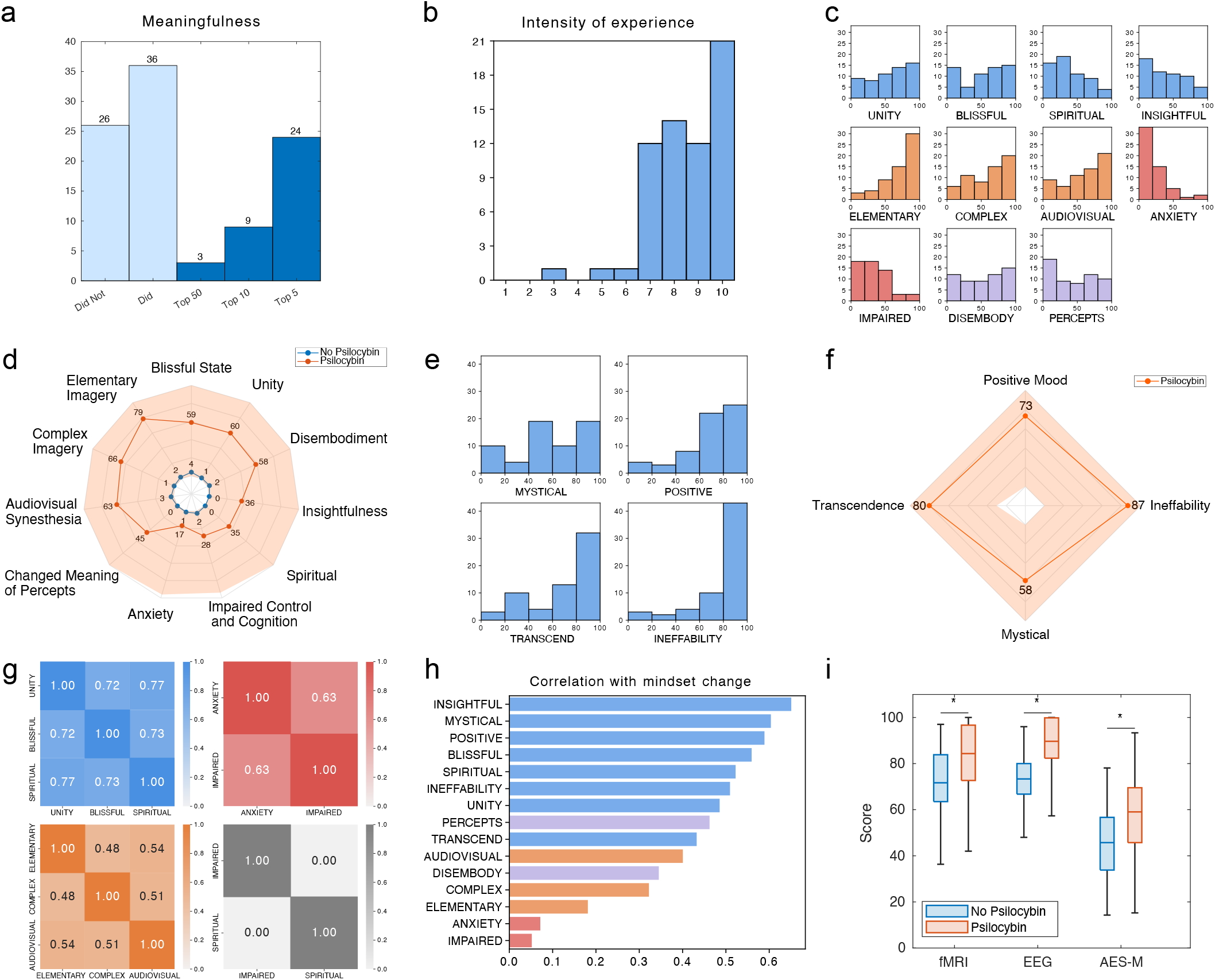
Phenomenological effects reported by participants. a) Distribution of meaningfulness ratings (see Methods). The first two bars represent the number of participants who did (second bar) and did not (first bar) rate their experience as personally or spiritually meaningful. The subsequent three bars indicate the number of participants who rated their experience in the top 50, top 10, and top 5 most meaningful life experiences. b) Histogram of intensity scores from the question “How intense would you rate the psilocybin experience?” reported the day after psilocybin, showing most participants rated the effects as substantially intense. c) Histogram of 11 Dimensions of Altered States of Consciousness (11D ASC) scores across participants. Colours indicate general theoretical distinctions: Blue = positively felt (euphoric) associative effects; Orange = sensory-hallucinogenic effects; Red = negative effects (anxiety, impaired control and cognition); Purple = other (combination) effects. d) Radar plot of 11D ASC showing minimum-maximum range (shaded; psilocybin) and mean scores for dimensions across participants after psilocybin (solid red line) and at baseline (solid blue line). e) Histogram of Mystical Experiences Questionnaire scores across participants, composed of three positively felt (euphoric) associative subdimensions and one valence-neutral (uncategorised) associative subdimension (ineffability). f) Radar plot of MEQ30 showing minimum-maximum range (shaded red) and mean scores (solid red line) for MEQ dimensions across participants. g) Associations between 11D ASC effects reveal strong correlations among positively felt associative effects (blue), sensory-hallucinogenic effects (orange), and negatively felt associative effects (red). No correlation was found between Impaired Control and Cognition and Spiritual Experience. See (Fig. Extended Data 12) for complete matrix of 11D ASC and MEQ within-sample pairwise correlations. h) Correlations between averaged next-day mindset change and acute psilocybin subjective effects (11D ASC and MEQ30), linking each participant’s mindset score to their respective subjective effects scores. This approach captures individual variability, highlighting how specific dimensions, like insightfulness or anxiety, relate to subsequent average mindset scores. See Supplementary Information for mindset score dimensions. i) Box plots of music experience scores (see Methods), showing ratings of music condition during EEG and MRI at baseline and under psilocybin (music experience (fMRI): +16%, Cohen’s *d* = 0.67, *p* = 6.3 × 10^−4^; music experience (EEG): +21%, Cohen’s *d* = 1.08, *p* = 3.2 × 10^−7^). AES-M scores comparing aesthetic music experience at baseline and under psilocybin are also displayed (AES-M: +25%, Cohen’s *d* = 0.75, *p* = 2.4 × 10^−4^). Asterisks indicate a significant increase using the Mann-Whitney U-test (left-tailed). The boxes indicate the quartiles, and the whisker length is 1.5 times the interquartile range.

A network perturbation analysis (see Methods) revealed that this context-aligned organisation is globally distributed but depends most strongly on altered dynamics in Default Mode and Visual networks (~ 6– 8 percentage-point accuracy reduction each when their psilocybin-induced dynamics are replaced with baseline; Fig. Extended Data 8a,b)—the two systems that anchor opposite poles of the principal cortical gradient from internal to external processing Margulies et al. (2016). This dependence was uniform across all four conditions rather than condition-specific (Fig. Extended Data 8c), indicating that the joint alteration of both gradient-endpoint systems sets the conditions for brain activity to align with context. This convergence held across individually trained models (Fig. Extended Data 8d), providing anatomical grounding for how psilocybin reorganises neural trajectories into structured patterns that reflect context. Crucially, if the observed effect merely increased random noise, trajectories would become diffuse and overlapping, degrading classification accuracy. The opposite was observed. Participants reporting stronger effects showed tighter clustering (higher silhouette scores; Fig. Extended Data 7d) and higher accuracy, indicating that the change is structured rather than random.

These findings were independently supported using another ML-based technique called Temporal Attention-enhanced Variational Graph Recurrent Neural Network (TAVRNN) (Khajehnejad et al., 2024), which captured a lower-dimensional representation of individual network connectivity during each condition. These 2D maps represent abstract latent dimensions learned from brain activity patterns, not physical brain coordinates, with each point corresponding to one ROI placed closer to others when their activity evolves similarly over time. TAVRNN captures temporal changes in network structure by modelling sequential snapshots of neuronal connectivity, enabling the identification of key connectivity patterns and shifts in communicability (see Methods for TAVRNN details).

TAVRNN revealed two distinct patterns under psilocybin. Nodes within individual networks exhibited tighter clustering, reflecting more cohesive within-network dynamics. See Supplementary Information and Fig. Extended Data 11 for quantification and method details. Simultaneously, embeddings across all networks showed tighter clustering, indicating a shift toward more integrated brain-wide organisation and increased global cohesion across contexts. This dual pattern of local within-network cohesion alongside global reorganisation scaled with subjective effects, suggesting that the increased within- and between-network cohesion observed under psilocybin depended on the strength of subjective experience and was broadly preserved across contexts (Fig. 3d).

### Phenomenological structure of psychedelic experience

Participants’ reports one day after psilocybin confirmed that their psychedelic experiences during imaging were profoundly altered and meaningful. Half the participants retrospectively ranked it among the most meaningful experiences of their lives (Fig. 4a), and most reported it as substantially intense (> 9*/*10) (Fig. 4b). We also collected semi-structured reports one day after psilocybin, with key excerpts describing the experience as “one of the most peaceful and profound things I could ever experience”, “…meld[ing] with the MRI machine, floor, walls, air”, “los[ing] the plot of who I was, where I was, if I was even here, what was happening” and “los[ing] all sense of self… bec[oming] at one with all my surroundings…as part of a bigger network of things” (see Supplementary Information for a table of qualitative excerpts).

Our large sample enabled a view of within-group subjective reports given a standard 19 mg dose of psilocybin. 11D ASC and MEQ30 scores illustrate substantial group-level intensity across key scales alongside wide individual variability (Fig. 4c,d,e,f). Subscales were grouped to reflect key conceptual distinctions in psychedelic effects: positively felt self- and boundary-dissolving effects (blue; e.g., transcendence, bliss, and unity), sensory-hallucinogenic effects (orange; e.g., visual imagery, synaesthesia), and negative effects (red; e.g., anxiety, impaired control and cognition). Positively felt self- and boundary-dissolving effects such as bliss and unity received higher ratings across our sample than cognitive insights or spiritual experiences. Negative effects, particularly anxiety, were infrequent or rated as low. These subscales were analysed post hoc to assess patterns of covariation (Vollenweider and Kometer, 2010), identifying groups of subscales that showed high intercorrelations in our sample (Fig. 4g). A complete matrix of 11D ASC and MEQ pairwise correlations is provided (see Fig. Extended Data 12).

The Life Attitudes Profile Revised (LAP-R) (Reker, 1992) was used to assess death acceptance and changes in personal meaning before and one month after psilocybin administration. Results showed statistically significant group-level improvements in death acceptance (p = 0.0006, d = 0.49) and personal meaning (p = 0.0004, d = 0.51), alongside individual variability (Fig. Extended Data 13). The Nature Relatedness Scale (Nisbet and Zelenski, 2013) was also administered before and one month after psilocybin to assess changes in participants’ connection to nature, with results indicating both group-level increases (t(55) = −3.37, p = 0.0014, d = 0.26) and individual shifts (Fig. Extended Data 14).

Our sample’s scores were comparable to those observed in higher-dose studies (300–400µg/kg, body-weight adjusted) (e.g., blissful state, experience of unity) (Hirschfeld and Schmidt, 2021). This supports the relevance of the setting conditions employed throughout administration—including space design, minimising interruptions, immersive ethereal music, and encouraging inward focus—in facilitating the states examined during imaging.

### Subjective effects link with next-day mindset change

We examined the relationship between each participant’s post-psilocybin mindset change score (measured one day after administration, see Supplementary Information) and their subjective experience scores from the 11-D ASC scale and the MEQ30. Mindset change, assessed across dimensions such as connection to self, others, and nature, as well as patience, harmony, and inner peace, correlated moderately with associative dimensions of the psychedelic experience. Insightfulness showed the strongest correlation (*r* = 0.65, *p* = 10^−7^), followed by mystical, positive, blissful, and spiritual sub-dimensions (*r* = 0.4 −0.6, *p* < 0.00003). Sensory-hallucinogenic effects were less correlated (*r* = 0.18 −0.4, *p* < 0.05), while negative experiences showed minimal associations (*r* < 0.1, not statistically significant) (Fig. 4h). These findings suggest that positively felt associative dimensions (self- and boundary-dissolving effects) are the primary drivers of psychological shifts, with cognitive insights and transcendent states—such as unitive, blissful, mystical and spiritual experiences—contributing more to mindset change than sensory-hallucinogenic or negative effects (anxiety, impaired control and cognition). The observed relationship between subjective experience and psychological shifts provides empirical context for ongoing debates about the necessity of the psychedelic experience for clinical efficacy (Yaden and Griffiths, 2020; Olson, 2020; Goodwin et al., 2025).

### Context-dependent hippocampal-cortical effective connectivity modulation under psilocybin

Dynamic causal modelling (DCM) (Friston et al., 2003, 2014; Razi and Friston, 2016) estimated directed coupling between the anterior hippocampus (aHip) and core DMN nodes across rest, meditation, music, and eyes-open movie, contrasting psilocybin and baseline (no-psilocybin) conditions (Fig. 5). Group Parametric Empirical Bayes (PEB) contrasts revealed condition-specific changes in effective connectivity. Eyes-closed conditions showed modest changes, while the movie condition exhibited the largest number and magnitude of changes. The total strength of between-region parameter changes (sum of absolute changes with posterior probability (PP) ≥0.99; in Hz) was: rest ≈0.75; meditation ≈0.66; music ≈0.73; movie ≈1.48 (largest). The greater magnitude of directed changes during movie provides mechanistic evidence for context-dependent reorganisation within the Default Mode Network. See Supplementary Information for parameter estimates, credible intervals, and condition-level interpretation.

**Fig. 5.**
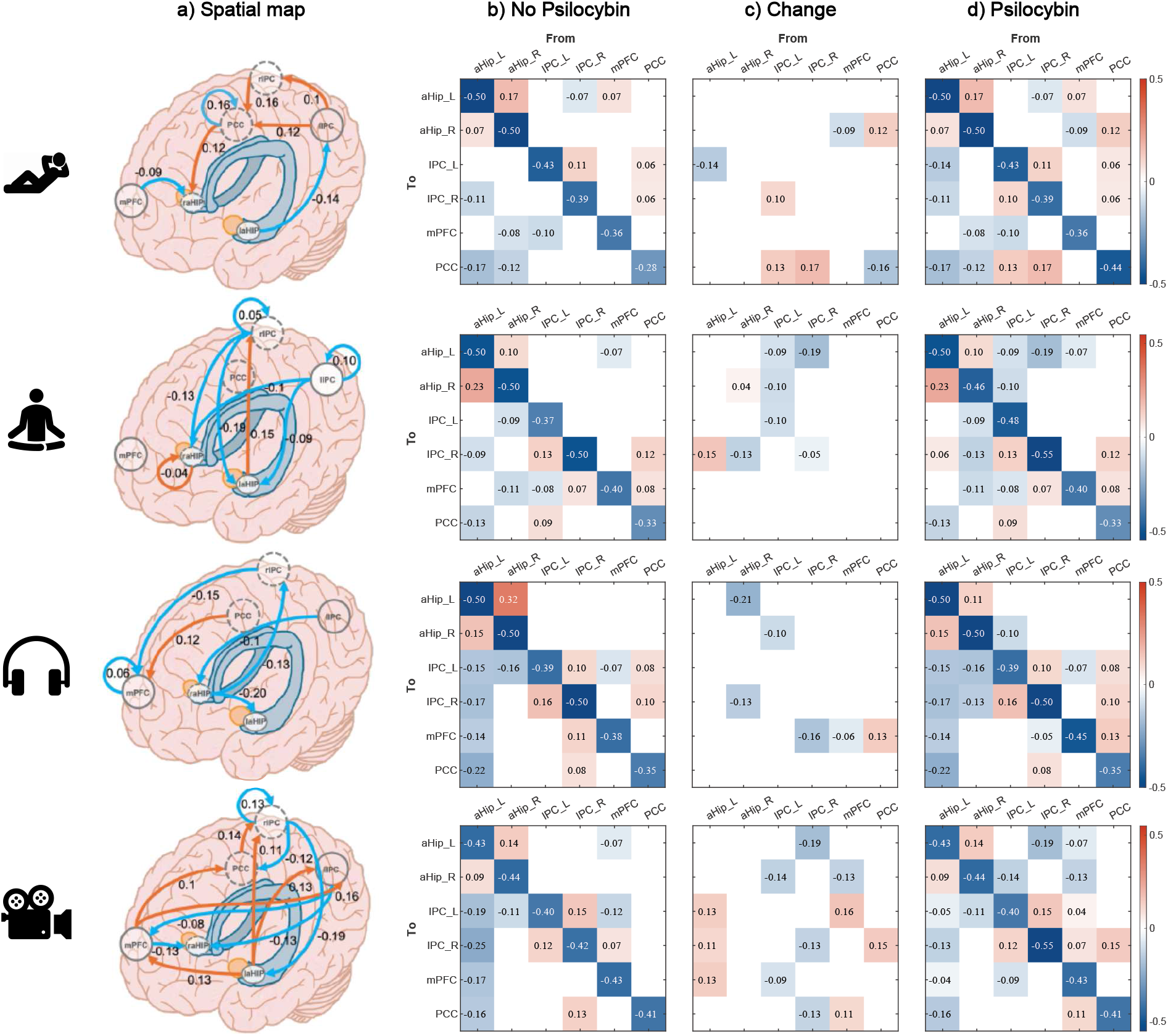
Model of effective connectivity changes from baseline (no-psilocybin) to psilocybin. Brain regions: Left anterior hippocampus (aHip L), right anterior hippocampus (aHip R), left inferior parietal cortex (IPC L), right inferior parietal cortex (IPC R), medial prefrontal cortex (mPFC), and posterior cingulate cortex (PCC). Scan sequences, top to bottom: rest, meditation, music, and movie watching. The model was specified as fully connected, allowing all possible directed interactions between regions. a) Spatial mapping of regions of interest with estimated changes in effective connectivity. Warm colours denote increases, cool colours denote decreases (Hz), based on posterior means averaged across participants using Bayesian model averaging. b) Mean effective connectivity at baseline. For mean effective connectivity, warm colours represent excitation and cool colours represent inhibition. c) Matrix of effective connectivity changes from baseline to psilocybin, corresponding to a). d) Mean effective connectivity under psilocybin. Connection strengths (posterior expectations) are reported in Hz. Self-connections are not log-scaled. See Supplementary Information for tables of posterior expectations and credible intervals, and further details. Displayed connections have a posterior probability > 0.99, indicating very strong evidence.

### Impact of mindfulness meditation training and music on psilocybin effects

Our study included an 8-week mindfulness meditation training program for half of the participants (Fig. 1). By assigning participants rather than allowing self-selection, we aimed to evaluate the clinical utility of structured pre-psilocybin meditation while addressing selection bias common in meditation research. Engagement was consistent across the cohort, with participants attending at least 6 of 8 sessions and averaging 85 minutes of independent practice per week (see Methods for program details). However, no statistically significant differences were observed in acute psilocybin effects (11-D ASC, MEQ30) or next-day mindset between meditators and non-meditators. We also observed no statistically significant changes in connectivity metrics under psilocybin (e.g., functional modularity; Fig. Extended Data 15).

Previous studies reporting synergistic effects between meditation and psilocybin have typically involved experienced practitioners engaged in intensive retreat-based practice (Smigielski et al., 2019; Singer et al., 2024). In contrast, our study participants completed a standardised mindfulness-based cognitive therapy program that was not designed to integrate with the psychedelic session. Despite including a guided meditation scan, no between-group differences emerged, suggesting that synergy may require longer-term meditative engagement, self-directed commitment, or a program explicitly tailored to support the psychedelic experience.

Music, a contextual element known to shape subjective experiences under psychedelics (Kaelen et al., 2018), played a central role in the study design. During MRI and EEG sequences, participants listened to ethereal music selected to enhance emotional depth and psychedelic effects without conflicting with the scanner’s acoustic environment. Participants rated the music as significantly more meaningful and emotionally resonant under psilocybin compared to non-psilocybin conditions (+16% and +21%, Cohen’s *d* = 0.67 and 1.08, *p* = 6.3 ×10^−4^ and 3.2 ×10^−7^; Mann-Whitney U-test, left-tailed; Fig. 4i). Moreover, TAVRNN embeddings showed the greatest global network organisation during music (Fig. 3d). The enhanced subjective experience of music presented throughout the administration day was independently confirmed by the Aesthetic Experiences in Music scale (AES-M; +25%, Cohen’s *d* = 0.75, *p* = 2.4 ×10^−4^), collected at the end of the day (Fig. 4i, see Methods for details).

### Context-sensitive neural integration under EEG

Context-sensitive psilocybin-induced neural dynamics observed using MRI were confirmed using 64-channel wet EEG, both in the power spectrum and in the signal complexity. Our findings extend preclinical and human observations of desynchronised local neural activity under psilocybin (Golden and Chadderton, 2022; Muthukumaraswamy et al., 2013) by demonstrating that these neural alterations are modulated by sensory context. We found that psilocybin expanded the power spectrum, with decreases in theta, alpha, and, to a lesser extent, beta power, accompanied by modest gamma increases during eyes-closed conditions, concentrated in Music, a contextual element known to shape subjective experiences under psychedelics (Kaelen et al., 2018), played a central role in the study design. During MRI and EEG sequences, participants listened to ethereal music selected to enhance emotional depth and psychedelic effects without conflicting with the scanner’s acoustic environment. Participants rated the music as significantly more meaningful and emotionally resonant under psilocybin compared to non-psilocybin conditions (+16% and +21%, Cohen’s *d* = 0.67 and 1.08, *p* = 6.3 ×10^−4^ and 3.2 × 10^−7^; Mann-Whitney U-test, left-tailed; Fig. 4i). Moreover, TAVRNN embeddings showed the greatest global network organisation during music (Fig. 3d). The enhanced subjective experience of music presented throughout the administration day was independently confirmed by the Aesthetic Experiences in Music scale (AES-M; +25%, Cohen’s *d* = 0.75, *p* = 2.4 ×10^−4^), collected at the end of the day (Fig. 4i, see Methods for details).

### Context-sensitive neural integration under EEG

Context-sensitive psilocybin-induced neural dynamics observed using MRI were confirmed using 64-channel wet EEG, both in the power spectrum and in the signal complexity. Our findings extend preclinical and human observations of desynchronised local neural activity under psilocybin (Golden and Chadderton, 2022; Muthukumaraswamy et al., 2013) by demonstrating that these neural alterations are modulated by sensory context. We found that psilocybin expanded the power spectrum, with decreases in theta, alpha, and, to a lesser extent, beta power, accompanied by modest gamma increases during eyes-closed conditions, concentrated in frontal regions. These became attenuated during movie, with increases localised to early visual areas, indicating condition-dependent topography (Fig. 6a). This attenuation in power and complexity during movie, relative to eyes-closed tasks, is consistent with reduced network-level disruption during externally directed attention, as previously shown using fMRI functional connectivity (Siegel et al., 2024). Meditation and music recordings aligned in EEG, showing similar power profiles across frequencies at baseline and under psilocybin, converging despite distinct stimulus properties, while rest modestly differed (Fig. 6b). Psilocybin also reduced the difference between eyes-open and eyes-closed alpha power by 48% (Cohen’s *d* = −0.79, Mann-Whitney U-test right-tailed p-value < 10^−19^), suggesting an integration of internally- and externally-focused neural processing (Fig. 6b,c and Fig. Extended Data 16).

**Fig. 6.**
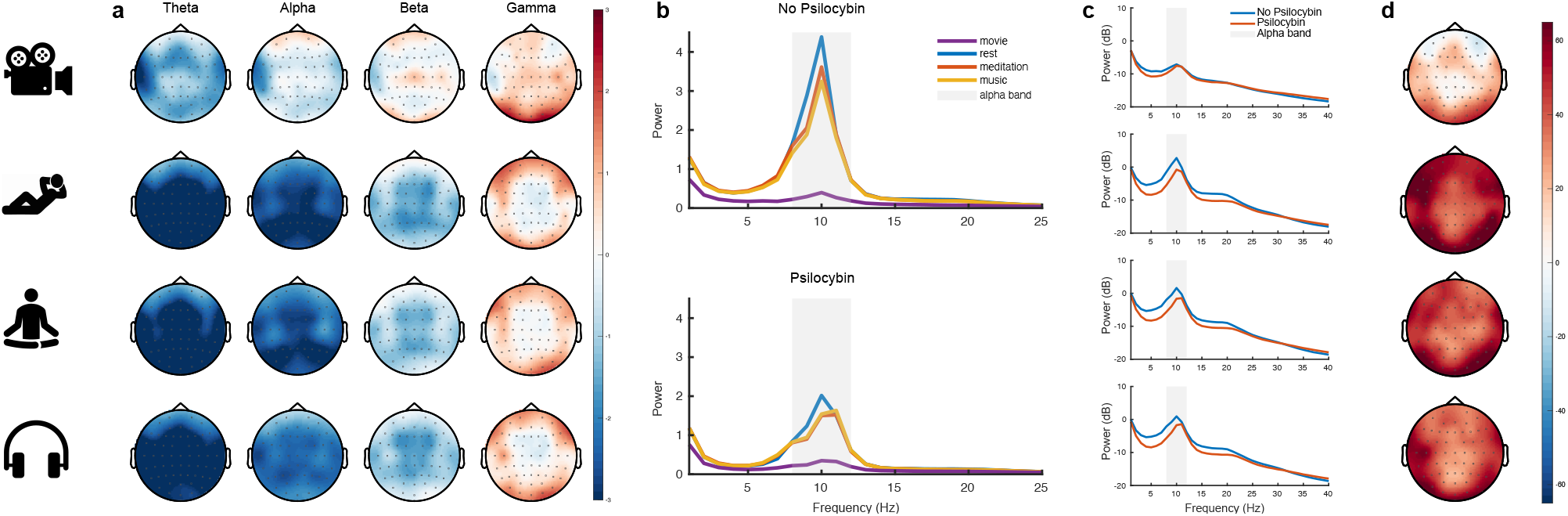
EEG spectral and complexity changes showed reduced eyes-open vs eyes-closed separation under psilocybin. a) Spatial difference in power (psilocybin minus baseline; dB units) for each frequency band. b) Group-averaged power spectra for each condition at baseline and psilocybin. Under psilocybin, all eyes-closed conditions (rest, meditation, and music) showed broad-band reductions in theta/alpha/beta power (meditation and music overlap; rest modestly differed), whereas the eyes-open movie spectrum was relatively unchanged. c) Group-averaged power spectra plotted separately for each condition, in order to compare psilocybin vs no-psilocybin. d) Lempel-Ziv complexity (psilocybin minus baseline) showed similar values across eyes-closed conditions and reduced complexity during eyes-open movie.

Signal diversity, quantified via Lempel-Ziv complexity, was highest in eyes-closed conditions, consistent with recent findings (Mediano et al., 2024), suggesting that internally generated perceptions are associated with increased neural complexity (Fig. 6d). Alpha-band activity (8–12 Hz), typically associated with suppressing visual input during eyes-closed conditions (Kometer et al., 2013; Toscani et al., 2010), decreased under psilocybin, particularly in visual regions. This reduced sensory filtering, combined with changes in MRI functional connectivity patterns, suggests diminished distinctions between eyes-open and eyes-closed conditions.

Comparable decreases in alpha power and increases in signal diversity have also been observed under intravenous administration of the serotonergic psychedelic N,N-dimethyltryptamine (DMT) Timmermann et al. (2023), suggesting these may be conserved markers of serotonergic psychedelic action, despite differences in compound, route of administration, timing, and EEG methodology. The convergence of the MRI and EEG findings confirms that this effect cannot be explained solely by vascular changes caused by psychedelics (Carhart-Harris et al., 2012). Together, these results establish context-alignment as a property of brain dynamics under psilocybin that is evident across imaging modalities, timescales, and analytical approaches.

### Psilocybin transforms embeddedness in brain activity and perception

Machine-learning (ML) embeddings revealed an association between positively felt self- and boundary-dissolving subjective effects and the reorganisation of brain networks into context-aligned clusters (Fig. 3b,d). The relative activation of these networks regulates the balance between internally (e.g., DMN) and externally directed processing, a dynamic that shapes perception and cognition (Fox et al., 2005; Raichle et al., 2001). Their increased integration under psilocybin suggests a shift toward more flexible connectivity and altered functional interactions (Fig. 3d).

The correlation between self- and boundary-dissolving effects and classification accuracy based on ML embeddings (Fig. 3c) indicates that these embeddings capture a connectivity state in which internal and external processes become less distinct. This is consistent with participants’ reports of feeling integrally part of a broader physical and psychological relational context. We refer to this state as *embeddedness*, the phenomenological correlate of context-alignment. Unlike connectedness, which implies links between separate entities, embeddedness is the subjective experience of being continuous with, rather than separate from, the environment, which emerges when brain dynamics become aligned with context in proportion to the depth of self- and boundary-dissolving effects. Psychedelics appear to facilitate this state—a perceptual and cognitive realisation of being part of a unified whole—by reorganising brain network dynamics to increase integration within and between networks (Fig. 3d). The context-sensitivity of this reorganisation— its dependence on what the participant is doing and experiencing during psilocybin administration—suggests that the underlying synaptic changes may themselves be shaped by ongoing neural activity.

While embeddedness refers to an experiential state, CEBRA and TAVRNN are machine learning methods that learn embedding spaces from imaging data. Although the experiential construct and the computational representations are conceptually distinct, the context-organised clustering of low-dimensional coordinate vectors within the learned embedding spaces covaried with the intensity of subjective effects (Fig. 3b,d; Fig. Extended Data 7a,b), indicating an association with experiential embeddedness.

CEBRA embeddings showed that the highest classification accuracy covaried with the occurrence of intense, positively felt, self- and boundary-dissolving subjective effects, those expressing the emotional and transformative quality of embeddedness (e.g., spiritual, blissful, and unitive). We observed weaker associations for sensory-hallucinogenic effects and no association with negative effects (Fig. 3c).

This pattern supports interpreting embeddedness as a state central to psychedelics’ therapeutic relevance, potentially by addressing distress arising from separation. We use *separation* to mean the pervasive, felt disconnection between self and world (including self–other boundaries and rigid self-referential attachment). Interpreted this way, embeddedness marks a transient reduction of that separation—an existentially salient state that may help explain reported reductions in death-related anxiety and existential distress following psychedelic treatment (Griffiths et al., 2016; Ross et al., 2016, 2025), and, in our non-clinical cohort, is consistent with self- and boundary-dissolving experiences (Fig. 4c–g; Fig. Extended Data 12), increased personal meaning and death acceptance (Fig. Extended Data 13), and greater nature relatedness (Fig. Extended Data 14).

Further, the link between positively felt associative effects and improved mindset (Fig. 4h) was replicated using CEBRA embeddings, where classification accuracy was associated with next-day mindset change (*r* = 0.4, *p* < 0.01) (Fig. Extended Data 10). Although this association is likely mediated by subjective experience, it supports the view that brain embeddings capture meaningful experiential features of the psychedelic state that relate to subsequent psychological change. This suggests that brain-derived embeddings may offer a complementary approach for tracking subject-specific correlates of change where self-report is unavailable or unreliable.

The blissfully felt state of boundary-dissolving embeddedness may reflect a general mechanism of psychological adaptation—one that opens a sense of boundlessness and remediates separateness. Under supportive conditions, this can enable personally meaningful acute experiences that translate into persisting psychological benefits (Malone et al., 2018). This interpretation is consistent with therapeutic efficacy observed in depression (Roseman et al., 2018), and with participants’ reports of communitas, empathy, and nature relatedness—reflecting an attenuation of self–world boundaries characteristic of embeddedness (Pokorny et al., 2017; Kettner et al., 2021; Forstmann et al., 2020; Lyons and Carhart-Harris, 2018). Embeddedness thus offers a construct that may inform clinical approaches to mental health, by characterising the extent to which a person’s brain and mind are aligned with context under pharmacologically altered experience. The learned embedding spaces point to a deeper principle. Ordinarily, segregation between internally and externally directed systems buffers brain dynamics from direct environmental coupling, maintaining the separation between internal models and sensory context on which predictive processing depends (Ma et al., 2006). Under psilocybin, this boundary dissolves. Rather than maintaining separation from context, brain activity differentiates more coherently across contexts and becomes temporally coherent within each context (Fig. 3a,d).

These results address a key gap in systems neuroscience: how large-scale functional brain embeddings shift under multiple contextual demands in a pharmacologically altered state. Network perturbation analysis grounds this organisation anatomically, identifying Default Mode and Visual networks as its primary contributors (Fig. Extended Data 8). These results also position embeddedness as a data-driven construct that bridges measurable neural organisation with both acute and enduring effects, providing an alternative to accounts of psychedelic experience that are difficult to operationalise, such as ego dissolution (Stoliker et al., 2022).

Crucially, ML embedding techniques show that the prevailing interpretation of psychedelic RSN integration–segregation dynamics, commonly summarised by modularity changes, is incomplete. Reductions in within-network functional connectivity were underpinned by coherent within-network clustering (Fig. 3d), and modularity did not predict the individual differences in subjective experience or mindset change that CEBRA classification accuracy predicted (Fig. Extended Data 17), confirming that experientially meaningful neural organisation resides within the temporal dynamics that time-averaged summaries do not preserve.

By revealing structured, context-aligned organisation where previous studies found desynchrony (Siegel et al., 2024; Muthukumaraswamy et al., 2013), these embeddings recast the psychedelic state: apparent disorder in time-averaged measures masks dynamic organisation that emerges in proportion to subjective experience. This view is now supported at the circuit level by preclinical evidence that psilocybin selectively weakens cortico–cortical recurrent pathways while strengthening specific feedforward routes from perceptual and medial regions (the latter a rodent DMN homologue) in an activity-dependent manner (Jiang et al., 2025).

Although further research is needed to determine whether ML embedding signatures generalise across different populations, CEBRA demonstrates how dynamic functional trajectories can uncover structured organisation missed by static connectivity approaches, with broad applications in consciousness research and psychiatry.

### Context-aligned reorganisation of brain dynamics

Our analyses converge on a redistribution of integration under psilocybin—associative regions show increased global functional connectivity while sensory regions show reduced integration—that provides the systems-level basis for how context shapes brain dynamics during the psychedelic state. Time-averaged connectivity (fMRI) and spectral power (EEG) confirmed this redistribution across modalities, while temporally resolved trajectory analysis revealed that context increasingly differentiated neural dynamics as subjective effects intensified (Fig. Extended Data 7a–d), indicating that context-alignment is graded by experiential depth rather than imposed by external stimulation alone.

The GFC analysis demonstrated that psilocybin rebalanced sensory and associative connectivity. During eyes-closed conditions, sensory integration decreased while associative integration increased, consistent with a redistribution across the cortical hierarchy. Our large-sample size advanced this area of research by clarifying controversial findings from earlier small-sample serotonergic psychedelic studies, where spatial patterns varied across datasets and GSR (Preller et al., 2018, 2020; Timmermann et al., 2023). Further analysis using histograms of GFC values revealed that psilocybin reduced the separation between eyes-open and eyes-closed states, both globally and across individual networks, and was particularly evident in the Visual network (Fig. 2d). Similar sensory-region degree centrality reductions have been reported under LSD, MDMA and d-amphetamine Avram et al. (2025), indicating that sensory global-connectivity decreases can occur outside the class of serotonergic psychedelics.

Psilocybin also redistributed neural signal variability across sensory and associative regions. During eyesclosed states, variance increased in ventral, temporal, and somatosensory areas and decreased in early visual cortex, consistent with the redistribution of integration between sensory and associative regions observed in our GFC findings. These changes align with prior evidence of increased neural signal diversity and entropy under psychedelics across MEG/EEG, fMRI, and modelled dynamics (Schartner et al., 2017; Timmermann et al., 2019; Lebedev et al., 2016; Herzog et al., 2023), demonstrating how psilocybin reshapes spatial signal dynamics in a context-sensitive manner.

The 64-channel wet EEG provided a separate modality that confirmed the pattern of context-sensitive changes we identified in fMRI. The reduced alpha-band inhibition observed under psilocybin during eyes-closed states suggests a mechanism for the spontaneous production of internally generated visual effects (Kometer et al., 2013). Lempel-Ziv complexity increased brain-wide during eyes-closed conditions, extending previous evidence of these effects in occipital-parietal regions (Schartner et al., 2017). Meditation and music show visually similar group-averaged power spectra at baseline and under psilocybin (Fig. 6b); due to our fixed order design, we report these findings descriptively (see Methods). Similar reductions in signal diversity (Lempel-Ziv complexity) under external stimulation have also been reported under psychedelics (Mediano et al., 2024).

Therefore, while the spatial organisation of connectivity (fMRI surface maps) and neural dynamics (EEG scalp topographies) remains distinct from baseline (no-psilocybin), the overall distributions of connectivity strength (GFC values; Fig. 2d,e) and spectral power (EEG power; Fig. 6b) converge under psilocybin, indicating that eyes-closed states shift toward a more externally engaged profile while preserving condition-specific topology. This pattern reflects a redistribution of integration in which associative networks participate more broadly during eyes-closed states while sensory networks become less constrained by baseline organisation. Importantly, the convergence of sensory-state boundaries in time-averaged FC (Fig. 2b,d) is not a loss of structure but a reorganisation compatible with the context-aligned organisation captured by low-dimensional trajectory analysis (Fig. 3a). As rigid distinctions between eyes-open and eyes-closed states relax, temporally ordered dynamics become more distinctly locked to each context.

DCM analysis also linked these findings to emerging preclinical and human evidence of anterior hippocampus-DMN neuroplasticity following psilocybin (Siegel et al., 2024). Using DCM, we demonstrated that eyes-open vs eyes-closed context differentially tunes the directed connectivity of these brain circuits during the acute effects, with the eyes-open movie showing the greatest magnitude of directed change, in line with eyes-open vs. eyes-closed distinctions in our fMRI and EEG findings.

Machine-learning embeddings distinguished patterns of psychedelic brain activity across experimental contexts (rest, meditation, music, movie), with classification accuracy scaling with the intensity of acute subjective experience—particularly positively felt, immersive self- and boundary-dissolving dimensions (e.g., mystical, blissful, and unity). Together, increased integration of cortical–subcortical connectivity—including interactions between systems that ordinarily segregate internal (self-referential) and external (environment-focused) processing—modelled with TAVRNN, and the context separability uncovered by CEBRA, converge to support the construct of embeddedness: a state in which individuals feel fundamentally continuous with their environment, marked by diminished self–world separation. These signatures were detectable at the individual level and associated with next-day mindset improvements, providing a brain-derived marker of large-scale neural reorganisation that connects the quality of subjective experience to the change that follows.

Our findings extend prevailing accounts that characterise the psychedelic state as desynchronised or entropically disordered by revealing structured, context-aligned organisation in neural dynamics—a latent order harboured within temporal dynamics that time-averaging obscures—which emerges in proportion to the depth of self- and boundary-dissolving experience. This organisation is distributed across networks but depends on the joint alteration of Default Mode and Visual systems: the gradient endpoints that ordinarily segregate internal from external processing. When both are altered, that functional boundary dissolves, setting the conditions for brain activity to align with context, an integration that provides a neurobiological rationale for how structured settings can shape outcomes under psychedelics.

That the felt boundary between self and world reorganises when these dynamics shift implies that the boundary is not a fixed datum of conscious experience but a construction actively maintained by neural activity. Embeddedness—the continuity that emerges when this construction relaxes—may thus reveal less about what psychedelics add to consciousness than about what ordinary neural dynamics keep apart.

## Methods

### Ethics and clinical trial registration

The study protocol was approved by the Monash University Human Research Ethics Committee. The trial was registered with the Australian New Zealand Clinical Trials Registry under the registration number ACTRN12621001375842.

### Study design

The PsiConnect study was open-label and included two imaging sessions: a baseline (no-psilocybin) session and a session following the administration of a 19 mg dose of psilocybin. Both sessions involved MRI and EEG scans, with four conditions repeated in each: resting state, guided meditation, music listening, and movie watching. In fMRI, conditions began ~80 minutes post-dose and followed a fixed order (rest → meditation → music → movie). The sequence progressed from low-stimulus to high-stimulus contexts to prioritise safety and experiential coherence for psychedelic-naïve participants. The EEG session started after a room transfer and 20–40 minutes of setup (cap placement and impedance checks), with the movie presented first, followed by the three eyes-closed conditions (movie → rest → meditation → music). Discussion addressing expectancy, task order and EEG timing is detailed in Supplementary Information.

In the resting-state condition (8 min MRI, 5 min EEG), participants were instructed to relax and keep their eyes closed while remaining still. During the guided meditation (6:30 min MRI, 5 min EEG), participants received brief instructions via MRI-safe audio, followed by silent periods of meditation. For the music listening condition (11:24 min MRI, 7 min EEG), a curated playlist was designed to evoke emotional depth and resonance. In the naturalistic movie condition (6 min MRI, 5 min EEG), participants watched a video of moving clouds without audio. EEG blocks were intentionally shorter (total ≈22 min) to acquire all four contexts within the acute-effects window and to minimise fatigue, motion, and impedance drift; see the accompanying PsiConnect data descriptor (Novelli et al., 2025). These conditions were studied in both the baseline (no-psilocybin) and psilocybin sessions and were repeated in both fMRI and EEG, allowing for comprehensive cross-modal and longitudinal analyses of brain activity and connectivity.

The eligible participants were stratified based on age and self-reported gender before allocation to the mindfulness meditation and control groups using a non-randomised, balanced procedure. Half of the participants were assigned to an 8-week mindfulness-based cognitive therapy (MBCT) program *Finding Peace in a Frantic World* run by a trained and registered instructor, which involved weekly group meetings and daily independent practice; the other half were assigned to a control group with no intervention. Each participant was scanned at both baseline (no-psilocybin) and under psilocybin. Because no statistically significant differences were observed between meditators and non-meditators in connectivity or subjective effect measures under psilocybin (see Impact of mindfulness meditation training), the two groups were pooled for all analyses reported here.

Our decision to administer a standardised dose of 19 mg psilocybin, rather than a body-weight-adjusted dose was based on previous research showing no clear advantage of weight-adjusted dosing in terms of subjective effect intensity or predictable differences in response across individuals of varying body weight (Garcia-Romeu et al., 2021). The dosage was administered as one oral capsule and selected in consultation with multiple collaborators who had previous psychedelic imaging experience. This dose was determined to be tolerable for the majority of healthy adults undergoing imaging procedures, while also sufficient to produce significant subjective effects.

Several behavioural measures were collected before and during the baseline (no-psilocybin) and psilocybin scans (see Reported measures). The follow-up conducted the day after psilocybin administration included semistructured, open-ended questions and experience ratings. Further follow-up measures were administered one week, and one, three, six and twelve months after psilocybin administration.

### Participants

Sixty-five healthy adults aged 18–55 (37.3 ± 10.7; 32 female, 33 male) with no psychedelic experience were recruited. Sex was recorded at imaging intake and is reported here; self-reported gender, which was used for stratification between meditators and non-meditators and is concordant with sex for all but one participant, is detailed in the Reporting Summary. Participants were required to have no formal meditation practice and limited previous exposure to meditation. They were first screened via short online survey and then detailed screening was performed by a suitably trained staff member for excluding any psychopathology using the long form SCID-V (First et al., 2015). Exclusion criteria included a history of psychiatric disorders or suicidality; a 5-year history of substance and/or alcohol use disorder; first-degree relatives with a diagnosed psychotic disorder; a history of major neurological disorders including stroke or epilepsy; formal meditation practice within the last 6 months or extensive prior exposure to mindfulness meditation; use of contraindicated medications; and any hallucinogen use within the past 6 months. Participants were also screened for MR contraindications and provided informed consent. We operationally defined ‘no psychedelic experience’ as no prior serotonergic psychedelic use with subjective effects. During structured screening, three participants disclosed nominal or remote lifetime exposures without subjective effects: two reported ineffective microdoses, and one reported remote exposure over 20 years prior with minimal or no recall. These cases are detailed in Supplementary Information. All other participants (n=59) reported no lifetime use across online, phone, and SCID checks. On the day of psilocybin administration, they were assisted by a study doctor, researchers, lab staff with relevant training, and volunteers from the community drug harm reduction support organisation “Dancewize”. Psilocybin was generally well tolerated, although some adverse effects were reported, including reports of transient headaches during the night (n=3). Three participants received follow-up support from a clinical psychologist familiar with psychedelic integration. These follow-up calls were conducted over the phone, and no further support was required. On no occasion was it deemed necessary by the study doctor to administer an anxiolytic.

Of the 65 participants enrolled, two were withdrawn before the psilocybin session after meeting exclusion criteria identified during the study period. Their baseline data were not analysed, leaving a baseline sample of 63 per condition for both fMRI and EEG. No baseline fMRI or EEG recordings required further exclusion on quality control. The remaining 63 participants received psilocybin. Of these, one did not complete any post-dose imaging or EEG, and a second partially completed the resting-state fMRI scan (which was retained) but did not complete EEG. This gave post-dose fMRI samples of 62 (rest) and 61 (meditation, music, movie). Two further participants did not complete the post-dose EEG session, giving 59 completed EEG recordings per condition. Quality control applied pre-established thresholds for head motion and signal quality (see MRI quality control). During the psilocybin session, seven participants had one or more fMRI conditions excluded (2 rest, 2 meditation, 5 music, 4 movie), resulting in final analysed fMRI samples of: rest = 60, meditation = 59, music = 56, movie = 57. For machine-learning analyses (CEBRA, TAVRNN), participants were included only if all four conditions passed quality control with complete acquisition (expected volume count across the scanning session), giving a balanced sample of n = 54. For psilocybin EEG, one recording was excluded due to missing event markers (final n = 58 per condition). A full table of per-participant quality control exclusions is included in the reporting summary and data descriptor (Novelli et al., 2025).

### Reported behavioural measures

Reported measures were part of a broader assessment conducted before and longitudinally after psilocybin administration, with a subset integrated into the present analysis. 11-Dimensional Altered States of Consciousness (11D-ASC) scale (42 items) (Dittrich, 1998; Studerus et al., 2010) was administered post-EEG at baseline and at the end of the psilocybin session (approximately 320 minutes after dose) to assess subjective alterations in consciousness. The Mystical Experience Questionnaire (MEQ30) (Barrett et al., 2015) was also collected at this time to measure mystical-type experiences, along with the Aesthetic Experience Scale–Music (AES-M), used to assess the intensity of emotional and aesthetic responses to music. The AES-M scale is derived from the Aesthetic Experience Scale (AES) (Stamatopoulou, 2004), refined through psychometric validation (Silvia and Nusbaum, 2011), and adapted for music-related experiences (Sachs et al., 2016). We adapted the instructions of this measure for collection on the day of psilocybin by asking participants to respond with reference to their musical experience during the psilocybin session. The order of scale presentation was randomised. A music experience measure (Kaelen et al., 2018) assessed participants’ ratings of liking, openness, and resonance with the music. It was administered after both the MRI and EEG, on both session days. Scores were averaged to produce a single music experience rating.

One day after psilocybin, participants provided self-reported intensity ratings assessing the perceived strength of the experience. To assess the perceived meaningfulness of the experience, participants were asked: “Would you rate the experience among the most meaningful and spiritually significant experiences of your life?” If they responded yes, they were then asked: “Where would you rate the experience among the most meaningful and spiritually significant experiences of your life?” Response options included top 200 (n=1), top 100 (n=0), top 50 (n=3), top 10 (n=9), and top 5 (n=24) (Griffiths et al., 2006). A novel mindset measure capturing psychological change was also collected. See Supplementary Information for mindset measure details.

The Nature Relatedness Scale (NR-6) (Nisbet and Zelenski, 2013) and Life Attitudes Profile–Revised (LAP-R) (Reker, 1992)—assessing death acceptance, coherence, and purpose—were collected before psilocybin and one month post-administration to examine changes in ecological connectedness and existential meaning.

Colour groupings (blue/orange/red/purple) shown in Fig. 3c and Fig. 4c,e,g,h were assigned post hoc based on within-sample intercorrelations Fig. Extended Data 12 and theoretical rationale; they are descriptive and are not a validated taxonomy.

### MRI acquisition

Structural and functional MRI data were acquired using a Siemens 3 Tesla Magnetom Skyra scanner at Monash Biomedical Imaging, Monash University, Australia. T1-weighted (T1w) anatomical images were obtained for each participant during two sessions: at baseline (no-psilocybin) and on the psilocybin administration day. The images were acquired using a 3D magnetisation-prepared rapid gradient-echo (MP-RAGE) sequence with a 32-channel head coil. The acquisition parameters were as follows: repetition time (TR) of 2300 ms, echo time (TE) of 2.07 ms, 192 slices per slab, 1 mm slice thickness, and 1 mm isotropic voxel size. The parallel acquisition technique was GRAPPA. For the structural T1-weighted scan only, the acceleration factor (PE) was 2 at baseline (5:12) and 3 on the psilocybin administration day (3:52) to minimise time under drug. A T2-weighted anatomical image was obtained only during the baseline session. The acquisition parameters were: TR of 3200 ms, TE of 452 ms, 176 slices per slab, 1 mm slice thickness, and 1 mm isotropic voxel size, with an acceleration factor of 2.

Blood-oxygenation-level-dependent (BOLD) fMRI data were collected using a multi-echo, multi-band, echoplanar imaging, T2*-weighted sequence. The acquisition parameters were: TR of 910 ms, multi-echo TE of 12.60 ms, 29.23 ms, 45.86 ms, 62.49 ms, multi-band acceleration factor of 4, field of view of 206 mm, RL phase encoding direction, and 3.2 mm isotropic voxels. The scan durations were: resting state with eyes closed (8 minutes, 505 volumes), audio-guided meditation with eyes closed (6:30 minutes, 405 volumes), music listening with eyes closed (11:24 minutes, 728 volumes), movie watching (6:00 minutes, 372 volumes).

The structural and functional MRI images acquired from the Siemens scanner were converted into the Neuroimaging Informatics Technology Initiative (NIfTI) format and organised according to the Brain Imaging Data Structure (BIDS) 1.7.0.

### MRI quality control

Quality control was performed using the MRIQC BIDS app (Esteban et al., 2017), which uses structural and functional MRI images to compute several quality metrics. These include the temporal signal-to-noise ratio and the frame-wise displacement that quantifies head motion (Power et al., 2014). Principal component analysis of these metrics revealed some outliers in the structural and functional images, which were further inspected visually. This process led to the exclusion of 6 structural T1w images from the psilocybin session, while all the T1w images from the baseline session were retained. For the functional MRI data, 7 participants had one or more conditions excluded from the analysis (rest = 2, meditation = 2, music = 5, movie = 4). The list of excluded scans per participant is available in the PsiConnect data descriptor (Novelli et al., 2025). Most of these outliers corresponded to scans where the head motion was large (mean frame-wise displacement greater than 0.5, which is often used as an exclusion threshold (Parkes et al., 2018)). The final analysed MRI samples were: rest = 60, meditation = 59, music = 56, and movie = 57.

### MRI preprocessing and cleaning

Anatomical and functional MRI data were preprocessed using *fMRIprep* 22.0.2. The anatomical MRI preprocessing involved correcting T1-weighted images for intensity non-uniformity, skull-stripping, and segmenting brain tissues. The images were then registered, brain surfaces reconstructed, and spatial normalisation performed using the ICBM 152 Nonlinear Asymmetrical template version 2009c (MNI152NLin2009cAsym). The functional MRI preprocessing involved generating a reference volume and skull-stripped version from the shortest echo of each BOLD run, estimating head-motion parameters, and performing slice-time correction. The BOLD reference was co-registered to the T1w reference, confounding timeseries were calculated, and the BOLD time-series were resampled into MNI152NLin2009cAsym space.

The preprocessed and optimally combined data produced by *fMRIprep* was cleaned via a single regression in *SPM12* (John Ashburner et al., 2020) using the general linear model. The regressors were: the white matter and cerebrospinal fluid signals computed by *fMRIprep*, the frame-wise displacement, and the non-BOLD components identified via multi-echo ICA performed using *tedana* (version 0.0.12) (DuPre et al., 2021). In short, multi-echo ICA identifies non-BOLD components that are independent of the echo time (Kundu et al., 2012). Very low-frequency components (especially the linear trend) were implicitly filtered out after cleaning (Novelli et al., 2025), which provides further evidence that multiecho ICA was able to identify spurious components in the data. Finally, to enable surface-based analyses, the cleaned volumetric data were projected to the FreeSurfer left-right-symmetric cortical surface template with 32000 vertices for each hemisphere (fsLR32k).

### Standard deviation maps

The standard deviation (SD) was computed for the BOLD time series of each vertex on the cortical surface. The percentage change (psilocybin vs no-psilocybin) was computed for each participant and then averaged across participants to obtain the mean standard deviation percentage change map shown in Fig. 2c. A surface-based general linear model was used to assess whether changes in BOLD SD were attributable to head motion, and in which cortical areas. For each condition and hemisphere, the vertex-wise SD percentage change between baseline and psilocybin (ΔSD_*v*_) was regressed on subject-level framewise displacement (FD) change using the following model: 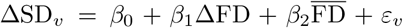, where ΔFD is the within-subject change in mean FD and 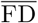 is the mean FD across sessions. Inference on *β*_1_ used nonparametric permutation testing with family-wise error correction using FSL PALM. Significant positive associations between SD change and FD change were confined to regions near air-tissue interfaces (orbitofrontal cortex, anterior temporal lobe). The corresponding voxels were masked out in Fig. 2c, so that only SD changes that were not significantly associated with head motion remain. These motion-related effects were specific to SD, and the spatial organisation of GFC changes was not significantly associated with framewise displacement.

While standard deviation and Shannon entropy are distinct, there is a monotonic relationship between the two when the data distribution is Gaussian. More generally, these two quantities are tightly related when the data distribution is continuous, unimodal, and has finite variance, such as in fMRI data. Variance is bounded above and below by scaled entropy powers, so that as one increases, the other is constrained to increase within bounded limits (Chung et al., 2017). Accordingly, under these conditions, greater standard deviation corresponds to greater Shannon entropy.

### Global functional connectivity maps

For each cortical vertex, Pearson’s correlation to all other vertices in the same brain hemisphere was computed, transformed to Fisher z-values, and averaged. This calculation yielded a global functional connectivity (GFC) map where each vertex value represents the mean correlation with all other vertices in the same hemisphere (Cole et al., 2010). This generated one map for each participant, in each session (baseline and psilocybin), and each condition (resting-state, meditation, music, movie). The histograms of GFC values were used to compare the sessions and conditions, ignoring the spatial distribution across the cortex (Fig. 2e). To appreciate the spatial distribution, the GFC percentage change was computed (psilocybin day minus baseline, then divided by baseline) for each participant and then averaged across participants, yielding the mean GFC percentage change maps shown in Fig. 2a.

Threshold-free cluster enhancement (TFCE) (Smith and Nichols, 2009) was used to identify significant clusters in the spatial maps without defining arbitrary thresholds for cluster size. Intuitively, TFCE tests all thresholds and gives higher scores to clusters of vertices that are both large and survive increasingly stringent thresholds. TFCE was computed using the PALM software with the default parameters (see https://github.com/andersonwinkler/PALM/blob/master/palm_defaults.m). The statistical TFCE maps were thresholded at *p* < 0.05 with 1000 permutations and a Gamma approximation for the distribution tail (computing permutations is computationally very costly; in our tests, the approximation produced nearly identical clusters to those obtained using 10000 permutations but reduced the computation time by one order of magnitude). Once the clusters of increased and decreased GFC were obtained, they were used to mask the GFC maps of the effect of psilocybin (percentage change) to hide the vertices outside these clusters (Fig. Extended Data 1). Effect sizes are reported as Cohen’s *d*, computed using the Algina–Keselman–Penfield robust estimator, which uses trimmed means and Winsorised variances to reduce sensitivity to outliers and non-normality (Algina et al., 2005).

### A note on global signal regression and z-scoring the global functional connectivity

We recommend not performing global signal regression (GSR) when computing the GFC, although it has been common practice in previous research (Preller et al., 2020, 2018; Timmermann et al., 2023). Mathematically, GSR alters the covariance matrix of the data such that the average of each row is zero (Murphy et al., 2009). Since GFC is computed by averaging the rows, it would become zero for each vertex across the cortex. The reason why GFC is not exactly zero in the literature using GSR is that the Pearson correlation matrix is used instead of the covariance matrix. The correlation matrix is a rescaled version of the covariance matrix where all the entries are divided by the standard deviations so that they are between −1 and 1. To achieve this normalisation, each entry *σ*_*ij*_ of the covariance matrix is divided by the product of the standard deviation of vertices *i* and *j*, that is, by 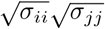. If all the vertices had the same standard deviation, this normalisation would equally rescale all entries and the average of each row of the correlation matrix would still be zero. In practice, the standard deviation varies across cortical vertices (Fig. 2c) so the average of each row of the correlation matrix is not exactly zero, but typically a small value. We argue that this complicates the interpretation of the GFC, confounding the intended goal of quantifying “the extent to which a cortical vertex is positively/negatively correlated with the other vertices, on average” with “the extent to which a vertex is positively/negatively correlated with vertices having large standard deviation, on average”.

Relatedly, plotting z-scored GFC values confounds the interpretation in two ways. If GSR is applied, z-scoring hides the fact that GSR significantly reduces the magnitude of GFC values. More generally, z-scoring is problematic when used to report GFC differences between two conditions. For example, in the main text, we have reported the percentage change between the psilocybin and baseline conditions (Fig. 2a). Positive and negative values indicate an increase or decrease in GFC, respectively. After z-scoring the changes, this interpretation is no longer possible: a positive z-score only indicates that the change is higher than average, regardless of its original sign (see Fig. Extended Data 3). If the average were negative, negative values slightly above the average would have a positive z-score.

### Parcel-wise FC

The denoised, volumetric data were divided into parcels for further functional connectivity analysis. The Schaefer parcellation (Schaefer et al., 2018) was used to partition the cortical voxels into 300 parcels, each belonging to one of the following 7 resting-state networks: Visual, Somato-motor, Dorsal Attention, Limbic, Salience/Ventral Attention, Default Mode, Control. The Melbourne subcortical atlas (Tian et al., 2020) was used to divide the subcortical voxels into 32 parcels including subdivisions of the striatum, thalamus, hippocampus, amygdala and globus pallidus. (This atlas is specified in voxel space, hence the use of volumetric rather than surface data for this analysis.) Combining the cortical and subcortical parcellations gave a total of 332 parcels. To extract a representative time series for each parcel, the first principal component of the voxel time series within each parcel was computed.

The parcel-wise functional connectivity (FC) matrix was obtained by computing Pearson’s correlation between the time series of each pair of parcels, and the average FC matrices across participants are shown in Fig. 2b. The within-network FC was computed by averaging the FC values within the diagonal blocks of the FC matrix, each corresponding to the pairwise correlations between parcels belonging to the same functional network. Similarly, the between-network FC was computed by averaging the FC matrix entries outside the diagonal blocks, i.e., the correlation values between pairs of parcels belonging to different networks (see Supplementary Information). The Mann-Whitney U-test was performed in MATLAB using the ‘ranksum’ function and the right-tailed option. That is, the alternative hypothesis is that the median within- or between-network FC under psilocybin is lower than the baseline median. To correct for multiple comparisons across eight resting-state networks, the statistical significance was set to *α* = 0.05*/*8 = 0.00625 (Bonferroni correction).

### Modularity

The FC modularity was computed using the Brain Connectivity Toolbox (Rubinov and Sporns, 2010) with the ‘negative asym’ option for asymmetric treatment of negative weights (Rubinov and Sporns, 2011). The Mann-Whitney U-test was performed in MATLAB using the ‘ranksum’ function and the right-tailed option. That is, the alternative hypothesis is that the median modularity under psilocybin is lower than the baseline (no-psilocybin) median (Fig. 2f). To correct for multiple comparisons across four conditions, the statistical significance was set to *α* = 0.05*/*4 = 0.0125 (Bonferroni correction).

### Spectral dynamic causal modelling

Spectral dynamic causal modelling (DCM) is a method used to infer the effective connectivity between brain regions from fMRI time series (Friston et al., 2014; Razi et al., 2015; Novelli et al., 2024). Effective connectivity is defined as the influence one neural system exerts over another (Friston et al., 2003), as opposed to functional connectivity, which describes statistical dependencies among BOLD signals (Friston, 2011; Razi and Friston, 2016). DCM employs a forward generative model with neuronal and observation equations. The neuronal model is a linear stochastic differential equation that describes the dynamics of hidden neuronal states. It models how the activity in one brain region is influenced by the activity in other regions, as well as by endogenous fluctuations in neuronal activity. The observation function maps the neuronal activity to the observed BOLD signal by modelling the biophysical processes involved in the haemodynamic response. Spectral DCM estimates the parameters of this model by fitting the generative model to the cross-spectral density of the observed BOLD signals, a second-order statistic that captures the correlations between time series at all time lags (Novelli et al., 2024). By fitting the model to the cross-spectral density, spectral DCM can estimate the effective connectivity between brain regions, as well as the hemodynamic parameters and the spectrum of the endogenous fluctuations. Because DCM is a Bayesian approach, all model parameters are equipped with prior distributions, which are updated based on the observed data to produce posterior distributions over the parameters. Spectral DCM explicitly models the endogenous fluctuations in neuronal activity, which makes it ideal for applications to the resting state and conditions that do not involve strong experimental inputs or block task designs.

Here, we applied spectral DCM to infer the effective connectivity of each subject in each condition. The analysis focused on six regions belonging to the DMN, each modelled as a 6 mm sphere centred on the following coordinates: Left anterior hippocampus (aHip L; X=-26, Y=-16, Z=-20), right anterior hippocampus (aHip R; X= 28, Y=-16, Z=-20), left inferior parietal cortex (IPC L; X=-44, Y=-60, Z=24), right inferior parietal cortex (IPC R; X=54, Y=-62, Z=28), medial prefrontal cortex (mPFC; X=2, Y=56, Z=-4), and posterior cingulate cortex (PCC; X=2, Y=-58, Z=30). Subject-level fully connected DCMs were combined at the group level with Parametric Empirical Bayes, which precision-weights individual parameter estimates by their posterior uncertainty. Specifically, Parametric Empirical Bayes (Zeidman et al., 2019) was used to compute the group-level effective connectivity at baseline and its change under psilocybin (Fig. 5). All DCM analyses were performed using SPM12 (John Ashburner et al., 2020). See Supplementary Information for further details.

### CEBRA

Contrastive Embeddings for Behavioural and Neural Analysis (CEBRA) is a self-supervised learning framework that extracts interpretable and consistent low-dimensional embeddings from high-dimensional neural and behavioural datasets (Schneider et al., 2023). By leveraging contrastive learning and auxiliary variables, such as time or behavioural labels, CEBRA enables analyses of population-level trajectories.

In this study, we used CEBRA-Time, a variant of CEBRA tailored for temporal analyses, to explore neural trajectories derived from fMRI data. CEBRA-Time is a contrastive learning-based method designed to extract meaningful latent representations from time series data, such as fMRI signals. It learns a low-dimensional embedding by maximizing temporal coherence while preserving task-related variability, allowing the transformation of high-dimensional neural data into structured latent trajectories. This enables the discovery of subject-specific neural dynamics in a more interpretable space.

CEBRA is well-suited for fMRI data, as it summarises the activity of all brain regions (ROIs) at each moment into a compact representation of the overall brain state and embeds these points in a lower-dimensional space. Each point in this space corresponds to a single fMRI volume—that is, a whole-brain snapshot across ROIs. The embeddings themselves are learned directly from the fMRI BOLD signals. Training is guided by a contrastive objective: volumes that are similar are encouraged to map closer together in the embedding space, while more dissimilar volumes are pushed farther apart. Plotting successive points yields a trajectory that makes shifts between rest, meditation, music, and movie watching easier to see and quantify.

Here, for each participant, we concatenated fMRI time series spanning four consecutive conditions: resting state, guided meditation, music listening, and movie (Fig. 3a). This comprehensive input captured temporal transitions in neural activity across conditions. Applying CEBRA-Time transformed each time point of these high-dimensional fMRI time series into a three-dimensional point in latent space, revealing neural trajectories unique to each subject (Fig. 3a,b). These trajectories provided insights into how dynamic brain activity evolves over different conditions, aligning with self-reported subjective experiences. We also confirmed that CEBRA embeddings were stable across repeated runs in our dataset (Fig. Extended Data 7e,f), ensuring robustness of the trajectories presented here.

Thus, while even more conventional ML embedding methods confirm the separability of brain states across conditions (see Fig. Extended Data 9), CEBRA provides a richer, temporally coherent view of how these states evolve into organised patterns of brain activity, making it uniquely suited to studying continuous experiential processes like those induced by psychedelics.

### Network perturbation analysis

To identify which functional systems contribute most to the context-aligned trajectory organisation revealed by CEBRA, we performed a network perturbation analysis using a baseline substitution approach. For each participant, the psilocybin time series of a given functional network was replaced with that participant’s corresponding baseline (no-psilocybin) dynamics, while all other networks retained their psilocybin activity. The CEBRA embedding and SVM classification were then re-run on this hybrid time series, and the drop in classification accuracy was measured. This procedure isolates the contribution of each network’s psilocybin-specific modulation to the overall context-aligned organisation, while preserving the natural statistical structure of the individual’s biological signal. Only positive contributions were retained when constructing an additive attribution, ensuring that individual network contributions sum to the total psilocybin-induced accuracy gain (see Supplementary Information for the full attribution formula). The analysis was performed on each individually trained CEBRA model to assess whether the anatomical anchoring was consistent across participants. This attribution quantifies each network’s contribution to the overall multi-context organisation; it does not assign unique network generators to individual context contrasts.

To further characterise condition-specific contributions, we computed per-condition recall (the fraction of time points truly belonging to a given condition that were correctly classified) from the confusion matrix of the SVM classifier trained on psilocybin embeddings. We then repeated inference after each network substitution and extracted the change in per-condition recall (diagonal of the row-normalised confusion matrix). The across-subject mean change in recall (psilocybin minus substituted) is reported for each network and condition.

### TAVRNN

While CEBRA captures overall neural trajectories, a deeper understanding requires analysing how individual brain regions (ROIs) evolve over time and interact with each other. TAVRNN allows us to track these ROI-specific dynamics and their changing patterns of interaction (i.e. functional connectivity), revealing finer details. This captures detailed, context-dependent communication between regions that is undetectable by static analyses.

This perspective inspired our application of Temporal Attention-enhanced Variational Graph Recurrent Neural Network (TAVRNN)(Khajehnejad et al., 2024). TAVRNN is a framework designed for analysing the temporal dynamics of evolving connectivity networks. TAVRNN is a deep learning framework that models temporal connectivity networks between units of a system by leveraging both structural relationships and past dynamics. Through temporal attention, recurrent neural networks, and variational graph techniques, it learns a lower-dimensional latent representation for each unit (ROIs in this work), preserving meaningful temporal patterns while enhancing interpretability. This allows for tracking the evolution of ROI activities over time, revealing insights into their dynamic interactions. TAVRNN captures both local temporal patterns (how individual ROIs change over time) and global temporal patterns (how the relationships as a whole evolve dynamically). By leveraging latent information from the structure of time-varying functional connectivity networks, TAVRNN enables a robust representation of individual ROI activity and their changing relationships in a low-dimensional space (Fig. 3d). Specifically, a temporal attention mechanism assesses the topological similarity of the network across time steps, incorporating varying time lags to capture complex network dynamics more effectively. This approach is particularly effective in uncovering temporal changes in network structures and their alignment with external behaviours or stimuli.

In our study, we used TAVRNN to analyse FC matrices derived from fMRI time series under four consecutive conditions: rest, meditation, music, and movie. For each condition, we computed the FC matrices between 332 brain parcels, which served as input nodes to TAVRNN (n = 54; the same participants used in the CEBRA analysis). No segmentation or overlapping windows were applied; each FC was computed using the full time series for that condition to capture stable, condition-specific network states. In this work, each scan under a specific condition (rest, meditation, music, or movie) was treated as a temporal snapshot. Therefore, the attention mechanism enables the lower-dimensional representation of one scan to inform the representation of the subsequent scans. The model then generated two-dimensional embeddings for each parcel, representing their temporal evolution across conditions (Fig. 3d).

For further details on biological interpretability of the latent spaces derived from our machine-learning analyses, and conventional and machine-learning metric complementarity, see Supplementary Information.

### EEG acquisition

EEG data were recorded using a 64-channel BrainAmp MR Plus amplifier (Brain Products GmbH, Germany) with BrainVision Recorder software (version 1.22.0001, Brain Products GmbH, Germany) and Ag/AgCl electrodes embedded in an actiCAP slim cap according to the standardized 10-10 system (Chatrian et al., 1985). The FCz electrode served as the reference and the FPz electrode served as the ground for online recording. The impedances were maintained below 10*k*Ω using an abrasive paste (Nuprep, Weaver and Company, USA) followed by applying a conductive gel (Easycap GmbH, Germany). EEG signals were sampled at 500 Hz with a bandpass filter of 0.01 −1000*Hz*. Participants were comfortably seated in an acoustically absorbent room to minimise environmental noise and their chin strap was securely fastened to ensure head stability. They were instructed to remain as still as possible, avoid jaw clenching, and keep their eyes closed during the recording to minimise artifacts.

The EEG recording session included the same four conditions as the fMRI described above. The naturalistic movie (5 minutes) was presented first, followed by the three eyes-closed conditions: resting state (5 minutes), audio-guided meditation (5 minutes), and music listening (7 minutes). The raw data were acquired as a single continuous recording over all conditions and then split into four separate files using the event markers for the start and end of each condition. Due to technical issues, event markers were not recorded for two participants in the baseline session and one participant in the psilocybin session, so these recordings were excluded from pre-processing and analysis. In summary, pre-processed EEG were analysed for 63 participants in the baseline session and 58 in the psilocybin session.

### EEG pre-processing

The raw EEG signals were preprocessed using the default parameters of the automated RELAX pipeline (Bailey et al., 2023) implemented in MATLAB using functions from EEGLAB (Delorme et al., 2004) and FieldTrip (Oostenveld et al., 2011a) toolboxes. First, the data were bandpass filtered between 0.25 and 80 Hz using a fourth-order Butterworth filter with zero phase, with a notch filter applied between 47 and 53 Hz to reduce line noise. Subsequently, bad channels were removed by applying a multistep process incorporating the “findNoisyChannels” function of the PREP pipeline (Bigdely-Shamlo et al., 2015). This was followed by the initial reduction of the artefacts related to eye movements, muscle activity and drift using a multi-channel Wiener filter (Somers et al., 2018), after which the data were re-referenced to the robust average reference (Bigdely-Shamlo et al., 2015). Residual artefacts were then removed by Wavelet-enhanced Independent Component Analysis (WICA) with the artefactual components identified by the automated ICLabel classifier (Hill et al., 2022). Electrodes rejected during the cleaning process were then restored to the data through spherical interpolation. Finally, all preprocessed data were visually inspected for data quality.

### EEG data analysis

The power spectrum was computed for all electrodes across all participants using the multitaper frequency transformation (1 −45 Hz) with a 2-second Hanning window and 50% overlap, implemented in FieldTrip (Oostenveld et al., 2011b). To quantify the effect of psilocybin, the power difference (psilocybin minus baseline) was computed for each channel and then averaged across subjects. The results are reported in Fig. 6 for each condition and four power bands: theta (4 − 7 Hz), alpha (8 − 12 Hz), beta (13 − 30 Hz), and gamma (30 − 80 Hz).

The Lempel-Ziv (LZ) complexity was calculated in two steps using the LZ 1976 algorithm (Lempel and Ziv, 1976). First, the time series for each channel was binarised by setting values above the mean to 1 and values below the mean to 0. The resulting binary sequence was then scanned sequentially to identify distinct patterns and build a dictionary of these patterns. The LZ complexity is given by the number of patterns in the constructed dictionary. Regular signals can be represented by a small number of patterns, resulting in low LZ complexity, while irregular signals have a greater diversity of patterns, leading to higher LZ complexity.

## Supplementary Information

See attached file.

## Acknowledgements

The authors acknowledge the facilities and scientific and technical assistance of the National Imaging Facility (NIF), a National Collaborative Research Infrastructure Strategy (NCRIS) capability at Monash Biomedical Imaging (MBI), a Technology Research Platform at Monash University. This research is funded by the Australian Research Council (Refs: DE170100128 and DP200100757) and the Australian National Health and Medical Research Council (Investigator Grant 1194910). A.R. is affiliated with The Wellcome Centre for Human Neuroimaging, supported by core funding from Wellcome [203147/Z/16/Z]. A.R. is a CIFAR Azrieli Global Scholar in the Brain, Mind & Consciousness Program. A.K.S is supported by European Research Council Advanced Investigator Grant CONSCIOUS (Ref: 101019254). K.H.P. is currently an employee of Boehringer Ingelheim GmbH & Co KG. We also acknowledge USONA Institute for providing psilocybin through their “Investigational Drug Supply Program”. This research was supported by Monash eResearch capabilities, including HPC (M3/MASSIVE).

## Data availability

All data reported has been uploaded to OpenNeuro (https://openneuro.org/datasets/ds006110/versions/1.1.0) and will be made fully open access upon publication.

## Code availability

Open-source code for all data analysis pipelines will be lication pipelines fMRIprep made available on GitHub upon pub-(https://github.com/razilab). These used the following software packages: 22.0.2 (https://fmriprep.org), tedana 0.0.12 (https://tedana.readthedocs.io), MRIqc 22.0.6 (https://mriqc.readthedocs.io), MATLAB R2022a (https://www.mathworks.com), SPM12 (https://www.fil.ion.ucl.ac.uk/spm/software/spm12), Freesurfer 7.2 (https://surfer.nmr.mgh.harvard.edu), FSL v.6.0.7 (https://fsl.fmrib.ox.ac.uk/fsl/fslwiki), FieldTrip 20240916 (https://www.fieldtriptoolbox.org). LZ complexity was computed by adapting open-source code by Fernando Rosas and Pedro Mediano (https://information-dynamics.github.io/complexity/information/2019/06/26/lempel-ziv.html). Projection of volumetric fMRI data to the cortical surface was performed by adapting open-source code by Pang et al. available on GitHub (https://github.com/NSBLab/BrainEigenmodes).

## Extended Data

**Fig. Extended Data 1.**
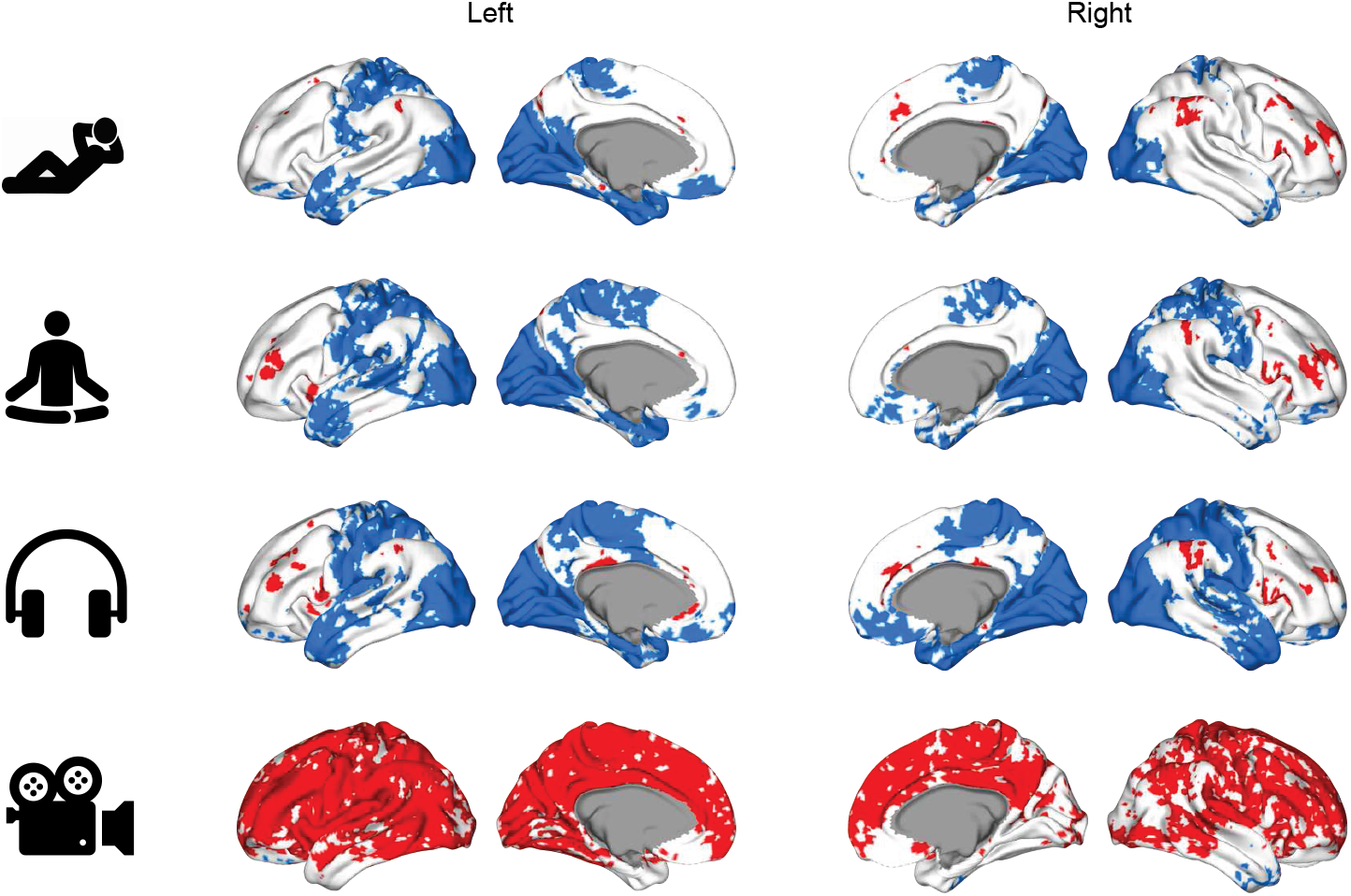
Clusters of statistically significant global functional connectivity increase (red) or decrease (blue) at *p* < 0.05 after threshold-free cluster enhancement.

**Fig. Extended Data 2.**
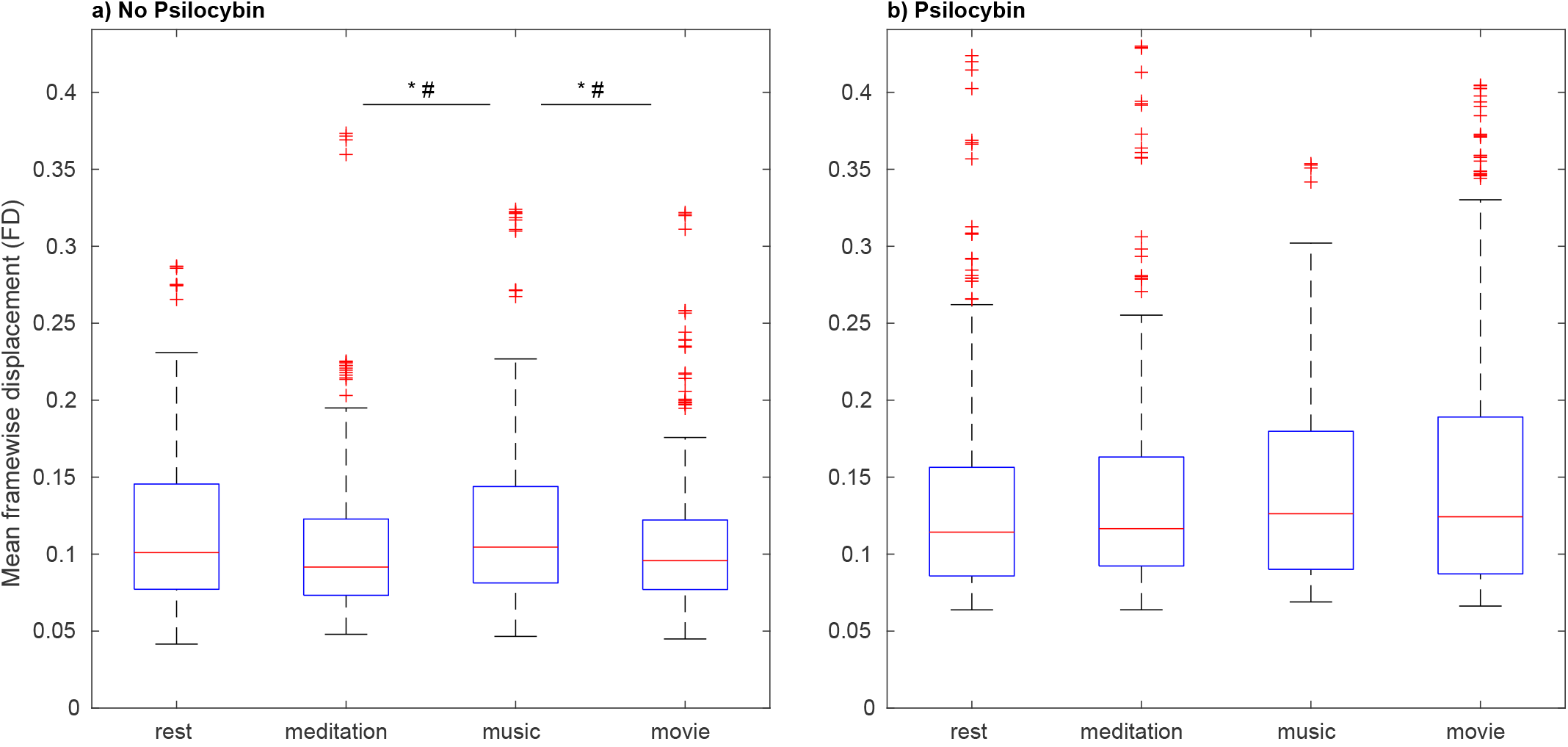
Head motion comparison across conditions within session, quantified via the mean frame-wise displacement (FD). a) Baseline session. Mean FD increased modestly between guided meditation and music listening (Mann-Whitney U-test, two-sided, *p* = 0.003) and decreased between music listening and movie watching (*p* = 0.004); both differences, though statistically significant, were small in magnitude. b) Psilocybin session. No statistically significant differences were observed between conditions (all *p* > 0.18).

**Fig. Extended Data 3.**
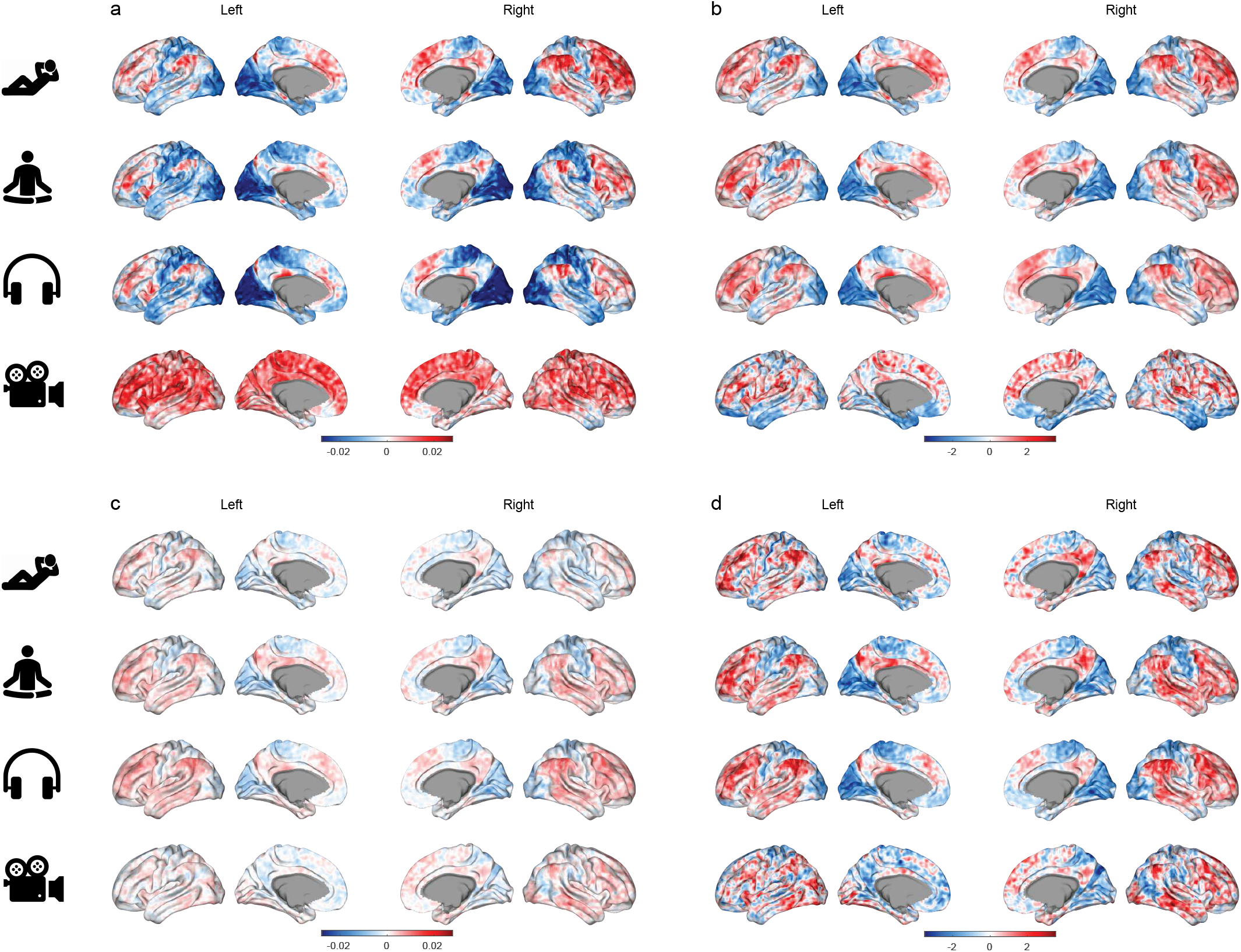
a) Global functional connectivity (GFC) difference between the psilocybin and the no-psilocybin sessions. b) The same GFC difference plotted after z-scoring. Note how z-scoring confounds the interpretation by rendering movie-watching similar to the three eyes-closed conditions. This is because positive values below the average became negative after z-scoring. c) GFC difference after global signal regression (GSR). Note the smaller magnitude of the differences compared to panel a. In the Methods section, we argue that GSR complicates the interpretation of the GFC, confounding the intended goal of quantifying “the extent to which a cortical vertex is positively/negatively correlated with the other vertices, on average” with “the extent to which a vertex is positively/negatively correlated with vertices having large standard deviation, on average”. d) GFC difference after both GSR and z-scoring. In addition to the previous confounds, z-scoring also hides the fact that GSR significantly reduces the magnitude of GFC values.

**Fig. Extended Data 4.**
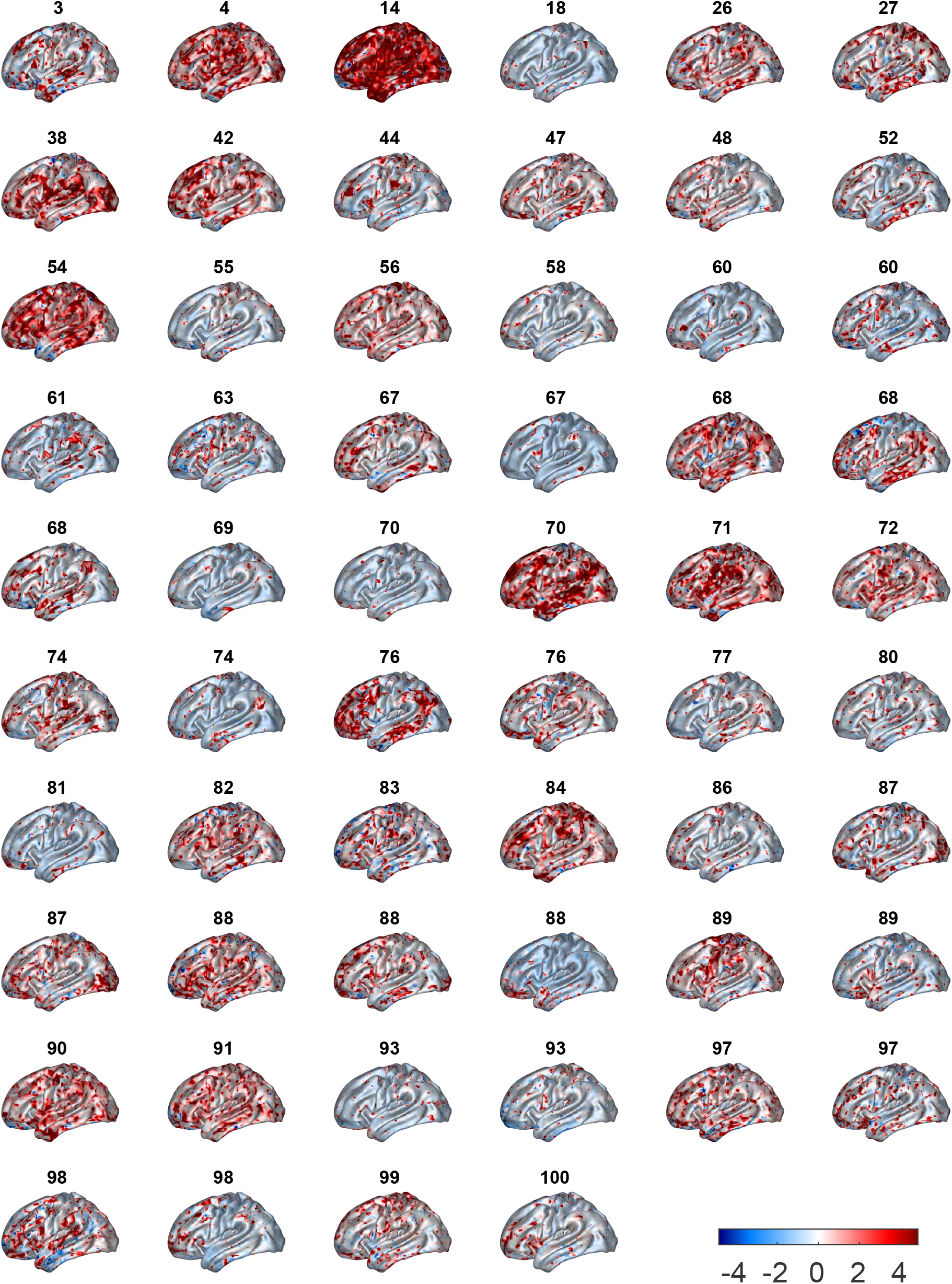
Despite the statistically significant group differences, GFC is highly variable across subjects, even when the subjective effects measured via the Mystical Experience Questionnaire (MEQ30) are similar. Here, the subjects are sorted by MEQ30 values and the individual GFC percentage difference maps (psilocybin minus baseline, divided by baseline) are shown for the resting-state condition.

**Fig. Extended Data 5.**
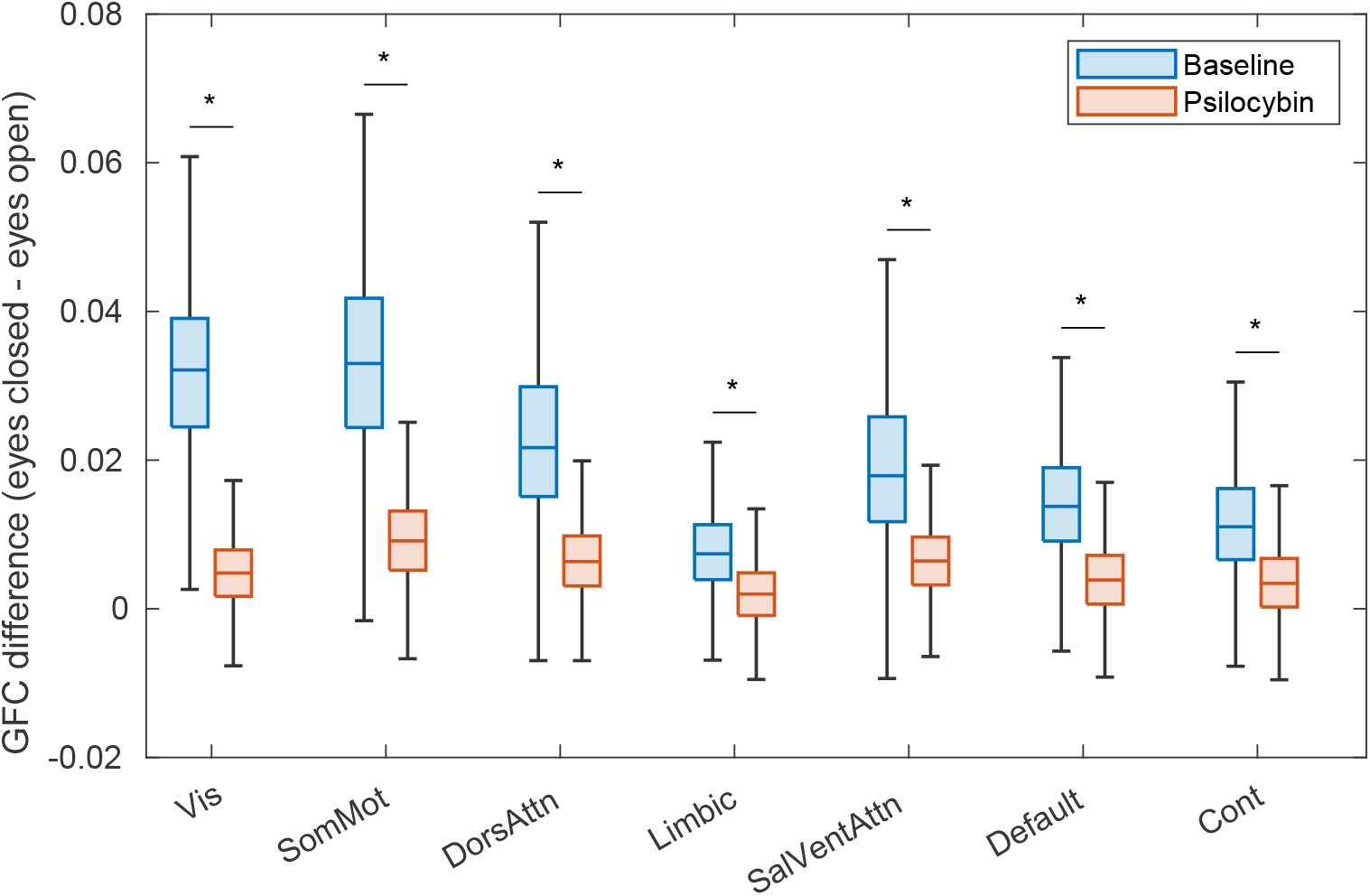
The GFC difference between eyes-closed and eyes-open conditions decreased after psilocybin administration. The largest effect was observed in the visual network (−85%, with a Cohen’s *d* effect size of −3.23), but the reduction was larger than −66% and statistically significant in all Schaefer networks (Mann-Whitney U-test right-tailed p-values *p* < 10^−307^).

**Fig. Extended Data 6.**
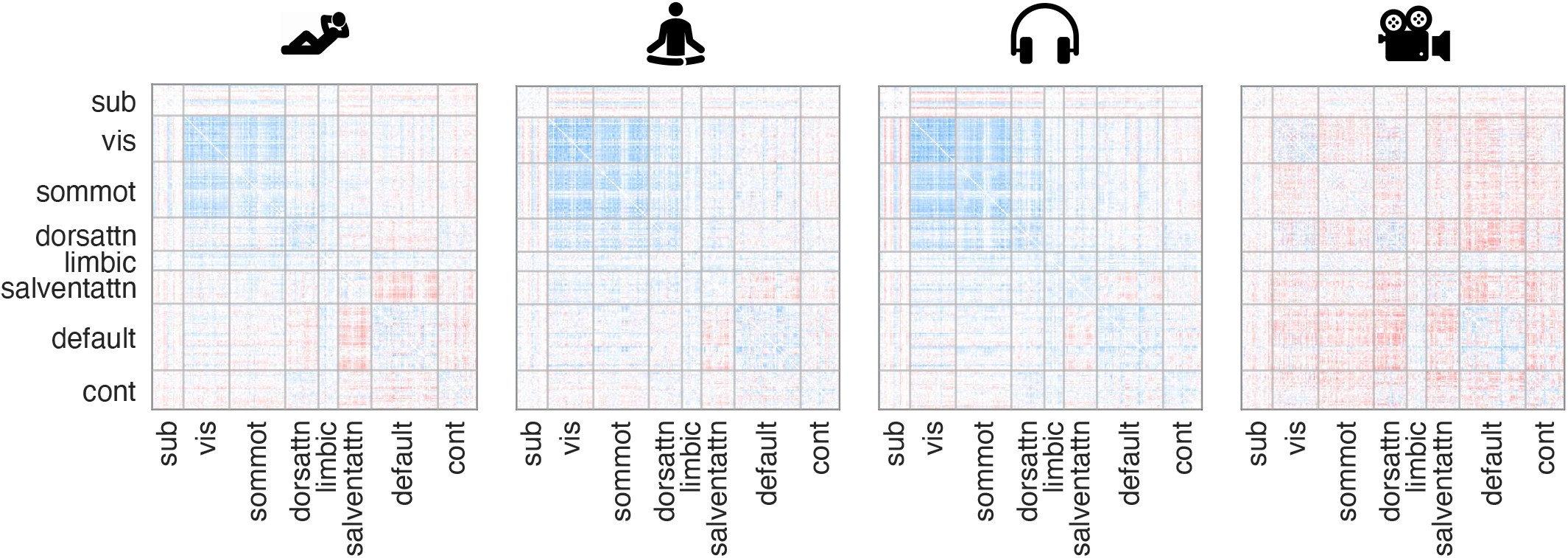
Enlarged version of Fig. 2b. The network names are: subcortex (sub), visual (vis), somato-motor (sommot), dorsal-ventral attention (dorsventattn), limbic, salience-ventral attention (salventattn), default mode, and control (cont).

**Fig. Extended Data 7.**
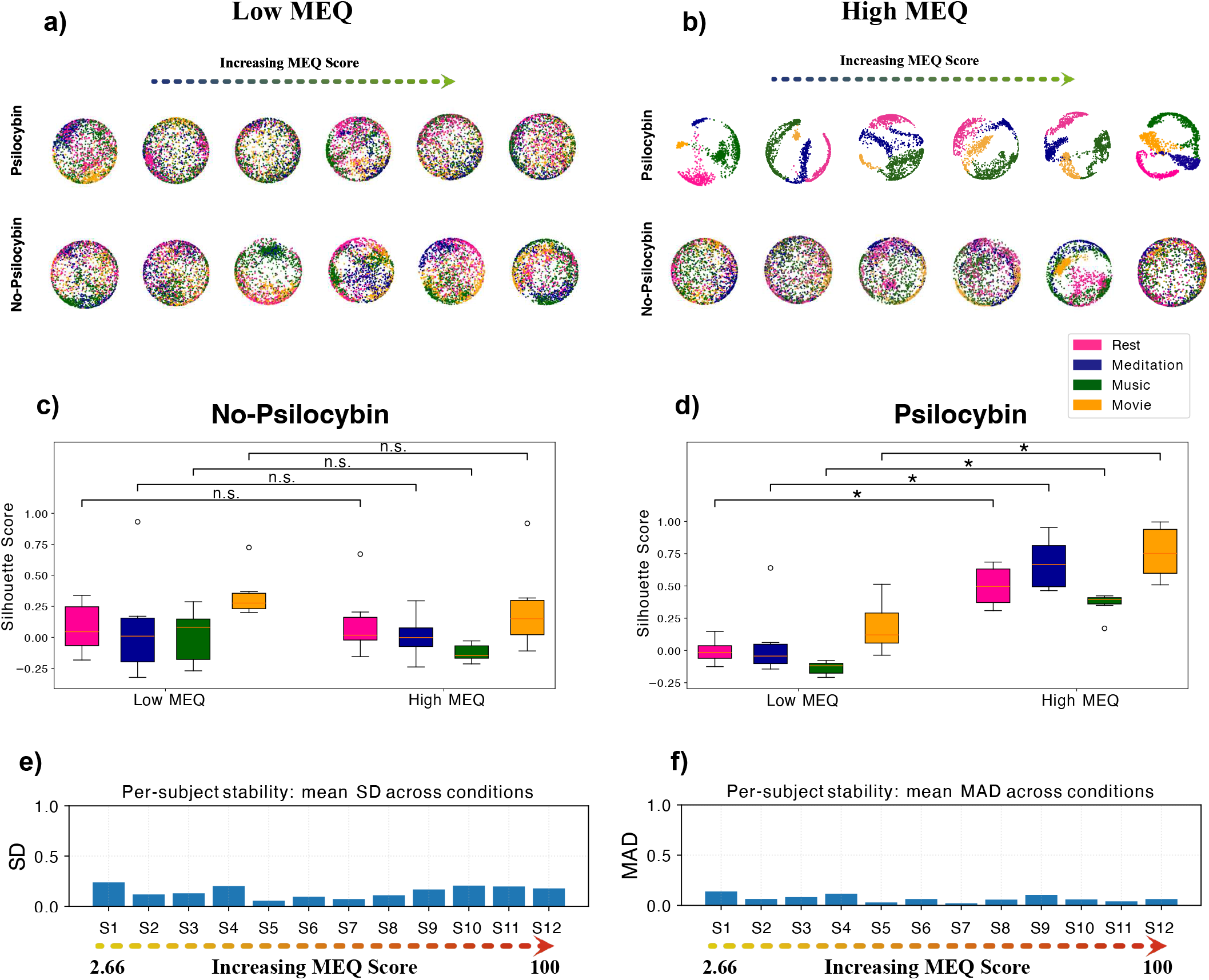
Comparison of CEBRA embeddings and silhouette scores for subjects with low vs. high MEQ scores. (a) Embeddings of *n* = 6 subjects with *low MEQ sorted by low classification accuracy* under Psilocybin and No-Psilocybin conditions. (b) Embeddings of six subjects with *high MEQ sorted by high classification accuracy* under Psilocybin and No-Psilocybin conditions. (c) Box plots of silhouette scores for the same participants during the No-Psilocybin condition, showing no significant (n.s.) differences between low and high MEQ groups (all *p* > 0.19, Welch’s two-sample t-test). For each participant, CEBRA training was repeated 10 times. (d) Box plots of silhouette scores for the same participants during the Psilocybin condition, revealing significantly higher clustering quality in high MEQ subjects compared to low MEQ subjects (Rest Δ*s* = 0.512, *p* = 0.0002; Meditation Δ*s* = 0.710, *p* = 0.0023; Music Δ*s* = 0.515, *p* < 0.0001; Movie Δ*s* = 0.631, *p* = 0.0007, Welch’s two-sample t-test). For each participant, CEBRA training was repeated 10 times. Run-to-run variability of silhouette scores was quantified using (e) the mean standard deviation (SD) and (f) the mean median absolute deviation (MAD) across the four conditions. Both SD and MAD values were consistently small (SD: ~0.05–0.25; MAD: ~0.03–0.14), indicating that repeated runs of CEBRA yield nearly identical clustering quality. Together, these results indicate that context-organised brain embeddings scale with subjective effects.

**Fig. Extended Data 8.**
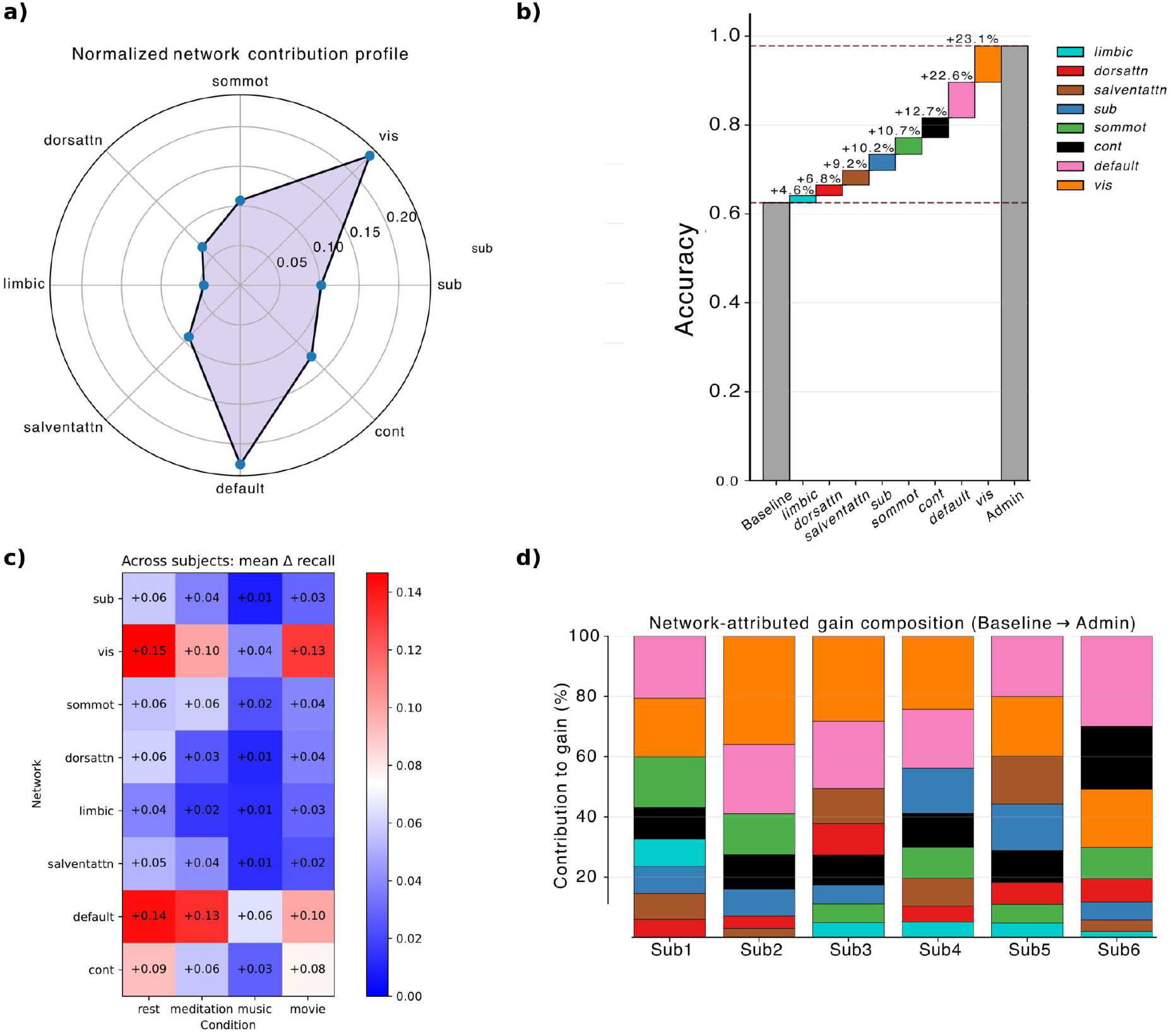
Network-attributed decomposition of context-aligned trajectory organisation under psilocybin, for the six high-MEQ participants shown in Fig. Extended Data 7 b,d. For each participant, the psilocybin time series of a given functional network was replaced with that participant’s corresponding baseline (no-psilocybin) dynamics, one network at a time. The CEBRA embedding and SVM classification were re-run on the resulting hybrid time series, and the drop in performance was measured (see Methods: Network perturbation analysis; Supplementary Information for the full attribution formula). (a) Normalised network contribution profile averaged across participants. Each axis represents one functional network; distance from centre indicates the relative contribution to the psilocybin-induced accuracy gain. Default Mode and Visual networks show the largest contributions. (b) Waterfall plot of the average additive decomposition across participants. Starting from baseline classification accuracy, each coloured segment shows one network’s attributed contribution, ordered by magnitude. The final bar corresponds to the observed psilocybin accuracy. Dashed lines mark baseline and psilocybin performance levels. Contributions are normalised to sum to the total accuracy gain (see Supplementary Information). (c) Per-condition decomposition of network contributions. Heatmap shows the across-subject mean change in per-condition recall (Δ recall = recall_psilocybin_ − recall_substituted_) for each network (rows) and condition (columns), where recall is the diagonal of the row-normalised confusion matrix. Positive values indicate that replacing that network’s psilocybin activity with baseline activity reduces correct classification of that condition, consistent with that network carrying condition-discriminative information under psilocybin. Default Mode and Visual networks show the largest effects across all four conditions, with graded modulation across contexts rather than condition-specific dissociation (see Supplementary Information for condition-level interpretation). Values are annotated within each cell. (d) Per-subject stacked bars showing the relative contribution (%) of each network to the total psilocybin-induced accuracy gain for each participant individually. Contributions are normalised so that the sum across networks equals 100% of each participant’s observed gain. The consistency of Default Mode and Visual dominance across all six individually trained models (control marginally exceeded visual in one participant) indicates that the anatomical anchoring is a stable property of the data. Network abbreviations: sub, subcortical; vis, visual; sommot, somatomotor; dorsattn, dorsal attention; limbic; salventattn, salience/ventral attention; default, default mode; cont, frontoparietal control.

**Fig. Extended Data 9.**
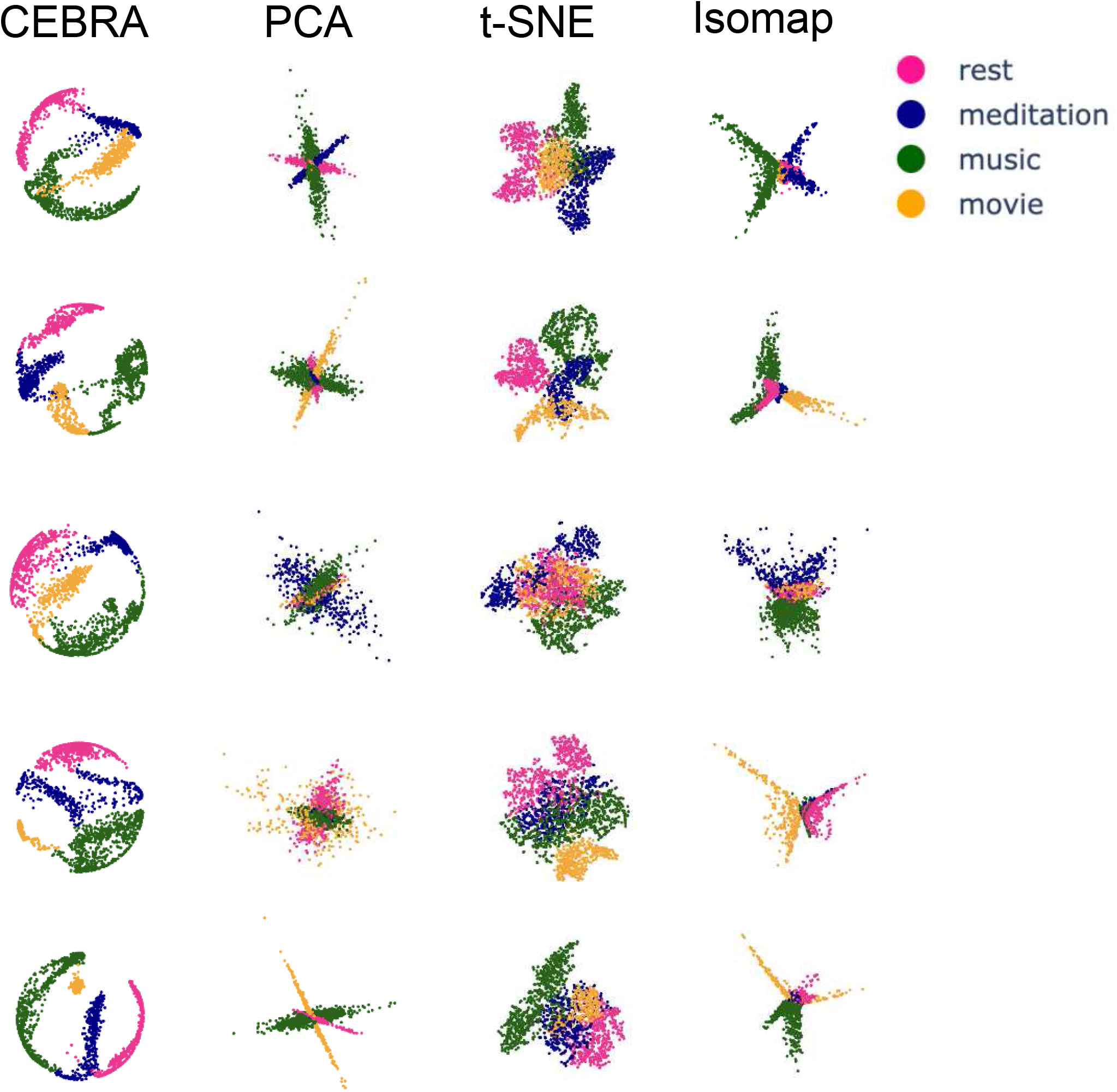
Comparison of low-dimensional embeddings of psilocybin fMRI data from high-MEQ participants using four dimensionality reduction methods: CEBRA, PCA, t-SNE, and Isomap. Each dot corresponds to a time point representing the activity of our 332 ROIs, coloured by condition (rest, meditation, music, movie). PCA shows limited separation due to its linear nature, while t-SNE and Isomap achieve clearer separation but ignore temporal continuity, leading to fragmented embeddings. CEBRA preserves temporal adjacency, producing smooth trajectories that reveal both separation between conditions and natural transitions across brain states. Despite methodological differences, embedding structure shows a generalisable pattern across dimensionality reduction approaches.

**Fig. Extended Data 10.**
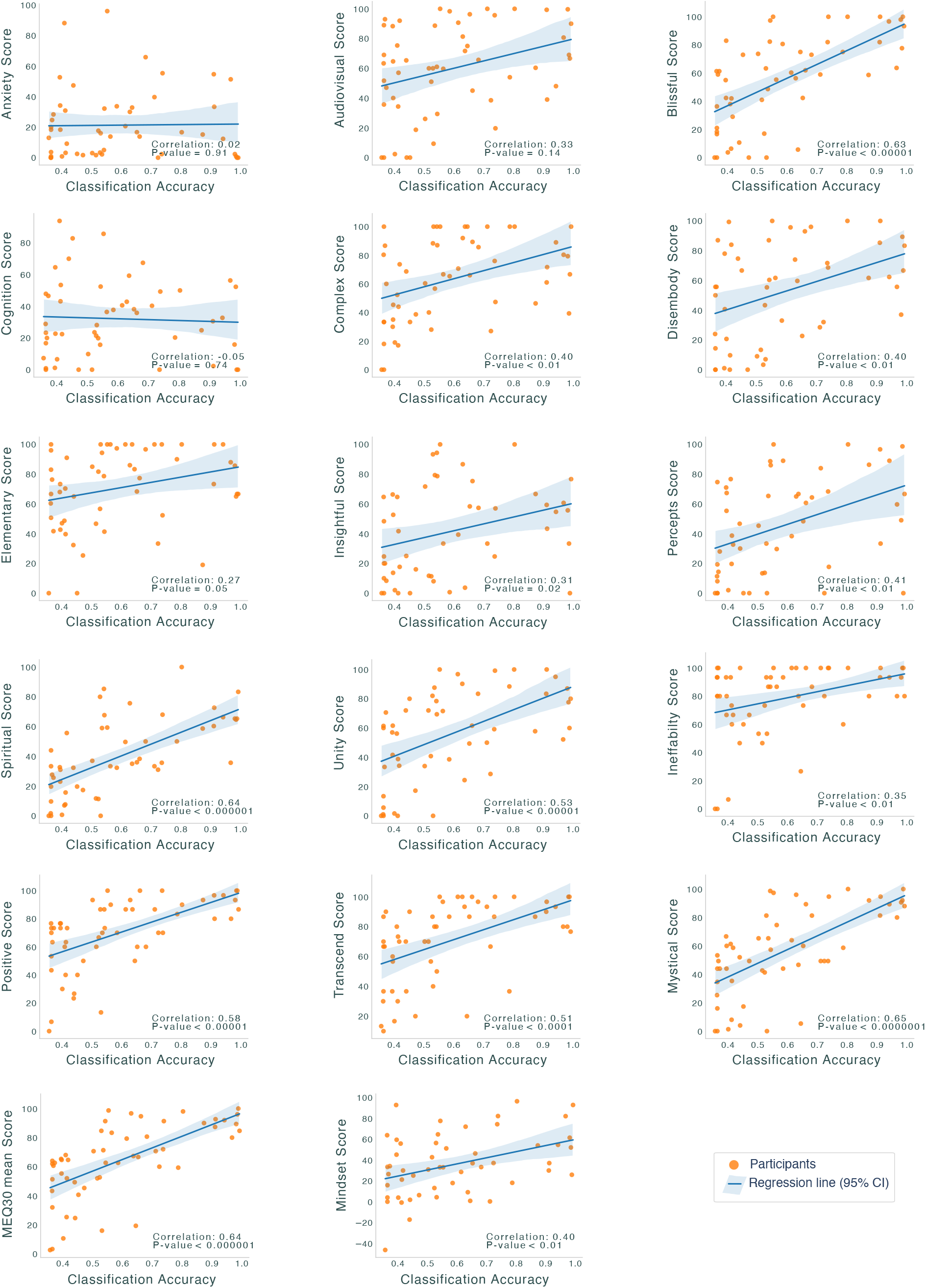
For each participant’s CEBRA trajectory, a Support Vector Machine (SVM) was used to classify the correct condition label at each time point (*n* = 54). The figure also presents the Pearson correlation between classification accuracy and participants’ 11D ASC, Mindset, MEQ30, and MEQ30 mean scores, reporting the correlation coefficient and statistical significance level (p-value ≤ 0.05, uncorrected). The blue solid line represents the regression fit, with the shaded region indicating the 95% confidence interval.

**Fig. Extended Data 11.**
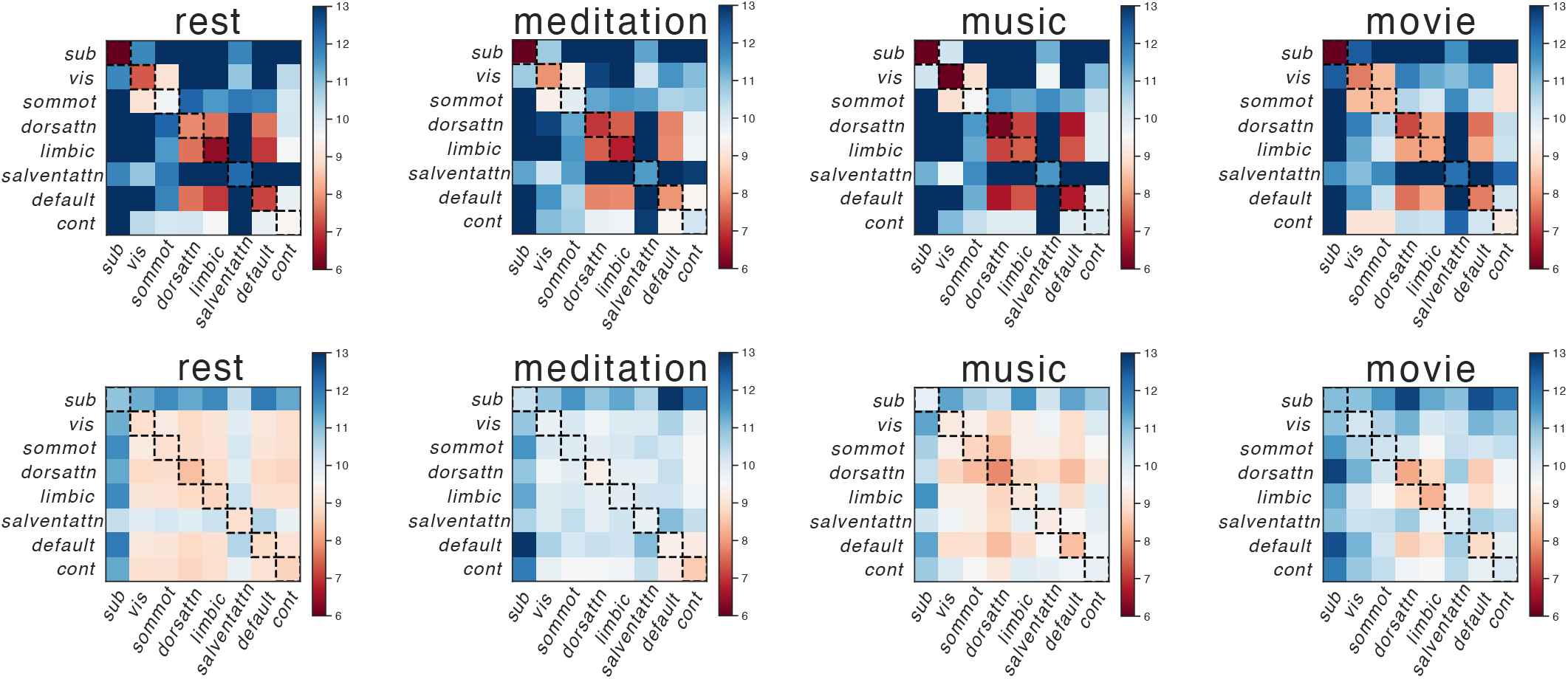
Heatmaps showing the average Euclidean distance between pairs of ROIs within each brain network in the TAVRNN embedding space across the four cognitive conditions: rest, meditation, music, and movie. The top row represents the heatmaps for participants with the top 5 MEQ scores, and the bottom row represents the heatmaps for participants with the lowest 5 MEQ scores. For all conditions, the diagonal of the top row exhibits lower values, indicating more densely clustered embeddings (ROIs positioned closer to each other) within the same network compared to the bottom row, which suggests a greater spread of ROIs in participants with lower MEQ scores. This highlights that network nodes are more closely clustered in the embedding space for participants with high reported subjective effects and this pattern is observed across all conditions, indicating a generalised effect of psilocybin on network compactness.

**Fig. Extended Data 12.**
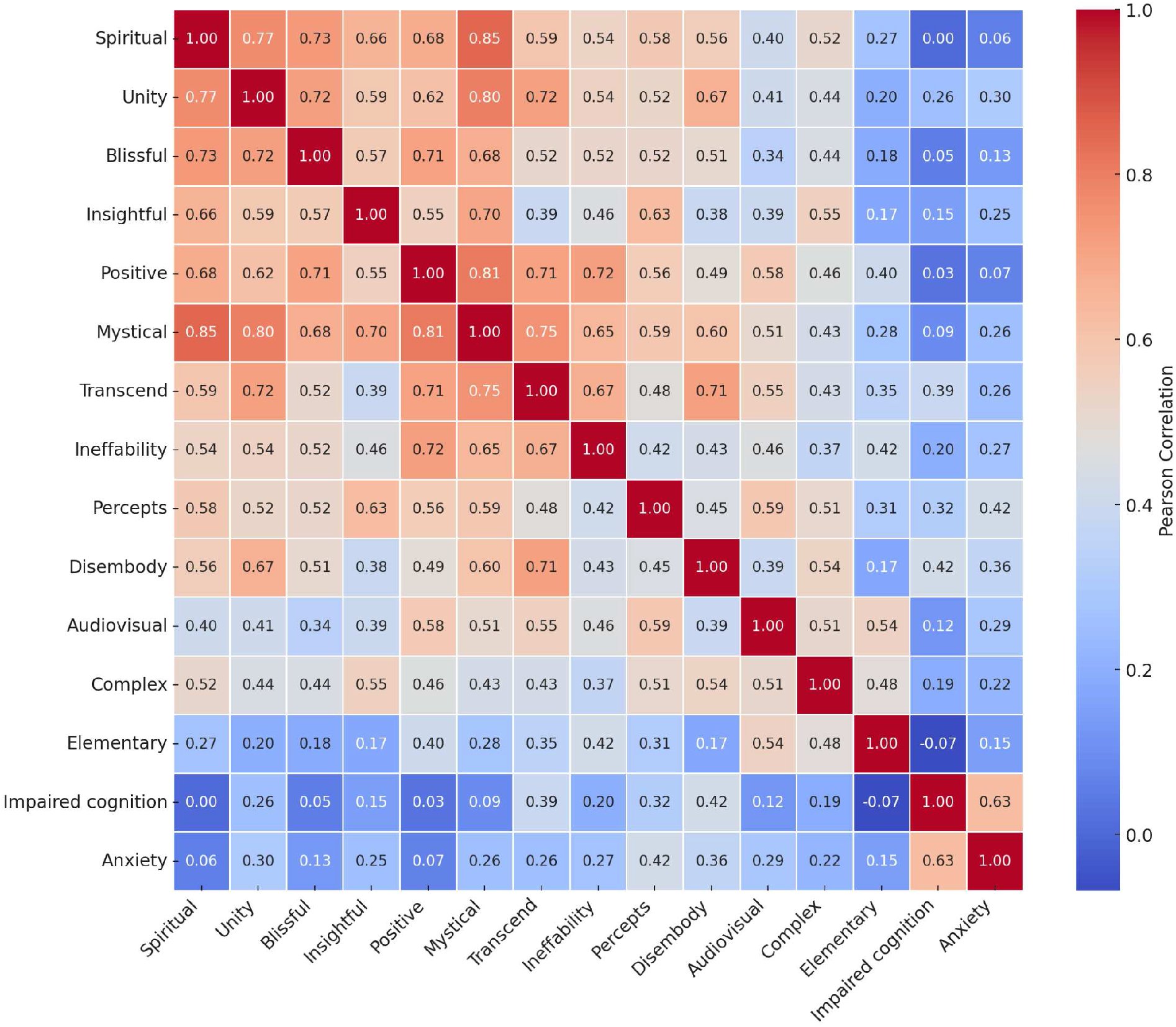
Matrix of correlation between MEQ and 11D ASC subscales, derived from our samples scores.

**Fig. Extended Data 13.**
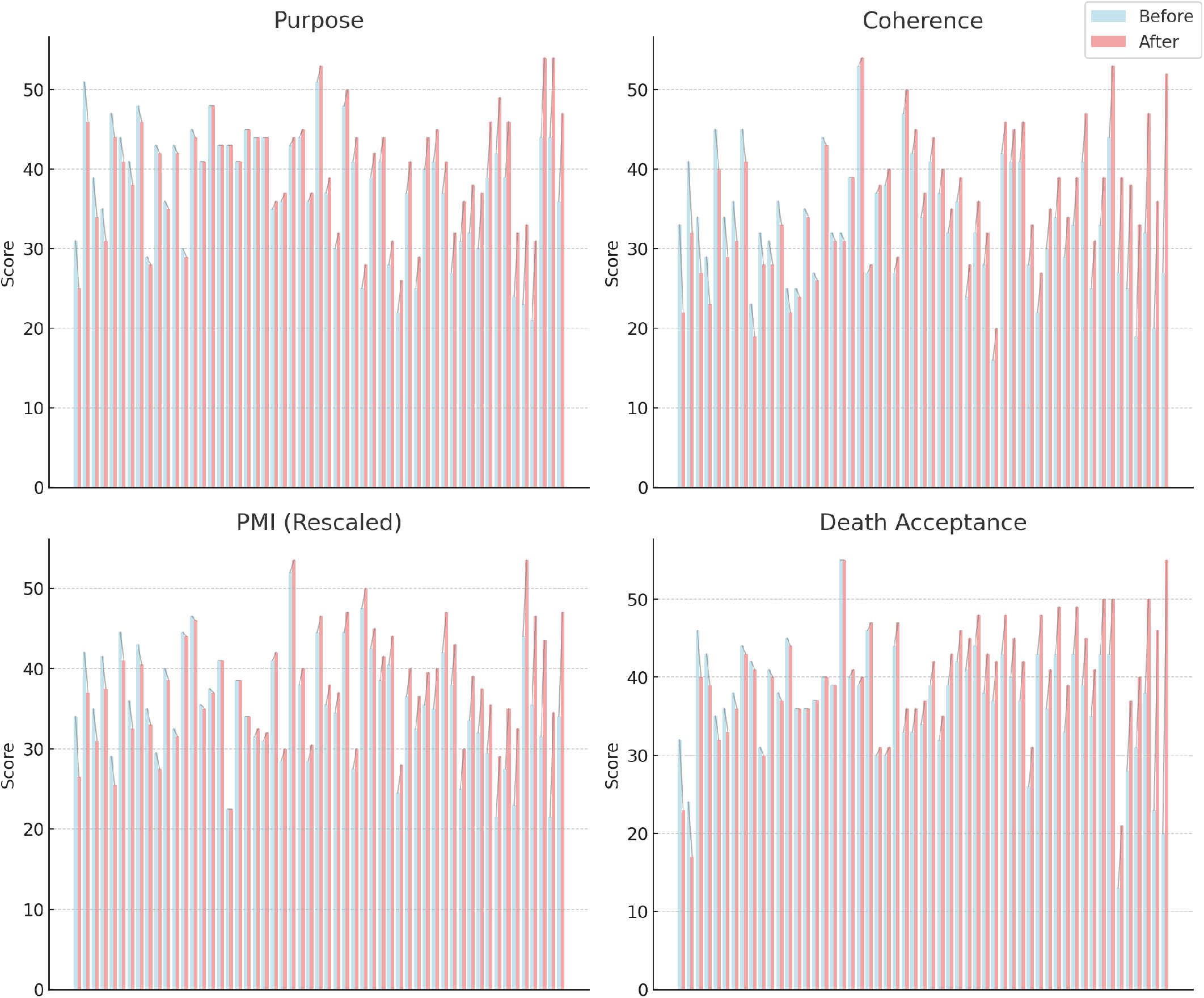
Individual Variability in Changes in Life Attitudes (LAP-R) dimensions Before and One Month After Psilocybin. Each panel displays individual participant changes for Purpose, Coherence, Personal Meaning Index (PMI) (Purpose + Coherence), and Death Acceptance. Participants are sorted by change magnitude, from largest decrease to largest increase. Each participant has two bars: blue (before psilocybin) and red (after psilocybin), with gray linking lines indicating the change for each individual. Personal Meaning was rescaled by dividing the score by 2. At the group level, paired t-tests revealed statistically significant improvements: Purpose: t(54) = −3.87, p = 0.0003, d = 0.52 (moderate effect). Coherence: t(54) = −2.81, p = 0.0069, d = 0.38 (small-to-moderate effect). Personal Meaning (Purpose + Coherence): t(54) = −3.77, p = 0.0004, d = 0.51 (moderate effect). Death Acceptance: t(54) = −3.66, p = 0.0006, d = 0.49 (moderate effect). These results indicate group-level increases in all four dimensions of interest post-psilocybin, while also highlighting substantial individual variability in response patterns. Some participants showed large increases, while others exhibited minimal or negative changes. The linking lines emphasize heterogeneity in individual responses, demonstrating that while the overall effect is positive, not all participants experience the same degree of change.

**Fig. Extended Data 14.**
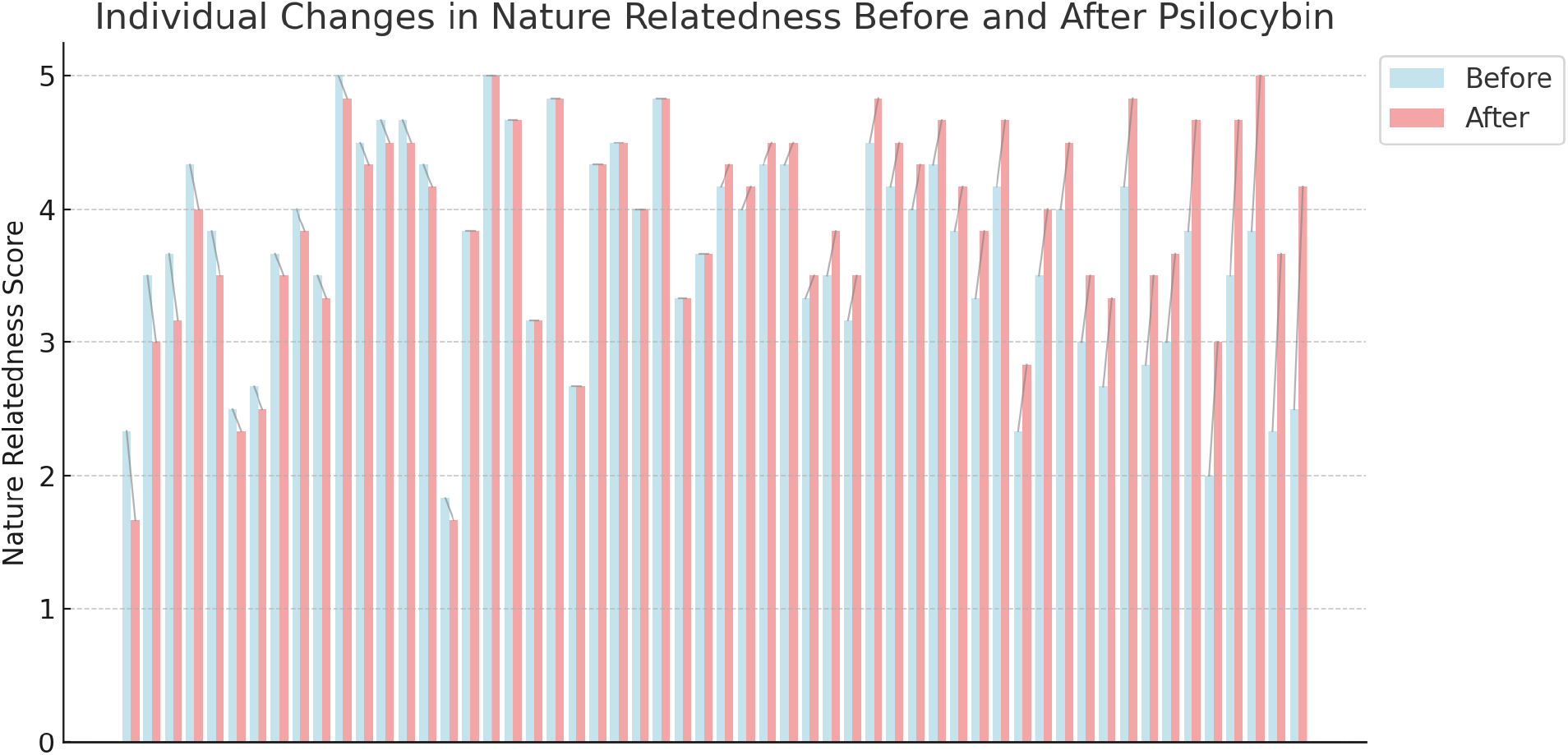
Individual Variability in Changes in Nature Relatedness Before and One Month After Psilocybin. Participants are sorted by change magnitude, from largest decrease to largest increase. Each participant has two bars: blue (before psilocybin) and red (after psilocybin), with gray linking lines indicating the change for each individual. At the group level, paired t-tests revealed statistically significant improvements t(55) = −3.37, p = 0.0014, d = 0.26 (moderate effect).

**Fig. Extended Data 15.**
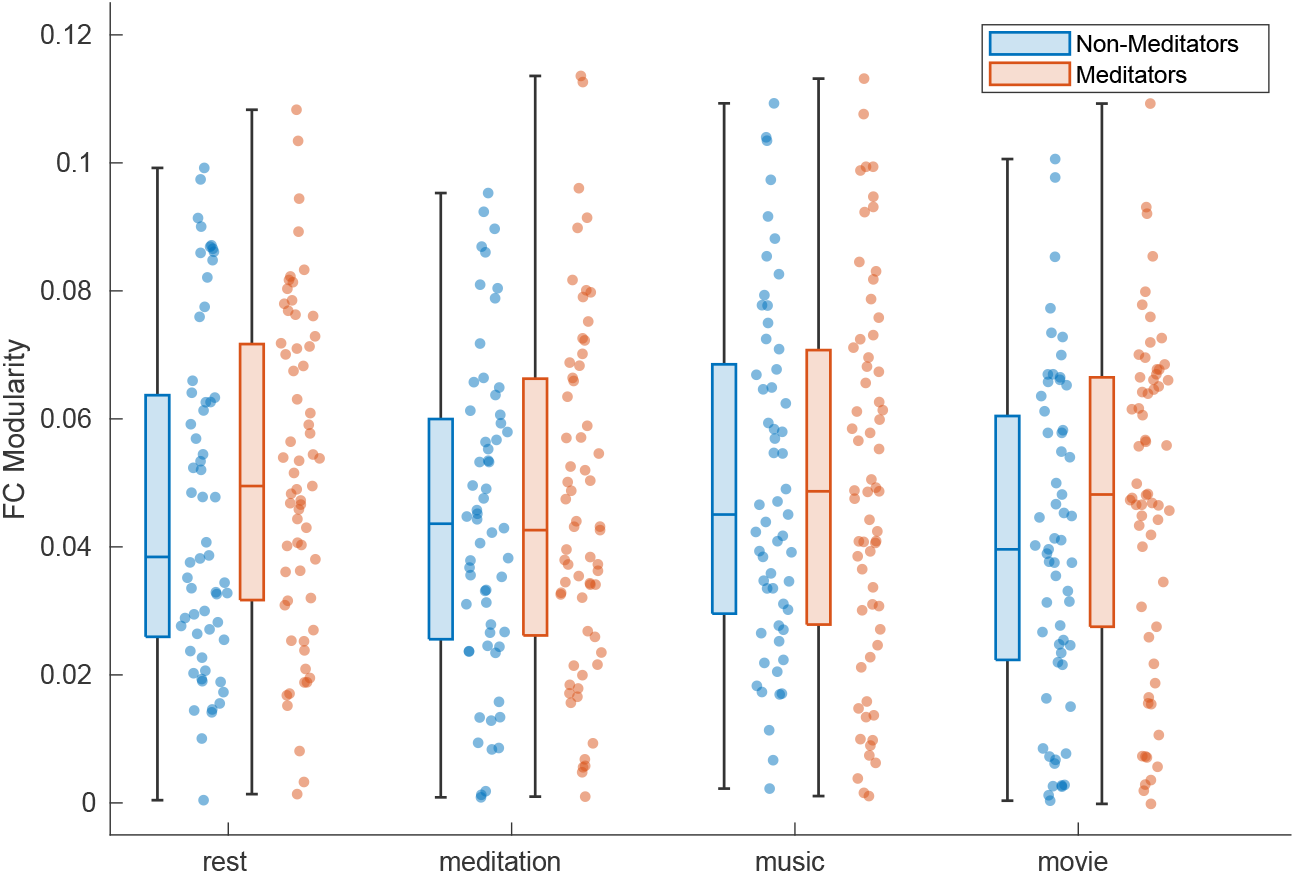
Mindfulness meditation training did not result in statistically significant changes in functional modularity (Mann-Whitney U-test, two-sided, all *p* > 0.05).

**Fig. Extended Data 16.**
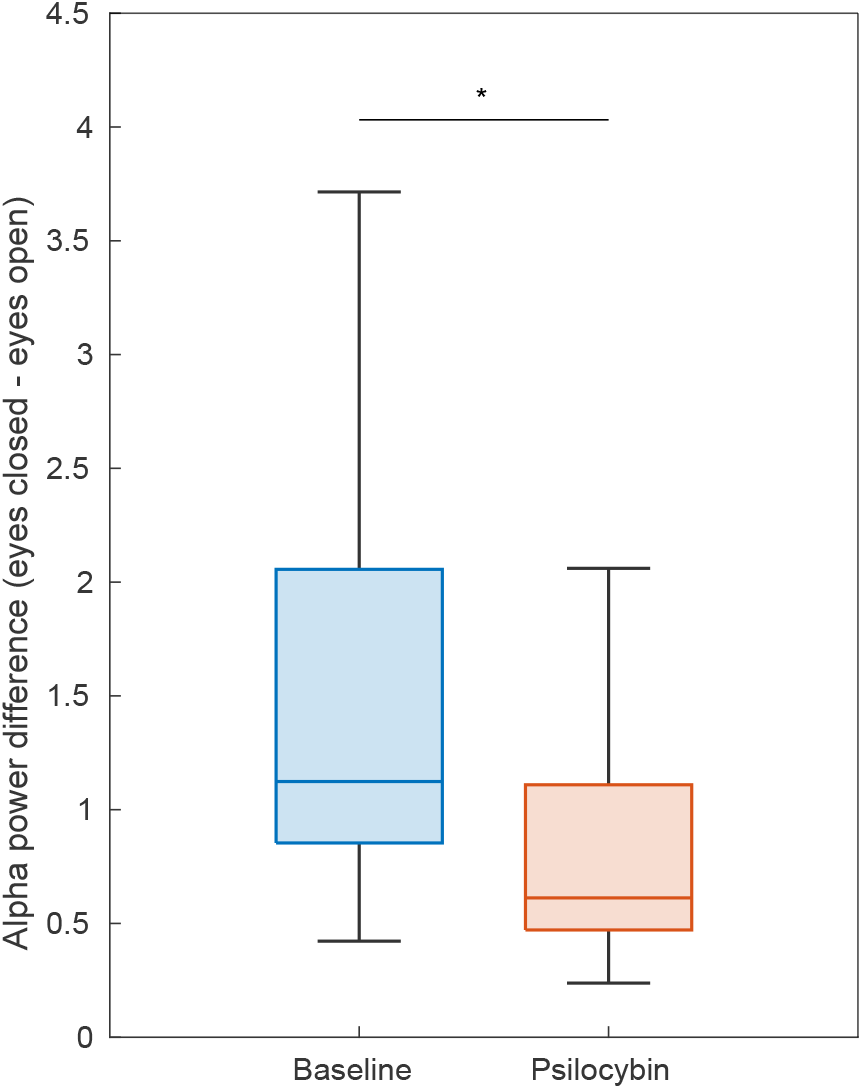
The alpha power difference between eyes-closed and eyes-open conditions decreased by 48% after psilocybin administration (Cohen’s *d* = −0.79, Mann-Whitney U-test right-tailed p-value < 10^−19^).

**Fig. Extended Data 17.**
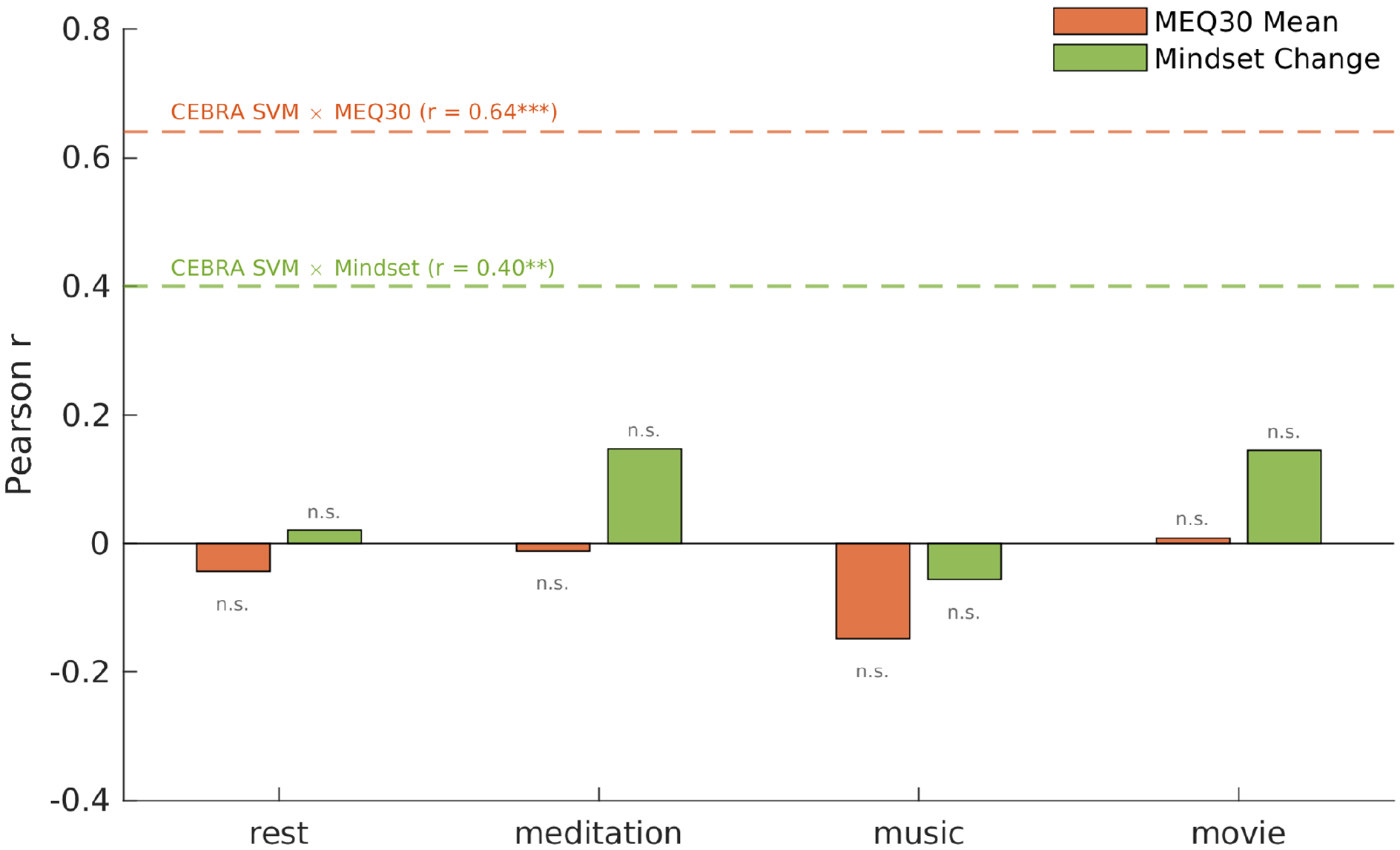
Pearson correlation coefficients between per-subject functional modularity (psilocybin session) and two behavioural outcomes—MEQ30 mean (*n* = 54) and next-day mindset change (*n* = 53)—across four conditions (rest, meditation, music, movie). Functional modularity decreased significantly at the group level under psilocybin in all conditions (Cohen’s *d* = −0.60 to −0.91; Fig. 2f), yet did not predict individual differences in either outcome (all |*r*| < 0.15, all *p* > 0.28 uncorrected; two-tailed Pearson correlations, Bonferroni-corrected across 8 tests, *α* = 0.0063; confirmed with Spearman rank correlations). Dashed lines indicate CEBRA SVM classification accuracy correlations with the same outcomes in the same participants (*r* = 0.64, *p* < 10^−6^ for MEQ30 mean; *r* = 0.40, *p* < 0.01 for mindset change). Significance annotations: ^∗∗∗^*p* < 0.001; ^∗∗^*p* < 0.01; n.s., not significant (two-tailed Pearson). Functional modularity does not predict behavioural outcomes that CEBRA classification accuracy predicts. For direct comparison of correlation magnitudes, and per-condition scatter plots (all n.s.), see Supplementary Information.

