## Supplementary_Information for "Psychedelics Align Brain Activity with Context"

*Devon Stoliker[1], Leonardo Novelli[1], Moein Khajehnejad[1], Mana Biabani[1], Matthew D. Greaves[1], Tamrin Barta[1], Martin Williams[2], Sidhant Chopra[3,4], Olivier Bazin[5], Otto Simonsson[6], Richard Chambers[7], Frederick Barrett[8], Gustavo Deco[9], Katrin H. Preller[10], Robin Carhart-Harris[11,12], Anil Seth[13,14], Suresh Sundram[15], Gary F. Egan[1,16], Adeel Razi[1,16, 17,18]*

[1]Turner Institute for Brain and Mental Health, School of Psychological Sciences, Monash University, Australia.
[2]School of Health Sciences, Swinburne University of Technology, Hawthorn, VIC 3122, Australia.
[3]Orygen, The University of Melbourne, Parkville, Australia.
[4]Centre for Youth Mental Health, The University of Melbourne, Australia.
[5]British Association of Mindfulness-based Approaches (BAMBA) Listed Teacher, Oxford, UK. [6]Department of Neurobiology, Care Sciences and Society, Karolinska Institute, Sweden.
[7]Monash Centre for Consciousness and Contemplative Studies, Monash University, Melbourne, Australia.
[8]Department of Psychiatry and Behavioral Sciences, Center for Psychedelic and Consciousness Research, Johns Hopkins University School of Medicine, Baltimore, MD, USA.
[9]Center for Brain and Cognition, Theoretical and Computational Group, Universitat Pompeu Fabra/ICREA, Barcelona, Spain.
[10]Department of Adult Psychiatry and Psychotherapy, Psychiatric University Clinic Zurich and University of Zurich, Switzerland.
[11]Centre for Psychedelic Research, Department of Brain Sciences, Imperial College, London, London, UK.
[12]Psychedelics Division, Neuroscape, University of California, San Francisco, USA.
[13]Sussex Centre for Consciousness Science, Department of Informatics, University of Sussex, Brighton, UK.
[14]Program on Brain, Mind, and Consciousness, Canadian Institute for Advanced Research, Toronto, Canada.
[15]Department of Psychiatry, School of Clinical Sciences, Monash University, Clayton, VIC 3168, Australia.
[16]Monash Biomedical Imaging, Monash University, Australia.
[17]Wellcome Centre for Human Neuroimaging, University College London, United Kingdom.
[18]CIFAR Azrieli Global Scholars Program, Toronto, Canada.

### Mindset Measure

We used a custom-made, unvalidated Likert scale to measure mindset (psychological) change the day after psilocybin, measuring 11 qualities on a scale from -100 to +100. The average scores across these 11 qualities were used a mindset score. The term *mindset* here denotes a composite of perceived psychological change across these dimensions.

**Survey Question:**
*" Please remark on the mindset qualities that you notice have changed since the psilocybin experience. Please note: The scale of this measure is -100 to +100. Any score below 0 means you experienced a negative change for the item after psilocybin. A middle score of 0 means no change. Any score above 0 means a positive change for the item after psilocybin."*

| Quality | Group Ave. Score (Scale -100 to +100) |
|---|---|
| **Patience** | +36.60 |
| **Creativity** | +21.65 |
| **Sense of meaningfulness in life** | +28.42 |
| **Sense of harmony** | +34.20 |
| **Sense of connectedness with yourself** | +41.34 |
| **Sense of connectedness with others** | +31.64 |
| **Sense of connectedness with nature** | +27.48 |
| **Sense of openness to new experiences** | +45.14 |
| **Sense of acceptance** | +44.39 |
| **Sense of inner peace** | +46.18 |
| **Sense of imagination** | +31.31 |

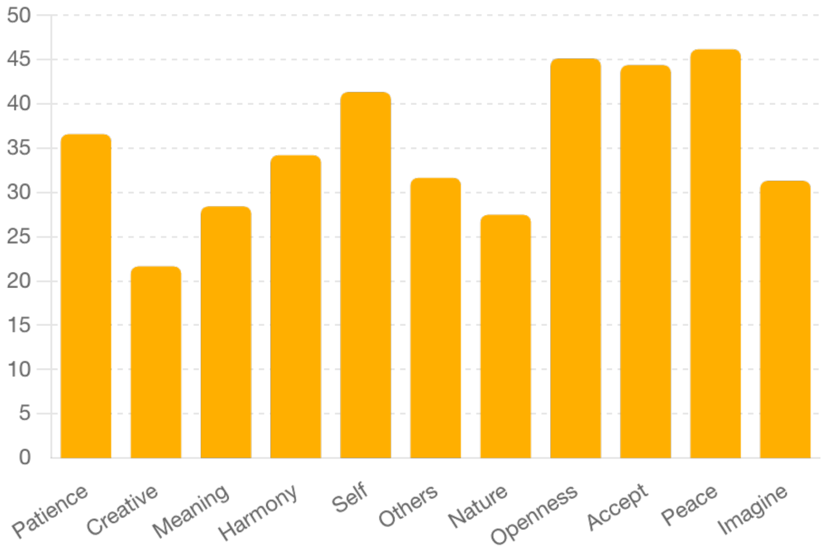

**Average Mindset Scores Across Different Dimensions**
This bar chart displays the average scores for various mindset dimensions, including Patience, Creativity, Meaning, Harmony, Self, Others, Nature, Openness, Acceptance, Peace, and Imagination. The highest scores are observed in Peace, Openness, and Acceptance, while Creativity and Nature have the lowest averages.

---

### Day-After Qualitative Self-Reports on Psilocybin Experiences

The following tables are drawn from semi-structured self-report questionnaires collected from participants the day after psilocybin. Key excerpts from participant reports are structured by the specific questions under which they were collected. Responses that appear in the main text are marked with an asterisk (*). Column headings and verbatim questions are listed above each table in brackets.

#### Emotional Responses During MRI (EXP.MRIEMOTIONS – "During the MRI my emotions were…")

| Theme | Raw Participant Report |
|---|---|
| Wave-like Sensory Immersion in MRI* | *"Honestly I can't even explain. I remember preparing and [names omitted] organising me for the MRI and lying down, then within minutes the sounds of the machine were creating a wave of music that was so beautiful, so intense. It was building from a couple of sounds, then louder and louder until it was an inevitable wave of sound that crashed through me. I don't know how to describe it other than to say I lost all sense of self, and I became at one with all my surroundings."* |
| Melding with the Environment (MRI)* | *"I melded with the MRI machine, floor, walls, air. It was like nothing existed except for me, or me as part of something bigger."* |
| Loss of Self & Integration into Surroundings (MRI)* | *"I lost all sense of self, and I became at one with all my surroundings. I melted into everything else around me and became part of a larger, bigger thing. I saw myself as part of a bigger network of things."* |

#### Visual and Perceptual Alterations (EXP.VISUALPT1, EXP.VISUALPT2 – "Did you have an altered visual experience? If yes, please describe your visual experience and note where you were.")

| Theme | Raw Participant Report |
|---|---|
| Seamless Transitions Between Eyes-Closed & Eyes-Open | *"Switching from eyes closed to eyes open and vice versa... visual experience with eyes closed was of being in an open 'sky' area... merging seamlessly into the EEG environment when opening my eyes."* |

| | |
|---|---|
| Perceptual Expansion | *"I saw my inner monologue (maybe all of language?) as a floating double helix that I could bend at will."* |

### Music, Synaesthesia, and Multisensory Integration (EXP.MUSIC, EXP.SYNAESTHESIA – "Did you experience synaesthesia? If yes, please describe your experience.")

| Theme | Raw Participant Report |
|---|---|
| Music as a Guide to the Experience | *"Became the landscape of my experience - I stopped being able to distinguish between myself, the music, the evolving patterns."* |
| Auditory-Induced Imagery | *"The music influenced what I saw. It wasn't necessarily like a visual representation of the music, but it was more like the music was the landscape."* |
| Rhythmic Synchronisation | *"Visual hallucinations were in rhythm/sync with the music, as the music changed, so would what I was seeing."* |
| MRI-Induced Visual Patterns | *"The sound of the MRI created incredibly real visual imagery – rolling waves of gold and silver like an ocean of watered steel with fine detailed Thai motifs."* |

### Meaning, Self-Dissolution, and Psychological Insights (EXP.FINDMEANING, EXP.MOSTMEANINGPT1, EXP.INTEREST – "Did you find meaning in your experience? If yes, please describe how." / "What aspects of the experience were most interesting or important to you?")

| Theme | Raw Participant Report |
|---|---|
| Altered Sense of Self & Narrative | *"The fact that I lost the narrative of myself and ended up just kind of vibing makes me realise how much of who I think I am is just a story that I am telling myself."* |
| Life Perspective & Acceptance | *"I now see I was not in total control of my life... I now feel grateful for this life and want to cherish my time better."* |
| Reduced Attachment to Stressors | *"The meaning that I derived most from the experience was realising the little stressors in life that often keep me in my head and ruminating became insignificant and no longer held me hostage."* |
| Profound Self-Dissolution in MRI | *"Definitely the MRI because I completely lost all sense of self, of body, of being. And I legitimately felt that I existed only in consciousness."* |
| Loss of Plot and Self During EEG* | *"During the EEG, everything blended into the colours/patterns/landscape... I lost any form of language and couldn't hold a thought. I lost the plot of who I was, where I was, if I was even here, what was happening."* |
| Ego Dissolution & Profound Experience (MRI)* | *"The MRI, that experience of what I now know is ego dissolution, and not existing, is one of the most peaceful and profound things I could ever experience."* |

### Supplement to Decreased functional modularity

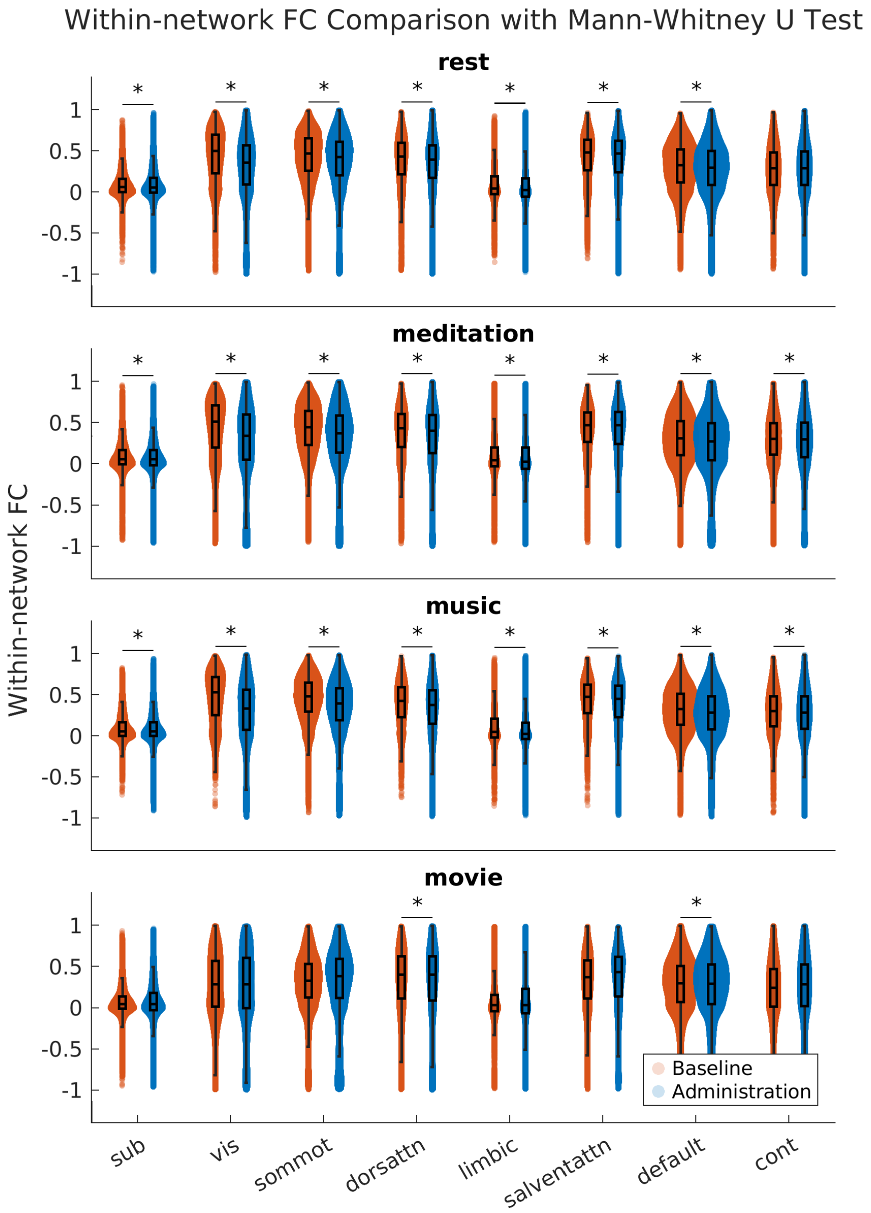


Psilocybin reduces functional connectivity within most networks during the eyes-closed conditions: resting-state, meditation, and music. Conversely, during the movie, the within-network functional connectivity only decreases in the dorsal attention network and the default-mode network (DMN).

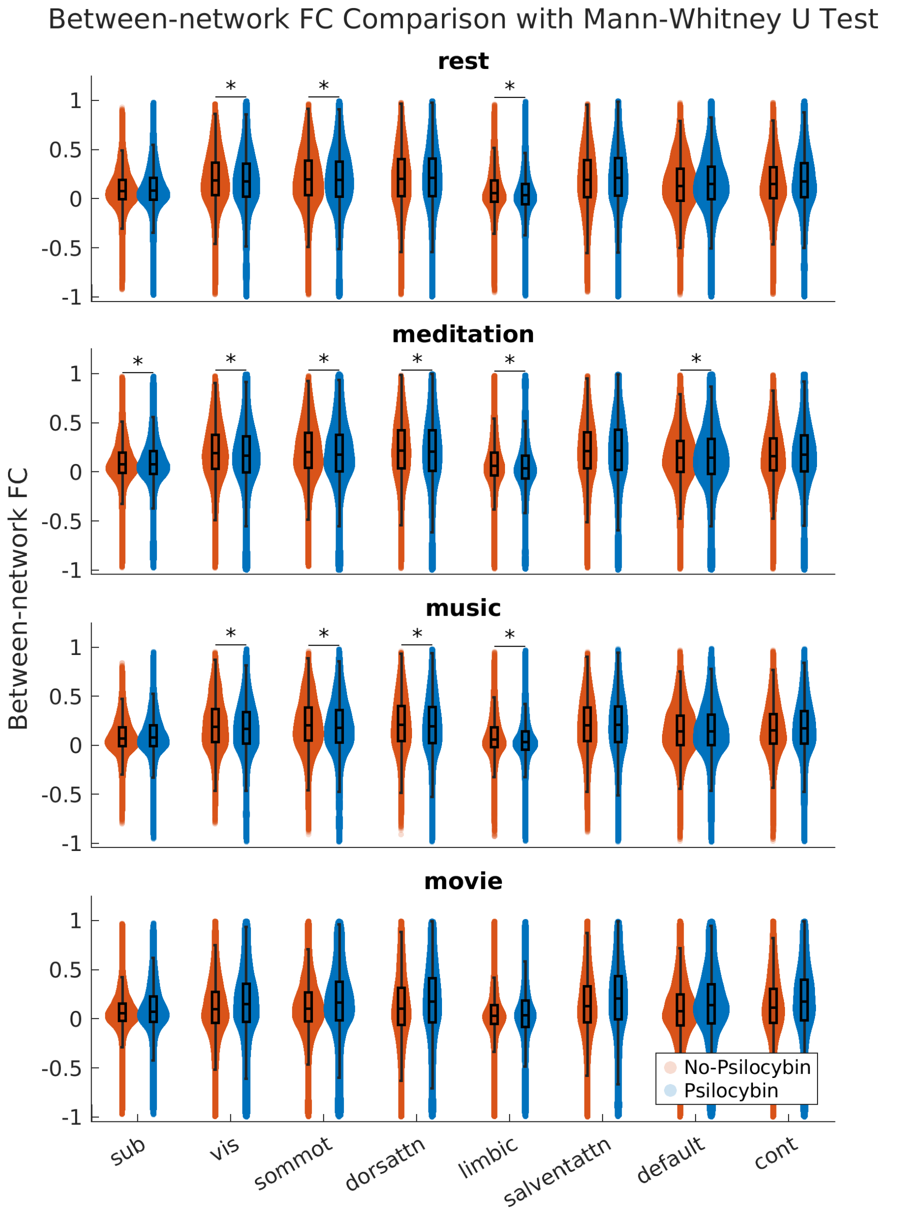


Psilocybin reduces functional connectivity between specific networks (visual, somato-motor, limbic) and the rest of the brain during the eyes-closed conditions. Conversely, during the movie, the functional connectivity increases between most networks. Each swarm chart and box plot indicates the average functional connectivity between one network and the rest of the brain. Each star symbol indicates a significant decrease based on the Mann-Whitney U-test (alpha = 0.00625 to correct for multiple comparisons across eight resting-state networks).

### Supplement to Network perturbation analysis: attribution formula

For the network perturbation analysis described in Methods, we defined a formal attribution framework to quantify each functional network's contribution to context-aligned trajectory organisation under psilocybin.

Let $s$ index subjects ($s = 1, \ldots, S$) and $k$ index functional networks ($k = 1, \ldots, K$). The total improvement in SVM classification accuracy from baseline to psilocybin is:

$$G^{(s)} = Acc^{(s)}_{admin} - Acc^{(s)}_{baseline}$$

The contribution of network $k$ is measured as the accuracy drop under baseline substitution of that network:

$$\Delta Acc^{(s)}_{k} = Acc^{(s)}_{admin} - Acc^{(s)}_{admin\ with\ k \to baseline}$$

Normalised weights, retaining only positive contributions:

$$w^{(s)}_{k} = \frac{max\left(\Delta Acc^{(s)}_{k}, 0\right)}{\sum_{j=1}^{K} max\left(\Delta Acc^{(s)}_{j}, 0\right)}$$

Each network's attributed contribution:

$$Contrib^{(s)}_{k} = w^{(s)}_{k} \times G^{(s)}$$

guaranteeing that:

$$\sum_{k=1}^{K} Contrib^{(s)}_{k} = G^{(s)}$$

For visualisation (Fig. XX), contributions were expressed as percentages of the total gain:

$$Contribution\%^{(s)}_{k} = \frac{Contrib^{(s)}_{k}}{G^{(s)}} \times 100$$

This additive decomposition ensures that each network's contribution is expressed relative to the total psilocybin-induced accuracy gain, allowing direct comparison across networks and participants. For the per-condition decomposition, recall was defined as the diagonal of the row-normalised confusion matrix for each condition, and Δ recall was computed as recall(psilocybin) − recall(substituted) for each network and condition.

### Per-condition decomposition of network contributions

The per-condition decomposition revealed that Default Mode and Visual networks showed the largest Δ recall across all four conditions, with graded modulation across contexts rather than condition-specific dissociation (Extended Data Fig. 8c).

Rest showed the strongest overall dependence on individual network substitutions (Visual: Δ recall = +0.15; Default Mode: +0.14), consistent with the absence of an external stimulus scaffold allowing internally generated dynamics, particularly within Default Mode and Visual systems, to dominate trajectory organisation.

Movie viewing showed strong Visual network contributions (Δ recall = +0.13), consistent with the dominant role of visual processing during the eyes-open condition.

Music listening consistently showed the weakest dependence on any single network's substitution (largest single contribution: Default Mode, Δ recall = +0.06), despite exhibiting strong overall clustering quality (silhouette scores: $\Delta s = 0.515$, $p < 0.0001$; Extended Data Fig. 7d). This indicates that the trajectory organisation for music is robust but draws on a more distributed set of neural resources. We note two factors that may contribute to this pattern. First, this condition involved a substantially longer, continuously evolving auditory stimulus (11:24 min vs. 6–8 min for other conditions), which is likely to increase temporal heterogeneity in BOLD trajectories under psilocybin. Second, auditory processing regions are not represented as a dedicated network in the Schaefer cortical parcellation, meaning that condition-specific contributions from auditory systems may be distributed across multiple network labels. We therefore interpret the attenuated per-network effects for music as a consequence of the richer temporal structure and parcellation coverage of this condition, consistent with context-dependent alignment mediated by a broader coalition of systems rather than a few dominant networks.

Taken together, the per-condition decomposition confirms that the anatomical anchoring identified in the aggregate analysis (Default Mode and Visual dominance) generalises across contexts, while the graded variation across conditions is consistent with context-dependent modulation of a common organisational motif.

### Screening disclosures of serotonergic psychedelic exposures

While our screening excluded any hallucinogen use within the past 6 months (to avoid subacute carryover/tolerance and expectancy reinforcement) and acted as an explicit lower bound threshold for exclusion, in practice, we were able to recruit a sample meeting our operational definition of “psychedelic-naïve”, meaning no prior psychedelic experience *with* *subjective effects*. The table above lists the 3 occurrences of nominal/remote serotonergic psychedelic exposures in our sample of 62 subjects.

| Study ID | Timing | Self-Report |
|---|---|---|
| PC-026 | 2017 | "Psilocybin, twice, microdose. No effects noticed." |
| PC-029 | 1998 | "Psilocybin, twice, unspecified dose. Minimal/No recollection of subjective effects." |
| PC-225 | > 6 months pre-screening | "Psilocybin, once, microdose. No effects noticed." |

Table of screening disclosures of nominal/remote serotonergic psychedelic exposures (n = 3; ≈4.8% of total sample)

### Expectancy considerations

At moderate–high psychedelic doses, effective blinding is difficult to achieve in practice because distinctive subjective effects make allocation easy to infer (Gasser et al., 2014; Holze et al., 2022; Holze et al., 2023; Studerus et al., 2011; Szigeti & Heifets, 2024), and suitable active placebos are lacking, with participants often discerning allocation when active placebos are used (Bogenschutz et al., 2022; Muthukumaraswamy et al., 2021). Several further features make it doubtful that expectancy accounts for our main findings: (i) large separations in reported subjective effects between active-dose and placebo arms in both oral clinical-dose and intravenous designs (Bogenschutz et al., 2022; Muthukumaraswamy et al., 2013), including 20 mg oral psilocybin (Ley et al., 2023), which is comparable to our 19 mg dose and shows very low placebo subjective scores; dose–response and time-dependent evidence across designs further corroborates pharmacological dominance over expectancy (Hirschfeld et al., 2023; Hirschfeld & Schmidt, 2021; Muthukumaraswamy et al., 2013; Preller et al., 2020); (ii) pharmacokinetic–pharmacodynamic coupling between psilocin exposure, receptor engagement, and time-varying neural dynamics (M. K. Madsen et al., 2019; Olsen et al., 2022; Preller et al., 2018; Preller et al., 2020); and (iii) condition-separated ML embeddings appear only when subjective effects are strong (high MEQ) and are absent at baseline/low-MEQ. Consistent with this, a published re-analysis of a head-to-head double-blind RCT found that pre-trial expectancy significantly predicted outcomes in the escitalopram arm but not in the psilocybin arm (Szigeti et al., 2024). Although expectancy is empirically difficult to eliminate in psychedelic trials, the subjective-response-dependent emergence of structured embeddings, their absence at baseline or with low subjective effects, and the replicated convergence between eyes-open and eyes-closed conditions across modalities despite reversed order collectively make expectancy an implausible primary driver of our main results.

### Order considerations

The fixed within-modality sequence was chosen for safety and experiential coherence. We note that: (i) the movie condition was reversed across modalities (last in fMRI, first in EEG) and separated by a 20–40 minute setup break; (ii) head-motion indices showed no monotonic rise across sessions (Extended Fig 2); (iii) inter-individual post-dose pharmacokinetic-pharmacodynamic timing variability (Brown et al., 2017; Hasler et al., 2004; Otto et al., 2025) the scan order does not correspond to a common exposure phase across participants; and (iv) each scan lasted 6–11 minutes, limiting carry-over effects. Together with the absence of sequence-aligned signatures in the ML embeddings and (Extended Fig. 7) cross-modal replication of reduced differentiation between eyes-open and eyes-closed (Fig. 2 and Fig. 6), these factors mitigate the risk that fixed order explains the main results (i.e., the ML findings and the eyes-open vs eyes-closed convergence).

Residual order influence cannot be excluded for fine-grained comparisons among the eyes-closed conditions analysed with conventional metrics. Because these scans were presented in a specific order—progressing from low- to high-stimulus (rest → meditation → music), subtle sequence effects (potentially state-dependent under psilocybin, and longer-timescale context drift) could contribute to their relative differences. Accordingly, eyes-closed comparisons are reported descriptively.

### EEG timing

Our EEG session sampled the late-peak/early-offset portion of the acute response. Human PK/PD data indicate that, after oral dosing, plasma psilocin Tmax typically occurs at ~2–3 h (≈90–150 min) with inter-individual variability, and concentrations remain elevated for several hours; subjective intensity closely tracks psilocin levels and neocortical 5-HT2A engagement across this window (Holze et al., 2022; Ley et al., 2023; Martin K. Madsen et al., 2019). Consistent with this coupling, psilocybin-induced changes in large-scale network integrity/segregation have been characterised over multiple hours post-dose, indicating robust central effects throughout our EEG interval (Madsen et al., 2021).

Evidence that EEG timing is within the acute state is demonstrated between consistency between our findings and canonical electrophysiological signatures are observed in established EEG/MEG psychedelic research---most notably broadband desynchronisation with pronounced parieto-occipital alpha reductions and increased neural signal diversity/entropy (Muthukumaraswamy et al., 2013; Schartner et al., 2017; Timmermann et al., 2023)

### Biological interpretability of the latent spaces derived from machine-learning analyses

Our machine learning analyses, while agnostic to underlying neurochemistry, allow biologically grounded interpretations of functional brain reorganisation under psilocybin. CEBRA embeddings capture temporally structured, condition-sensitive trajectories of multiregional activity within individuals, revealing neural states that scale with the quality of subjective experience. TAVRNN complements this by modelling time-evolving changes in brain network topology, capturing both increased within-network cohesion and enhanced cross-network integration across conditions.

To quantify within-network cohesion in the TAVRNN embedding space, we computed the mean Euclidean distance between all pairs of ROIs belonging to the same functional network, for each condition and participant. These distances were then averaged separately across participants with the highest (top 5) and lowest (bottom 5) MEQ scores. Across all four conditions, within-network distances were lower in high-MEQ participants than in low-MEQ participants, as reflected by the heatmap diagonals in Fig. Extended Data 11. This indicates that ROIs within the same network cluster more compactly in the learned embedding space under stronger subjective effects. The consistency of this pattern across rest, meditation, music, and movie confirms that psilocybin-enhanced within-network cohesion is not condition-specific but reflects a broad reorganisation that scales with subjective intensity. This pattern is reported descriptively as convergent support for the formally tested CEBRA findings (Fig. Extended Data 7c,d), where silhouette scores confirmed statistically significant differences in clustering quality between high- and low-MEQ groups across all four conditions.

These low-dimensional embeddings do not measure anatomy or causal directionality. Instead, they reflect how cortical activity reorganises over time in response to psilocybin-dependent subjective effects and context, and reveal behaviourally relevant structure in the temporal organisation of whole-brain activity. The biological inference is therefore about the organisation of distributed network dynamics under psilocybin linked to subjective experience.

### Conventional and ML metric complementarity

Our analytic strategy combined conventional methods (e.g., SD, GFC, parcel FC, modularity, DCM), which quantify amplitude fluctuations, global integration–segregation balance, and directed interactions. These approaches typically characterise macrodose psychedelic states as exhibiting increased signal variance and reduced modular organisation (Carhart-Harris et al., 2016; Daws et al., 2022; Tagliazucchi et al., 2014). In parallel, we applied ML-based embeddings (CEBRA, TAVRNN) that recover latent temporal and regional structure within these high-variance dynamics, and that scale meaningfully with subjective effects.

Whereas traditional metrics primarily index amplitude, variance, and network coupling, ML embeddings capture the high-dimensional temporal geometry of brain state trajectories. These approaches thus probe complementary features of brain activity. Accordingly, differences in apparent similarity—such as high similarity between eyes-closed scans (rest, meditation, music) in traditional metrics versus their condition-separability in ML embeddings, may be expected rather than contradictory.

This complementarity extends to brain–behaviour prediction: per-subject functional modularity under psilocybin was not associated with MEQ30 mean or next-day mindset change in any condition, whereas CEBRA classification accuracy predicted both outcomes.

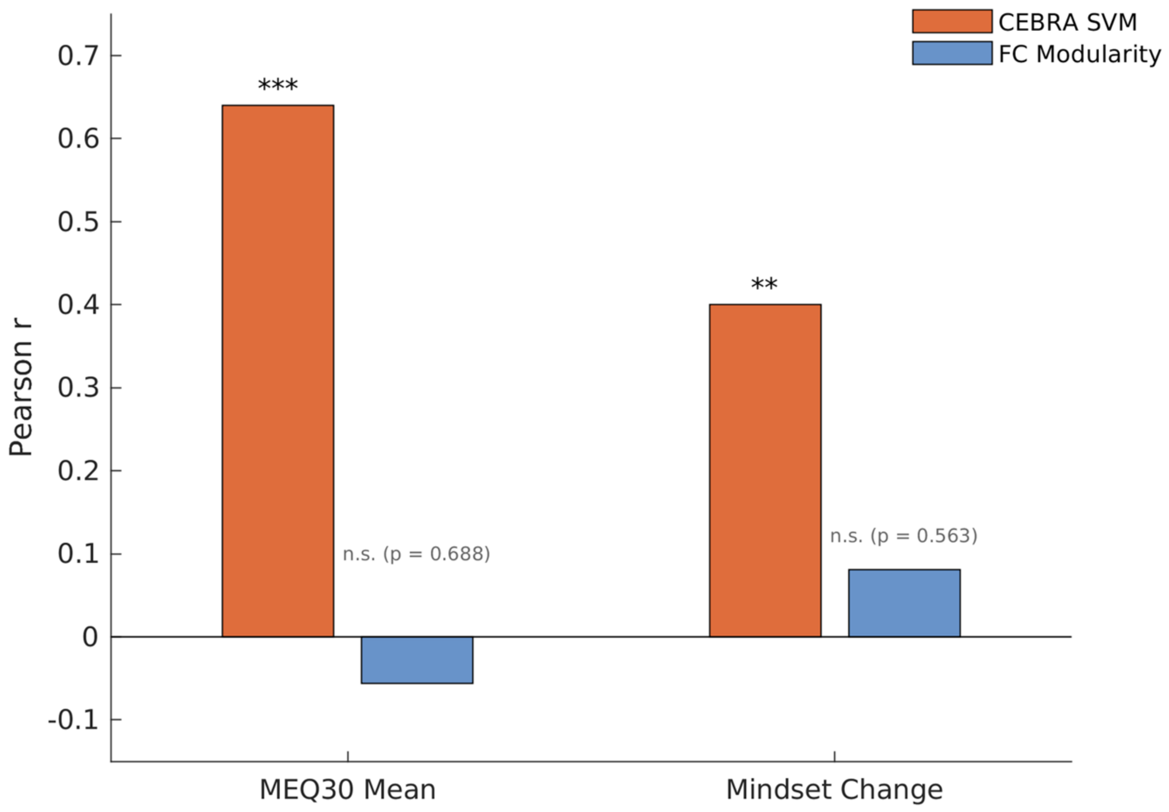


**Comparison of CEBRA SVM and functional modularity correlations with behavioural outcomes.** Functional modularity values were averaged across all four conditions to match the CEBRA analysis, which concatenates all conditions. CEBRA SVM classification accuracy correlated with MEQ30 mean ($r = 0.64$, $p < 10^{-6}$) and mindset change ($r = 0.40$, $p < 0.01$), whereas averaged functional modularity did not (n.s.; exact p values shown; two-tailed Pearson; $n = 53–54$). Per-condition scatter plots are shown below.

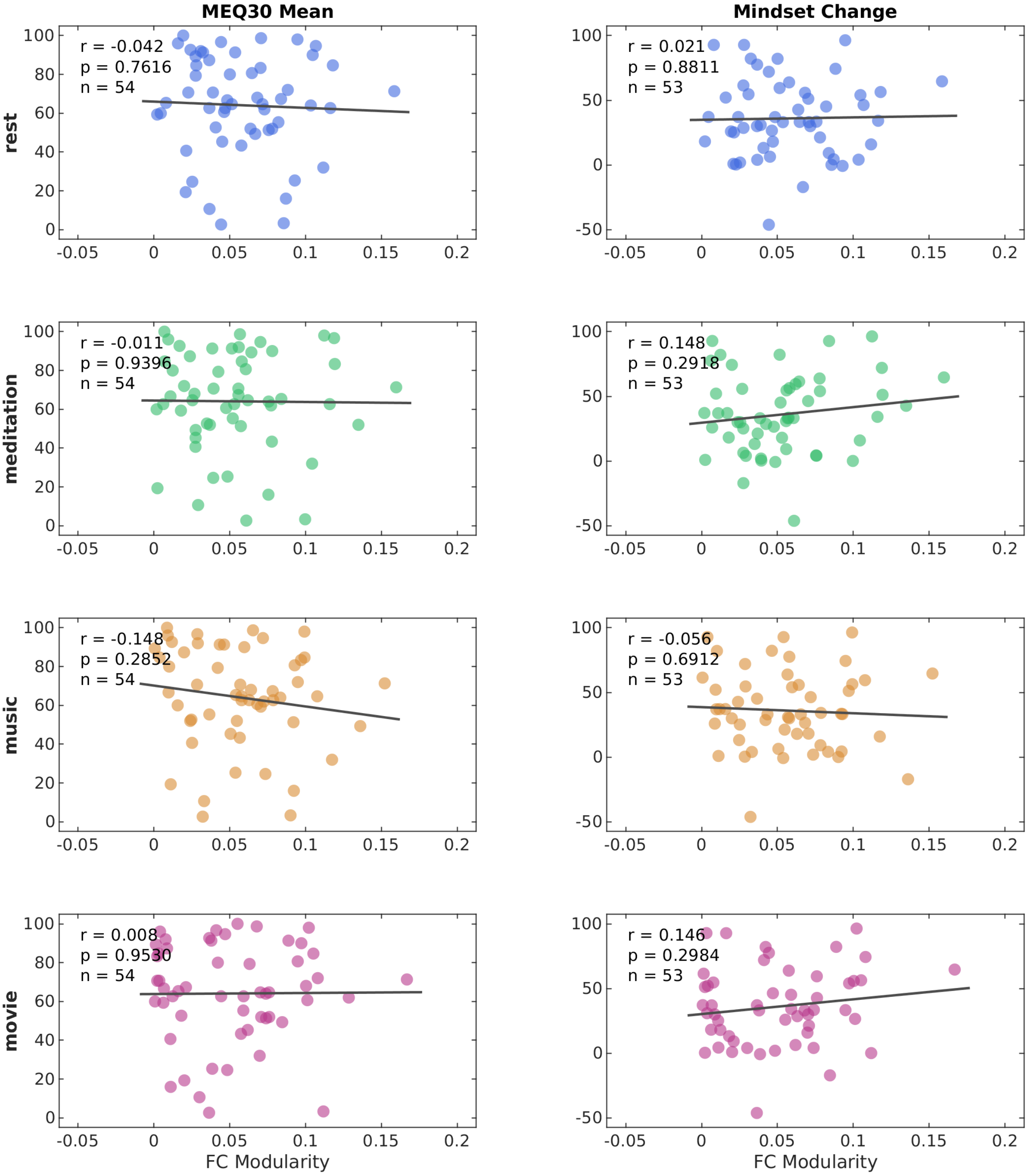


**Per-condition scatter plots of functional modularity vs behavioural outcomes.** Per-subject functional modularity (psilocybin session) plotted against MEQ30 mean (left column, n = 54) and next-day mindset change (right column, n = 53) for each condition (rows: rest, meditation, music, movie). Pearson r, uncorrected p, and n are annotated in each panel. Black lines indicate least-squares regression fits. No correlation was significant (two-tailed Pearson correlations, Bonferroni-corrected across 8 tests, α = 0.0063; confirmed with Spearman rank correlations).

### Spectral Dynamic Causal Modelling

Dynamic causal modelling (DCM) is Bayesian framework that infers the directed (causal) connectivity among the neuronal systems – referred to as effective connectivity. We recently proposed a new DCM for resting state fMRI – based upon a deterministic model that generates predicted cross spectra – referred to as spectral DCM. In order to model resting state activity – in the absence of external stimuli – we will have to add a stochastic component, i.e. neural fluctuations, to the classical DCM based on ordinary differential equations. Mathematically, we can express the formulation of the stochastic generative model using a set of two equations. First is the neuronal state equation, namely

$$\dot{x}(t) = f(x(t), u(t), \theta) + v(t), \tag{S1}$$

and second is the observation equation, which is a static nonlinear mapping from the hidden physiological states in (1) to the observed BOLD activity and is written as:

$$y(t) = h(x(t), \varphi) + e(t), \tag{S2}$$

where $\dot{x}(\mathrm{t})$ is the rate of change of the neuronal states $x(t)$, $\theta$ are unknown parameters (i.e. the effective connectivity) and $v(t)$ (resp. $e(t)$) is the stochastic process – called the state noise (resp. the measurement or observation noise) – modelling the random neuronal fluctuations that drive the resting state activity. In the observation equations, $\varphi$ are the unknown parameters of the (haemodynamic) observation function and $u(t)$ represents any exogenous (or experimental) inputs that drive the hidden states – that are usually absent in resting state designs (Friston et al., 2014). Spectral DCM furnishes a constrained inversion of the stochastic model by parameterising the neuronal fluctuations $v(t)$. Spectral DCM simplifies the generative model by replacing the original timeseries with their second-order statistics (i.e., cross spectra). This means, instead of estimating time varying hidden states, we are estimating their covariance which is time invariant. Then we simply need to estimate the covariance of the random fluctuations; where a scale free (power law) form for the state noise (resp. observation noise) is used – motivated from previous work on neuronal activity (Beggs & Plenz, 2003; Shin & Kim, 2006; Stam & de Bruin, 2004) – as follows:

$$g_v(\omega, \theta) = \alpha_v \omega^{-\beta_v}$$

$$g_e(\omega, \theta) = \alpha_e \omega^{-\beta_e} \tag{S3}$$

Here, $\{\alpha, \beta\} \subset \theta$ are the parameters controlling the amplitudes and exponents of the spectral density of the neural fluctuations. The parameterisation of endogenous fluctuations means that the states are no longer probabilistic; hence the inversion scheme is significantly simpler, requiring estimation of only the parameters (and hyperparameters) of the model.

We used standard Bayesian model inversion to infer the parameters of the model in (1), (2) and (3), from the observed signal $y(t)$. The description of the Bayesian model inversion procedures based on variational Laplace can be found elsewhere for the interested readers (Friston et al., 2007; Friston et al., 2003; A. Razi & K. Friston, 2016).

### Parametric Empirical Bayes

Empirical Bayes refers to the Bayesian inversion or fitting of hierarchical models. In hierarchical models, constraints on the posterior density over model parameters at any given level are provided by the level above. These constraints are called empirical priors because they are informed by empirical data. We recently introduced a second-level or between-subjects model over parameters, which represents how individual (within-subject) connections derive from the subjects' group membership (Friston et al., 2016) – based on parametric empirical Bayes (PEB). This approach calls on Bayesian Model Reduction (BMR) to finesse the inversion of multiple models of a single dataset or a single (hierarchical) model of multiple datasets. BMR allows one to compute posterior densities over model parameters, under new prior densities, without explicitly inverting the model again. For example, one can invert a DCM for each subject in a group and then evaluate the posterior density over group effects, using the posterior densities over parameters from the single subject inversion. This may improve subject-specific parameter estimates, by using group-level estimates to rescue individual DCM from local optima. Mathematically, for DCM studies with *N* subjects and *M* parameters per DCM, we have a hierarchical model, where the responses of the *i*-th subject and the distribution of the parameters over subjects can be modeled as:

$$y_i = \Gamma_i^{(1)}(\theta^{(1)}) + \varepsilon_i^{(1)} \qquad \text{(S4)}$$

$$\theta^{(1)} = \Gamma^{(2)}\big(\theta^{(2)}\big) + \varepsilon^{(2)}$$

$$\theta^{(2)} = \eta + \varepsilon^{(3)}$$

where, $y_i$ is the BOLD time series from *i-th* subject *and* $\Gamma_i^{(1)}$ is a nonlinear mapping from the parameters of a model to the predicted response $y$ for e.g. as shown in Eq. S1 above. $\varepsilon_i^{(1)}$ is independent and identically distributed (i.i.d.) observation noise (equivalent to $e(t)$ in Eq. S2). In this hierarchical form, *empirical priors* encoding second (between-subject) level effects place constraints on subject-specific parameters. The second level would be a linear model where the random effects are parameterised in terms of their precision:

$$\Gamma^{(2)}\big(\theta^{(2)}\big) = (X \otimes W)\beta$$

where, $\beta \subset \theta$ are group means or effects encoded by a design matrix with between $X$ and within-subject $W$ parts. The between-subject part encodes differences among subjects or covariates such as age, while the within-subject part specifies mixtures of parameters that show random effects. We assume that the first column of the design matrix is a constant term, modelling group means and subsequent columns encode group differences or covariates such as age.

### Self-Connections

While effective connectivity matrices are traditionally plotted with log-scaled diagonal elements, here the diagonal elements are transformed via an exponential function (-1/2 * exp(x)) that reverses the log-scaling. This ensures that all the connections (diagonals and offdiagonals) are presented using the same units (Hz), and facilitates the interpretation of changes in self-connection values: a positive change means that a region becomes more responsive to inputs; a negative change means that it becomes less responsive to inputs.

---

### Supplement to subsection Context-dependent Hippocampal-cortical Effective Connectivity Modulation under Psilocybin

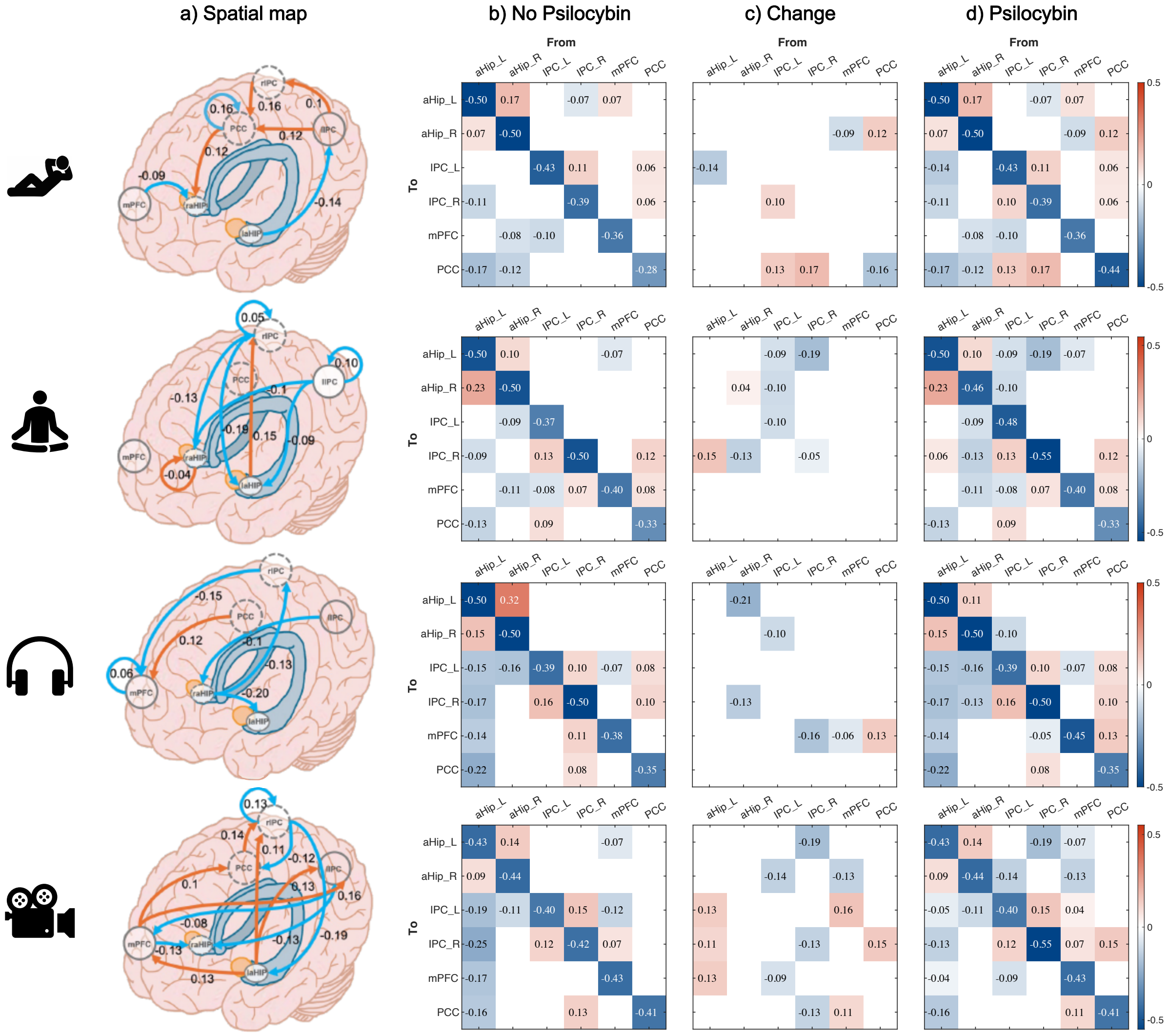


**Model of effective connectivity changes from baseline (no psilocybin) to psilocybin**. Brain regions: Left anterior hippocampus (aHip_L), right anterior hippocampus (aHip_R), left inferior parietal cortex (IPC_L), right inferior parietal cortex (IPC_R), medial prefrontal cortex (mPFC), and posterior cingulate cortex (PCC). Scan sequences, top to bottom: rest, meditation, music, and movie watching. The model was specified as fully connected, allowing all possible directed interactions between regions. a) Spatial mapping of regions of interest with estimated changes in effective connectivity. Warm colours denote increases, cool colours denote decreases (Hz), based on posterior means averaged across participants using Bayesian model averaging. b) Mean effective connectivity at baseline. For mean effective connectivity, warm colours represent excitation and cool colours represent inhibition. c) Matrix of EC changes from baseline to psilocybin, corresponding to a). d) Mean effective connectivity under psilocybin.} Connection strengths (posterior expectations) are reported in Hz. Self-connections are not log-scaled. Displayed connections have a posterior probability $>0.99$, indicating very strong evidence.

Dynamic causal modelling (DCM) (Friston et al., 2003; Friston et al., 2014; A. Razi & K. J. Friston, 2016) revealed context-dependent changes in effective connectivity from baseline (no-psilocybin) to psilocybin administration between the anterior hippocampus (aHip) and nodes of the core Default Mode Network (DMN).

Connectivity patterns varied significantly depending on experimental conditions, with left inferior parietal cortex (IPC_L) consistently inhibiting the right anterior hippocampus (aHip_R) across tasks but not at rest. During the resting state, excitation from the posterior cingulate cortex (PCC) to the aHip_R and inhibition from the medial prefrontal cortex (mPFC) to the aHip_R were observed, alongside inhibition from the left anterior hippocampus (aHip_L) to the IPC_L. These connections suggest a regulatory mechanism, with the PCC driving hippocampal activations, the mPFC attenuating them, and the aHip_L suppressing parietal engagement, potentially tuning self, evaluative and context-related contributions of these regions' integration with memory. During meditation, bilateral inferior parietal cortex (IPC_L and IPC_R) inhibited bilateral anterior hippocampi (aHip_L and aHip_R), suggesting a regulatory role that may inhibit processes involved in memory retrieval and self-related thinking. Under music conditions, inhibition from the aHip_R to the aHip_L was observed, which may alter interhemispheric coordination of memory representations through asymmetric dampening of hippocampal connectivity. The strongest connectivity changes were elicited during eyes-open movie stimuli, where inhibition from the IPC_L, IPC_R, and mPFC to the hippocampi and intra-cortical excitation predominated, in response to a shift toward externally driven visual processing. Excitation from the aHip_L to the mPFC, IPC_R, and IPC_L during this condition may indicate heightened integration of narrative memory with evaluative processes, potentially reflecting a redistribution of influence in this network from higher-order regions to subcortical-driven memory integration. Overall, eyes-open movie watching produced a more complex pattern of connectivity changes relative to the predominantly decrease-biased patterns observed in eyes-closed states, demonstrating that aHip–DMN connectivity is sensitive to context.

Together, these results demonstrate acute, context-dependent reconfiguration of directed aHip–DMN coupling under psilocybin. The dynamic nature of these connectivity changes highlights how psilocybin supports neural adaptability, providing a temporal window for context-dependent neural reorganisation (Calder & Hasler, 2023). These directed connectivity changes offer mechanistic insights that bridge preclinical and post-psilocybin imaging evidence of hippocampal plasticity in humans with acute mechanisms, illustrating how psychological and sensory context (i.e., set and setting) modulate brain function at the circuit level.

These findings extend preclinical and post-administration imaging evidence, highlighting a potential role for the aHip–DMN circuit in psychedelic-induced plasticity (Shao et al., 2021; Siegel et al., 2024; Vaidya et al., 1997) and suggest that structuring a sensory and cognitive environment can shape neural activity and may be used to guide behavioural outcomes. This avenue of research can advance precision psychiatry, along with the examination of the longitudinal behavioural effects of specific connectivity changes in culturally diverse healthy and clinical populations; however, it currently remains underexplored.

These circuit-level changes also complement broader eyes-open vs eyes-closed patterns across analyses, linking circuit-level dynamics to broader transformations—fMRI (global functional connectivity [GFC] and BOLD signal variability [SD]) and EEG (spectral power and Lempel–Ziv complexity [LZC])—while providing directional estimates that complement condition-specific ML embeddings (Main Fig. 2, Fig. 6).

Rest Effective Connectivity Change

| From | To | Posterior expectation | Credible interval |
|---|---|---|---|
| aHip_L | IPC_L | -0.14 | (-0.216,-0.063) |
| IPC_L | PCC | 0.13 | (0.014,0.238) |
| IPC_L | IPC_R | 0.1 | (0.040,0.160) |
| mPFC | aHip_R | -0.09 | (-0.137,-0.046) |
| PCC | aHip_R | 0.12 | (0.078,0.167) |
| IPC_R | PCC | 0.17 | (0.045,0.286) |

Meditation Effective Connectivity Change

| From | To | Posterior expectation | Credible interval |
|---|---|---|---|
| aHip_L | IPC_R | 0.15 | (0.058,0.251) |
| IPC_L | aHip_L | -0.09 | (-0.147,-0.037) |
| IPC_L | aHip_R | -0.1 | (-0.189,-0.017) |
| aHip_R | IPC_R | -0.13 | (-0.201,-0.065) |
| IPC_R | aHip_L | -0.19 | (-0.245,-0.135) |

Music Effective Connectivity Change

| From | To | Posterior expectation | Credible interval |
|---|---|---|---|
| IPC_L | aHip_R | -0.1 | (-0.154,-0.045) |
| PCC | mPFC | 0.13 | (0.080,0.175) |
| aHip_R | aHip_L | -0.21 | (-0.311,-0.100) |
| aHip_R | IPC_R | -0.13 | (-0.203,-0.064) |
| IPC_R | mPFC | -0.16 | (-0.235,-0.082) |

Movie Viewing Effective Connectivity Change

| From | To | Posterior expectation | Credible interval |
|---|---|---|---|
| aHip_L | IPC_L | 0.14 | (0.020,0.249) |
| aHip_L | mPFC | 0.13 | (0.032,0.235) |
| aHip_L | IPC_R | 0.11 | (0.021,0.203) |
| IPC_L | mPFC | -0.09 | (-0.150,-0.026) |
| IPC_L | aHip_R | -0.14 | (-0.205,-0.066) |
| mPFC | IPC_L | 0.16 | (0.086,0.239) |
| mPFC | PCC | 0.11 | (0.034,0.178) |
| mPFC | aHip_R | -0.13 | (-0.189,-0.077) |
| PCC | IPC_R | 0.15 | (0.055,0.238) |
| IPC_R | aHip_L | -0.19 | (-0.259,-0.124) |

| | | | |
|---|---|---|---|
| IPC_R | PCC | -0.13 | (-0.235,-0.017) |

Tables of effective connectivity changes across different scan conditions (rest, meditation, music, and movie watching) for between region connections. Brain regions include the left anterior hippocampus (aHip_L), right anterior hippocampus (aHip_R), left inferior parietal cortex (IPC_L), right inferior parietal cortex (IPC_R), medial prefrontal cortex (mPFC), and posterior cingulate cortex (PCC). The model was specified as fully connected, allowing all possible directed interactions between regions. Effect sizes (posterior expectations) are presented in Hz. Self-connections are not log-scaled. These estimates represent group-level posterior means derived via Bayesian model averaging, highlighting context-dependent modulation of connectivity patterns. Results demonstrate differential changes in aHipp-DMN connectivity across conditions, reinforcing the influence of context on psilocybin-induced network dynamics. Connections with a posterior probability >0.99 are reported, indicating very strong evidence.

Note on model space

Multiple valid operationalisations of the DMN exist (e.g., PCC-centric, precuneus-centric, or combined posterior-midline schemes) (Andrews-Hanna et al., 2010; Thomas Yeo et al., 2011; Utevsky et al., 2014). We selected the posterior cingulate cortex (PCC) as the posterior-midline DMN node to align with our focus on changes to self-related cognition across sensory and cognitive contexts. Although the PCC and precuneus are cytoarchitecturally distinct regions, our DCM analysis models the PCC and precuneus as separate nodes defined by stereotactic coordinates (see Methods); our use of "PCC" therefore refers specifically to the posterior cingulate node centred at the coordinates specified, not to the broader posterior midline cortex. The PCC is implicated in psychedelic neuroimaging (Carhart-Harris et al., 2012; Muthukumaraswamy et al., 2013; Smigielski et al., 2019) and this choice is in line with our prior work, including psychedelic and DCM/DMN studies (Stoliker et al., 2023, 2024) (Preller et al., 2019; Zhou et al., 2018). Given causal evidence implicating the anterior precuneus in bodily-self processing (Lyu et al., 2023), we flag an expanded model variant including an explicit precuneus node as a future, hypothesis-driven extension for studies of the bodily sense of self under psychedelics.

*References*


Andrews-Hanna, J. R., Reidler, J. S., Sepulcre, J., Poulin, R., & Buckner, R. L. (2010). Functional-anatomic fractionation of the brain's default network. *Neuron*, *65*(4), 550–562. https://doi.org/10.1016/j.neuron.2010.02.005

Beggs, J. M., & Plenz, D. (2003). Neuronal avalanches in neocortical circuits. *J Neurosci*, *23*(35), 11167–11177.

Bogenschutz, M. P., Ross, S., Bhatt, S., Baron, T., Forcehimes, A. A., Laska, E., Mennenga, S. E., O'Donnell, K., Owens, L. T., Podrebarac, S., Rotrosen, J., Tonigan, J. S., & Worth, L. (2022). Percentage of Heavy Drinking Days Following Psilocybin-Assisted Psychotherapy vs Placebo in the Treatment of Adult Patients With Alcohol Use Disorder: A Randomized Clinical Trial. *JAMA Psychiatry*, *79*(10), 953–962. https://doi.org/10.1001/jamapsychiatry.2022.2096

Brown, R. T., Nicholas, C. R., Cozzi, N. V., Gassman, M. C., Cooper, K. M., Muller, D., Thomas, C. D., Hetzel, S. J., Henriquez, K. M., Ribaudo, A. S., & Hutson, P. R. (2017). Pharmacokinetics of Escalating Doses of Oral Psilocybin in Healthy Adults. *Clin Pharmacokinet*, *56*(12), 1543–1554. https://doi.org/10.1007/s40262-017-0540-6

Calder, A. E., & Hasler, G. (2023). Towards an understanding of psychedelic-induced neuroplasticity. *Neuropsychopharmacology*, *48*(1), 104–112. https://doi.org/10.1038/s41386-022-01389-z

Carhart-Harris, R. L., Erritzoe, D., Williams, T., Stone, J. M., Reed, L. J., Colasanti, A., Tyacke, R. J., Leech, R., Malizia, A. L., Murphy, K., Hobden, P., Evans, J., Feilding, A., Wise, R. G., & Nutt, D. J. (2012). Neural correlates of the psychedelic state as determined by fMRI studies with psilocybin. *Proceedings of the National Academy of Sciences of the United States of America*, *109*(6), 2138–2143.

Carhart-Harris, R. L., Muthukumaraswamy, S., Roseman, L., Kaelen, M., Droog, W., Murphy, K., Tagliazucchi, E., Schenberg, E. E., Nest, T., Orban, C., Leech, R., Williams, L. T., Williams, T. M., Bolstridge, M., Sessa, B., McGonigle, J., Sereno, M. I., Nichols, D., Hellyer, P. J.,…Nutt, D. J. (2016). Neural correlates of the LSD experience revealed by multimodal neuroimaging. *Proceedings of the National Academy of Sciences of the United States of America*, *113*(17), 4853–4858.

Daws, R. E., Timmermann, C., Giribaldi, B., Sexton, J. D., Wall, M. B., Erritzoe, D., Roseman, L., Nutt, D., & Carhart-Harris, R. (2022). Increased global integration in the brain after psilocybin therapy for depression. *Nature Medicine*, *28*(4), 844–851. https://doi.org/10.1038/s41591-022-01744-z

Friston, K., Mattout, J., Trujillo-Barreto, N., Ashburner, J., & Penny, W. (2007). Variational free energy and the Laplace approximation. *Neuroimage*, *34*(1), 220–234. https://doi.org/10.1016/j.neuroimage.2006.08.035

Friston, K. J., Harrison, L., & Penny, W. (2003). Dynamic causal modelling. *NeuroImage*, *19*(4), 1273–1302. https://doi.org/10.1016/s1053-8119(03)00202-7

Friston, K. J., Kahan, J., Biswal, B., & Razi, A. (2014). A DCM for resting state fMRI. *NeuroImage*, *94*, 396–407. https://doi.org/https://doi.org/10.1016/j.neuroimage.2013.12.009

Friston, K. J., Litvak, V., Oswal, A., Razi, A., Stephan, K. E., van Wijk, B. C., Ziegler, G., & Zeidman, P. (2016). Bayesian model reduction and empirical Bayes for group (DCM) studies. *Neuroimage*, *128*, 413–431. https://doi.org/10.1016/j.neuroimage.2015.11.015

Gasser, P., Holstein, D., Michel, Y., Doblin, R., Yazar-Klosinski, B., Passie, T., & Brenneisen, R. (2014). Safety and efficacy of lysergic acid diethylamide-assisted psychotherapy for anxiety associated with life-threatening diseases. *J Nerv Ment Dis*, *202*(7), 513–520. https://doi.org/10.1097/nmd.0000000000000113

Hasler, F., Grimberg, U., Benz, M. A., Huber, T., & Vollenweider, F. X. (2004). Acute psychological and physiological effects of psilocybin in healthy humans: a double-blind, placebo-controlled dose-effect study. *Psychopharmacology (Berl)*, *172*(2), 145–156. https://doi.org/10.1007/s00213-003-1640-6

Hirschfeld, T., Prugger, J., Majić, T., & Schmidt, T. T. (2023). Dose-response relationships of LSD-induced subjective experiences in humans. *Neuropsychopharmacology*, *48*(11), 1602–1611. https://doi.org/10.1038/s41386-023-01588-2

Hirschfeld, T., & Schmidt, T. T. (2021). Dose–response relationships of psilocybin-induced subjective experiences in humans. *Journal of Psychopharmacology*, *35*(4), 384–397. https://doi.org/10.1177/0269881121992676

Holze, F., Becker, A. M., Kolaczynska, K. E., Duthaler, U., & Liechti, M. E. (2022). Pharmacokinetics and pharmacodynamics of oral psilocybin administration in healthy participants. *Clin Pharmacol Ther*. https://doi.org/10.1002/cpt.2821

Holze, F., Gasser, P., Müller, F., Dolder, P. C., & Liechti, M. E. (2023). Lysergic Acid Diethylamide–Assisted Therapy in Patients With Anxiety With and Without a Life-Threatening Illness: A Randomized, Double-Blind, Placebo-Controlled Phase II Study. *Biological Psychiatry*, *93*(3), 215–223. https://doi.org/10.1016/j.biopsych.2022.08.025

Ley, L., Holze, F., Arikci, D., Becker, A. M., Straumann, I., Klaiber, A., Coviello, F., Dierbach, S., Thomann, J., Duthaler, U., Luethi, D., Varghese, N., Eckert, A., & Liechti, M. E. (2023). Comparative acute effects of mescaline, lysergic acid diethylamide, and psilocybin in a randomized, double-blind, placebo-controlled cross-over study in healthy participants. *Neuropsychopharmacology*, *48*(11), 1659–1667. https://doi.org/10.1038/s41386-023-01607-2

Lyu, D., Stieger, J. R., Xin, C., Ma, E., Lusk, Z., Aparicio, M. K., Werbaneth, K., Perry, C. M., Deisseroth, K., Buch, V., & Parvizi, J. (2023). Causal evidence for the processing of bodily self in the anterior precuneus. *Neuron*, *111*(16), 2502–2512.e2504. https://doi.org/10.1016/j.neuron.2023.05.013

Madsen, M. K., Burmester, D., Stenbaek, D. S., Kristiansen, S., Dyssegaard, A., Lehel, S., Linnet, K., Johansen, S. S., Svarer, C., Ozenne, B., Erritzoe, D., Fisher, P. M., & Knudsen, G. M. (2019). Psilocybin occupancy of brain serotonin 2A receptors correlates with psilocin levels and subjective experience: a 11C Cimbi-36 PET study in humans. *European Neuropsychopharmacology*, *29*, S304–S305. https://doi.org/10.1016/j.euroneuro.2018.11.474

Madsen, M. K., Fisher, P. M., Burmester, D., Dyssegaard, A., Stenbæk, D. S., Kristiansen, S., Johansen, S. S., Lehel, S., Linnet, K., Svarer, C., Erritzoe, D., Ozenne, B., & Knudsen, G. M. (2019). Psychedelic effects of psilocybin correlate with serotonin 2A receptor occupancy and plasma psilocin levels. *Neuropsychopharmacology*, *44*(7), 1328. https://doi.org/10.1038/s41386-019-0324-9

Madsen, M. K., Stenbæk, D. S., Arvidsson, A., Armand, S., Marstrand-Joergensen, M. R., Johansen, S. S., Linnet, K., Ozenne, B., Knudsen, G. M., & Fisher, P. M. (2021). Psilocybin-induced changes in brain network integrity and segregation correlate with plasma psilocin level and psychedelic experience. *European Neuropsychopharmacology*, *50*, 121–132. https://doi.org/https://doi.org/10.1016/j.euroneuro.2021.06.001

Muthukumaraswamy, S. D., Carhart-Harris, R. L., Moran, R. J., Brookes, M. J., Williams, T. M., Errtizoe, D., Sessa, B., Papadopoulos, A., Bolstridge, M., Singh, K. D., Feilding, A., Friston, K. J., & Nutt, D. J. (2013). Broadband Cortical Desynchronization Underlies the Human Psychedelic State. *The Journal of Neuroscience*, *33*(38), 15171–15183. https://doi.org/10.1523/jneurosci.2063-13.2013

Muthukumaraswamy, S. D., Forsyth, A., & Lumley, T. (2021). Blinding and expectancy confounds in psychedelic randomized controlled trials. *Expert Review of Clinical Pharmacology*, *14*(9), 1133–1152. https://doi.org/10.1080/17512433.2021.1933434

Olsen, A. S., Lykkebo-Valløe, A., Ozenne, B., Madsen, M. K., Stenbæk, D. S., Armand, S., Mørup, M., Ganz, M., Knudsen, G. M., & Fisher, P. M. (2022). Psilocybin modulation of time-varying functional connectivity is associated with plasma psilocin and subjective effects. *NeuroImage*, *264*, 119716. https://doi.org/10.1016/j.neuroimage.2022.119716

Otto, M. E., van der Heijden, K. V., Schoones, J. W., van Esdonk, M. J., Borghans, L., Jacobs, G. E., & van Hasselt, J. G. C. (2025). Clinical Pharmacokinetics of Psilocin After Psilocybin Administration: A Systematic Review and Post-Hoc Analysis. *Clin Pharmacokinet*, *64*(1), 53–66. https://doi.org/10.1007/s40262-024-01454-4

Preller, K. H., Burt, J. B., Ji, J. L., Schleifer, C. H., Adkinson, B. D., Stämpfli, P., Seifritz, E., Repovs, G., Krystal, J. H., Murray, J. D., Vollenweider, F. X., & Anticevic, A. (2018). Changes in global and thalamic brain connectivity in LSD-induced altered states of consciousness are attributable to the 5-HT2A receptor. *eLife*, *7*, e35082. https://doi.org/10.7554/eLife.35082

Preller, K. H., Duerler, P., Burt, J. B., Ji, J. L., Adkinson, B., Stämpfli, P., Seifritz, E., Repovš, G., Krystal, J. H., Murray, J. D., Anticevic, A., & Vollenweider, F. X. (2020). Psilocybin Induces Time-Dependent Changes in Global Functional Connectivity. *Biological Psychiatry*, *88*(2), 197–207. https://doi.org/https://doi.org/10.1016/j.biopsych.2019.12.027

Preller, K. H., Razi, A., Zeidman, P., Stämpfli, P., Friston, K. J., & Vollenweider, F. X. (2019). Effective connectivity changes in LSD-induced altered states of consciousness in humans. *Proceedings of the National Academy of Sciences*, *116*(7), 2743–2748. https://doi.org/10.1073/pnas.1815129116

Razi, A., & Friston, K. (2016). The Connected Brain: Causality, models, and intrinsic dynamics. *IEEE Signal Processing Magazine*, *33*(3), 14–35.

Razi, A., & Friston, K. J. (2016). The Connected Brain: Causality, models, and intrinsic dynamics. *IEEE Signal Processing Magazine*, *33*(3), 14–35. https://doi.org/10.1109/MSP.2015.2482121

Schartner, M. M., Carhart-Harris, R. L., Barrett, A. B., Seth, A. K., & Muthukumaraswamy, S. D. (2017). Increased spontaneous MEG signal diversity for psychoactive doses of ketamine, LSD and psilocybin. *Scientific Reports*, *7*(1), 46421. https://doi.org/10.1038/srep46421

Shao, L. X., Liao, C., Gregg, I., Davoudian, P. A., Savalia, N. K., Delagarza, K., & Kwan, A. C. (2021). Psilocybin induces rapid and persistent growth of dendritic spines in frontal cortex in vivo. *Neuron*, *109*(16), 2535–2544.e2534. https://doi.org/10.1016/j.neuron.2021.06.008

Shin, C. W., & Kim, S. (2006). Self-organized criticality and scale-free properties in emergent functional neural networks. *Phys Rev E Stat Nonlin Soft Matter Phys*, *74*(4 Pt 2), 045101. https://doi.org/10.1103/PhysRevE.74.045101

Siegel, J. S., Subramanian, S., Perry, D., Kay, B. P., Gordon, E. M., Laumann, T. O., Reneau, T. R., Metcalf, N. V., Chacko, R. V., Gratton, C., Horan, C., Krimmel, S. R., Shimony, J. S., Schweiger, J. A., Wong, D. F., Bender, D. A., Scheidter, K. M., Whiting, F. I., Padawer-Curry, J. A.,…Dosenbach, N. U. F. (2024). Psilocybin desynchronizes the human brain. *Nature*. https://doi.org/10.1038/s41586-024-07624-5

Smigielski, L., Scheidegger, M., Kometer, M., & Vollenweider, F. X. (2019). Psilocybin-assisted mindfulness training modulates self-consciousness and brain default mode network connectivity with lasting effects. *NeuroImage*, *196*, 207–215. https://doi.org/10.1016/j.neuroimage.2019.04.009

Stam, C. J., & de Bruin, E. A. (2004). Scale-free dynamics of global functional connectivity in the human brain. *Hum Brain Mapp*, *22*(2), 97–109. https://doi.org/10.1002/hbm.20016

Stoliker, D., Novelli, L., Vollenweider, F. X., Egan, G. F., Preller, K. H., & Razi, A. (2023). Effective Connectivity of Functionally Anticorrelated Networks Under Lysergic Acid Diethylamide. *Biological Psychiatry*, *93*(3), 224–232. https://doi.org/10.1016/j.biopsych.2022.07.013

Stoliker, D., Novelli, L., Vollenweider, F. X., Egan, G. F., Preller, K. H., & Razi, A. (2024). Neural Mechanisms of Resting-State Networks and the Amygdala underlying the Cognitive and Emotional Effects of Psilocybin. *Biological Psychiatry*. https://doi.org/https://doi.org/10.1016/j.biopsych.2024.01.002

Studerus, E., Kometer, M., Hasler, F., & Vollenweider, F. X. (2011). Acute, subacute and long-term subjective effects of psilocybin in healthy humans: a pooled analysis of experimental studies. *Journal of Psychopharmacology*, *25*(11), 1434–1452. https://doi.org/10.1177/0269881110382466

Szigeti, B., & Heifets, B. D. (2024). Expectancy Effects in Psychedelic Trials. *Biological Psychiatry: Cognitive Neuroscience and Neuroimaging*, *9*(5), 512–521. https://doi.org/https://doi.org/10.1016/j.bpsc.2024.02.004

Szigeti, B., Weiss, B., Rosas, F. E., Erritzoe, D., Nutt, D., & Carhart-Harris, R. (2024). Assessing expectancy and suggestibility in a trial of escitalopram v. psilocybin for depression. *Psychological Medicine*, *54*(8), 1717–1724. https://doi.org/10.1017/S0033291723003653

Tagliazucchi, E., Carhart-Harris, R., Leech, R., Nutt, D., & Chialvo, D. R. (2014). Enhanced repertoire of brain dynamical states during the psychedelic experience. *Human Brain Mapping*, *35*(11), 5442–5456. https://doi.org/10.1002/hbm.22562

Thomas Yeo, B. T., Krienen, F. M., Sepulcre, J., Sabuncu, M. R., Lashkari, D., Hollinshead, M., Roffman, J. L., Smoller, J. W., Zöllei, L., Polimeni, J. R., Fischl, B., Liu, H., & Buckner, R. L. (2011). The organization of the human cerebral cortex estimated by intrinsic functional connectivity. *Journal of Neurophysiology*, *106*(3), 1125–1165. https://doi.org/10.1152/jn.00338.2011

Timmermann, C., Roseman, L., Haridas, S., Rosas, F. E., Luan, L., Kettner, H., Martell, J., Erritzoe, D., Tagliazucchi, E., Pallavicini, C., Girn, M., Alamia, A., Leech, R., Nutt, D. J., & Carhart-Harris, R. L. (2023). Human brain effects of DMT assessed via EEG-fMRI. *Proceedings of the National Academy of Sciences*, *120*(13), e2218949120. https://doi.org/10.1073/pnas.2218949120

Utevsky, A. V., Smith, D. V., & Huettel, S. A. (2014). Precuneus is a functional core of the default-mode network. *The Journal of Neuroscience*, *34*(3), 932–940. https://doi.org/10.1523/jneurosci.4227-13.2014

Vaidya, V. A., Marek, G. J., Aghajanian, G. K., & Duman, R. S. (1997). 5-HT <sub>2A</sub> Receptor-Mediated Regulation of Brain-Derived Neurotrophic Factor mRNA in the Hippocampus and the Neocortex. *The Journal of Neuroscience*, *17*(8), 2785–2795. https://doi.org/10.1523/JNEUROSCI.17-08-02785.1997

Zhou, Y., Friston, K. J., Zeidman, P., Chen, J., Li, S., & Razi, A. (2018). The Hierarchical Organization of the Default, Dorsal Attention and Salience Networks in Adolescents

and Young Adults. *Cereb Cortex*, *28*(2), 726–737.
https://doi.org/10.1093/cercor/bhx307
